# Mechanistically Defined Epoxide- and Aziridine-2-carboxamide Electrophiles Enable Stereoselective Covalent RNA Modulation

**DOI:** 10.64898/2026.01.15.699758

**Authors:** Chungen Li, Xueyi Yang, Jingsong Shan, Kyle A. Dickerson, Noah A. Springer, Nikhil C. Munshi, Robert T. Batey, Matthew D. Disney

## Abstract

RNA remains a largely untapped target for covalent small-molecule intervention due to the lack of electrophiles with predictable reactivity and stability in biological settings. Here, a mechanistically defined and tunable class of epoxide- and aziridine-2-carboxamide electrophiles that enable structure-guided covalent targeting of RNA is described. These warheads arise from an unexpected hydrolytic rearrangement of 3-chloropivalamide precursors under physiological conditions and selectively react with guanine N7, with reactivity and stability controlled by substitution pattern, linkage chemistry, and stereochemistry. Application to two distinct RNA targets demonstrates generality: epoxide- and aziridine-based ligands covalently modify pathogenic r(CUG)^exp^ repeat RNA and disrupt RNA–protein interactions *in vitro* and in cells, while structure-guided placement on a flavin scaffold yields stereoselective covalent modulators of the flavin mononucleotide (FMN) riboswitch with validated reaction site and cellular activity. Together, this work establishes epoxide- and aziridine-2-carboxamides as a versatile platform for covalent RNA targeting and provides a general framework for the rational design of stereochemically controlled RNA-reactive small molecules.

## INTRODUCTION

RNA is an attractive target for therapeutic intervention, although its lack of chemical diversity and conformational dynamics are challenges for small-molecule discovery. Increasing attention has been directed toward the development of both noncovalent and covalent modulators of RNA function.^1^ Covalent ligands are known to provide enhanced potency, sustained target engagement, and potentially improve selectivity through the formation of stable covalent bonds with defined nucleotides or residues.^2^ Beyond these classical advantages, covalent modification offers unique opportunities to modulate RNA function directly. Covalent attachment within an mRNA can suppress translation, meanwhile modification of folded regulatory elements—such as riboswitches in 5′ leader sequences—can stabilize specific conformations that alter gene expression.^3^ In addition, covalent small molecules have been used as chemical probes to elucidate RNA structure and function, thereby expanding the range of RNA folds that can be effectively targeted.^4,5, 6^

Progress in this field has been constrained by the limited availability of electrophiles that combine broad applicability with tunable reactivity and stability. A comprehensive framework for the rational design of RNA-directed electrophiles has yet to be established. Several electrophilic classes have been reported, including diazirines,^7^ *N*-(2-chloroethyl)anilines,^5^ acyl imidazoles,^8^ epoxides,^9^ activated sulfonyl groups,^10^ and alkyl bromides.^11^ In a previous study, a structure-guided approach was applied to design a phenylglyoxal-based covalent modulator of the flavin mononucleotide (FMN) riboswitch by targeting unpaired guanines required for ligand recognition.^3^ In separate work, 3-chloropivalamide was identified through high-throughput mass spectrometry screening as a new class of RNA-reactive electrophile (Fig. 1A).^12^

**Figure 1.**
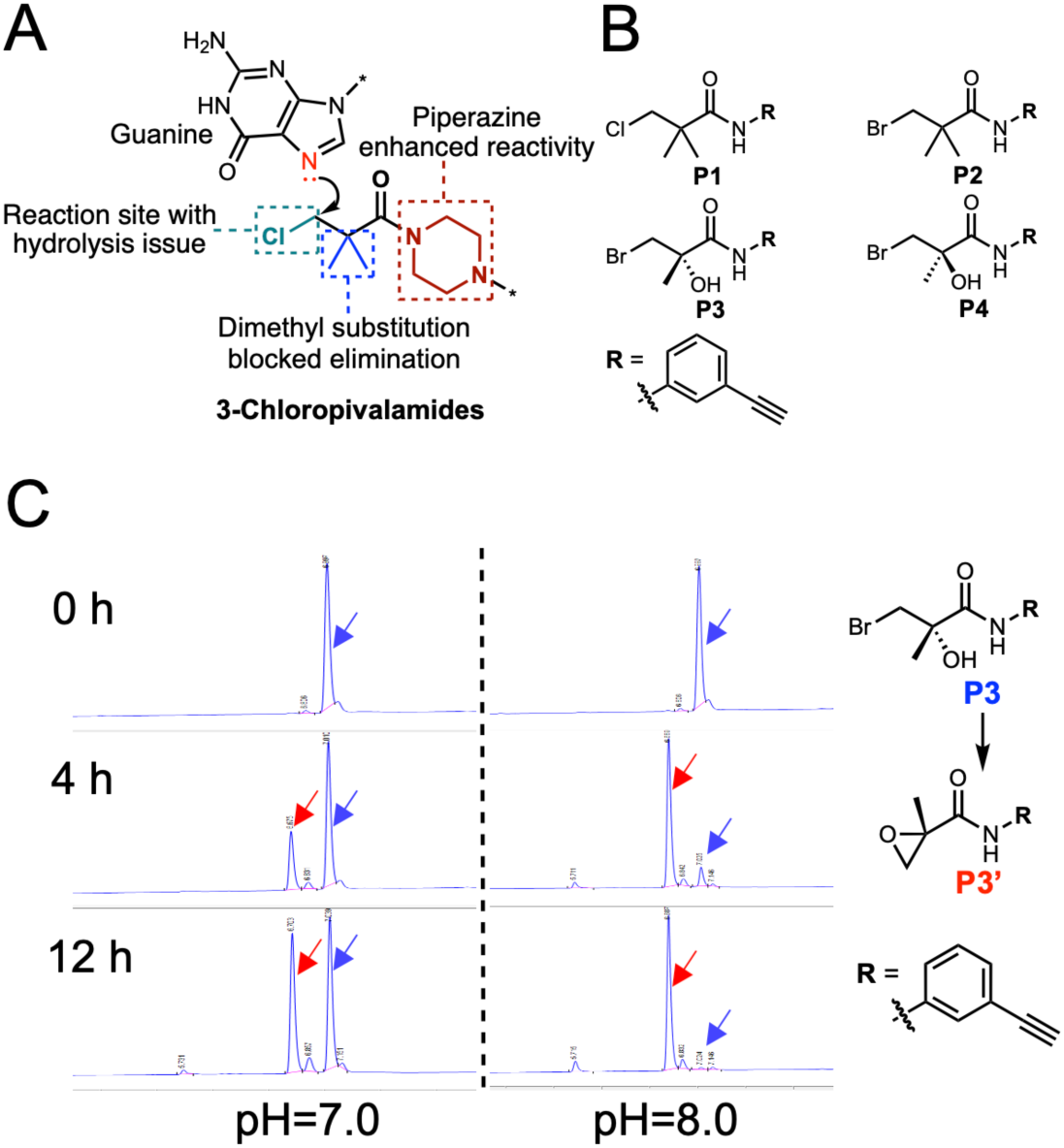
Discovery of epoxide-2-carboxamide electrophiles. A) Chemical structure and key properties of the 3-chloropivalamide electrophile. B) Chemical structures of probe compounds designed based on the 3-chloropivalamides. C) Conversion of probe **P3** to epoxide-2-carboxamide **P3’**. LC-MS traces of **P3** in buffer at pH 7.0 (left) and 8.0 (right). The peaks of **P3** (blue) and **P3’** (red) are indicated with arrows.

The 3-chloropivalamide electrophile was observed to react with the nucleophilic N7 nitrogen of guanine, enabling attack on electrophilic carbon centers *via* an S_N_2 mechanism (Fig. 1A).^12^ This electrophile displayed promising covalent activity and could be readily synthesized by coupling 3-chloro-2,2-dimethylpropanoyl chloride (or its acid) with a variety of amines, providing broad synthetic accessibility. Significant hydrolytic instability, however, was observed, as rapid dechlorination in aqueous buffer yielded unreactive hydroxylated byproducts. Moreover, reactivity depends strongly on the amine substituent, with saturated cyclic amines such as piperidine and piperazine supporting reactivity, whereas aromatic or primary alkyl amines significantly attenuated the reactivity.^12^

In this study, the structure of 3-chloropivalamides was optimized, leading to the identification of an unexpected rearrangement to an epoxide-2-carboxamide warhead under physiological conditions. Through rational design, this epoxide-2-carboxamide enabled the development of covalent small molecules that modify a r(CUG) repeat RNA, a validated structural model of the pathogenic expanded r(CUG) repeat [r(CUG)^exp^] that causes myotonic dystrophy type 1 (DM1), an incurable RNA gain-of-function disease.^13^ These compounds exhibit improved stability and tunable reactivity in biological settings. Structure–activity relationship (SAR) studies further revealed that an aziridine-2-carboxamide analog provides a complementary electrophile with similarly favorable stability and controllable reactivity.

Moreover, applying these warheads in a structure-guided manner enabled the identification of stereochemically defined covalent ligands targeting the flavin mononucleotide (FMN) riboswitch, a regulatory RNA element whose modulation has potential as an antibacterial strategy, yielding validated reaction site and general design principles for the development of selective RNA-reactive small molecules. Collectively, these results establish epoxide- and aziridine-2-carboxamides as a versatile and tunable class of electrophiles for RNA-targeted covalent ligand development.

## RESULTS

### Epoxide-2-carboxamide Is Identified as an RNA-Targeted Covalent Electrophile

A series of alkyne-tagged probe compounds (**P1**–**P4**) was designed based on the 3-chloropivalamide scaffold (Fig. 1B). Attachment of 3-chloropivalamide to a 3-ethynylphenyl motif yielded control probe **P1**, and replacement of the chlorine substituent with a more labile bromine afforded **P2**. The geminal dimethyl substitution of the 3-chloropivalamide scaffold, which suppresses *β*-elimination, was retained (Fig. 1A). The disubstituted scaffold of **P2** was further modified by introduction of a chiral hydroxyl group to generate probes **P3** and **P4**.

The reactivity of probes **P1**–**P4** was first studied toward the r(CUG)^exp^ that cause myotonic dystrophy type 1 (DM1). Here, Cy5-labeled r(CUG)_12_ was evaluated; r(CUG)_12_ is a validated model of r(CUG)^exp^, folding into an array of 1×1 nucleotide U×U internal loops (Fig. S1A).^14^ Covalent modification of the RNA by the probes was assessed by using an established tetramethylrhodamine (TAMRA) labeling assay (Fig. S1A).^5^ At 200 μM, **P1** produced negligible fluorescence labeling of the RNA (Fig. S1B), consistent with previously reported reactivity.^12^ **P2** exhibited only trace TAMRA labeling relative to **P1**. In contrast, **P3** and **P4** produced robust and markedly enhanced fluorescent labeling of r(CUG)_12_ (Fig. S1B), indicating a requirement for the hydroxyl group in covalent modification. The 3-bromo-2-hydroxy-2-methylpropanamide warhead present in **P3** and **P4** has been shown to undergo intermolecular S_N_2 substitution with nucleophiles.^15^ In addition, the hydroxyl group may function as an internal nucleophile, enabling intramolecular displacement of the bromide-bearing carbon to form the corresponding epoxide.^16^

Formation of the epoxide product in reaction buffer was monitored by liquid chromatography–mass spectrometry (LC–MS). Approximately 50% of **P3** was converted to the epoxide-2-carboxamide **P3′** after 12 h of incubation at 37 °C in a pH 7.0 buffer (Fig. 1C). This intramolecular cyclization displayed pronounced pH dependence, with increased conversion observed at higher pH; at pH 8.0, approximately 90% of **P3** was converted to **P3′** within 4 h (Fig. 1C). Consistent with these observations, pH-dependent TAMRA labeling of r(CUG)_12_ was observed for **P3** (Fig. S1C). In contrast, purified epoxide-2-carboxamide **P3′** produced strong and essentially identical fluorescence signals across the tested pH range (Fig. S1C). These results indicate that the covalent reactivity observed for **P3** and **P4** is mediated by formation of an epoxide-2-carboxamide warhead.

### Rational Design of Epoxide-2-carboxamide-based Covalent Ligands for r(CUG)^exp^

Epoxide-2-carboxamide and 3-chloropivalamide warheads were conjugated to a previously validated r(CUG)^exp^-binding compound, **H**,^17^ to generate compounds **H1**–**H2** and **H3**, respectively (Fig. 2A). The only structural differences among **H1**–**H3** were the identity of the covalent warhead (**H1** and **H2** vs. **H3**) and the stereochemistry of the epoxide warhead in **H1** and **H2**. Thus, direct comparisons of the properties of the two warhead classes can be made.

**Figure 2.**
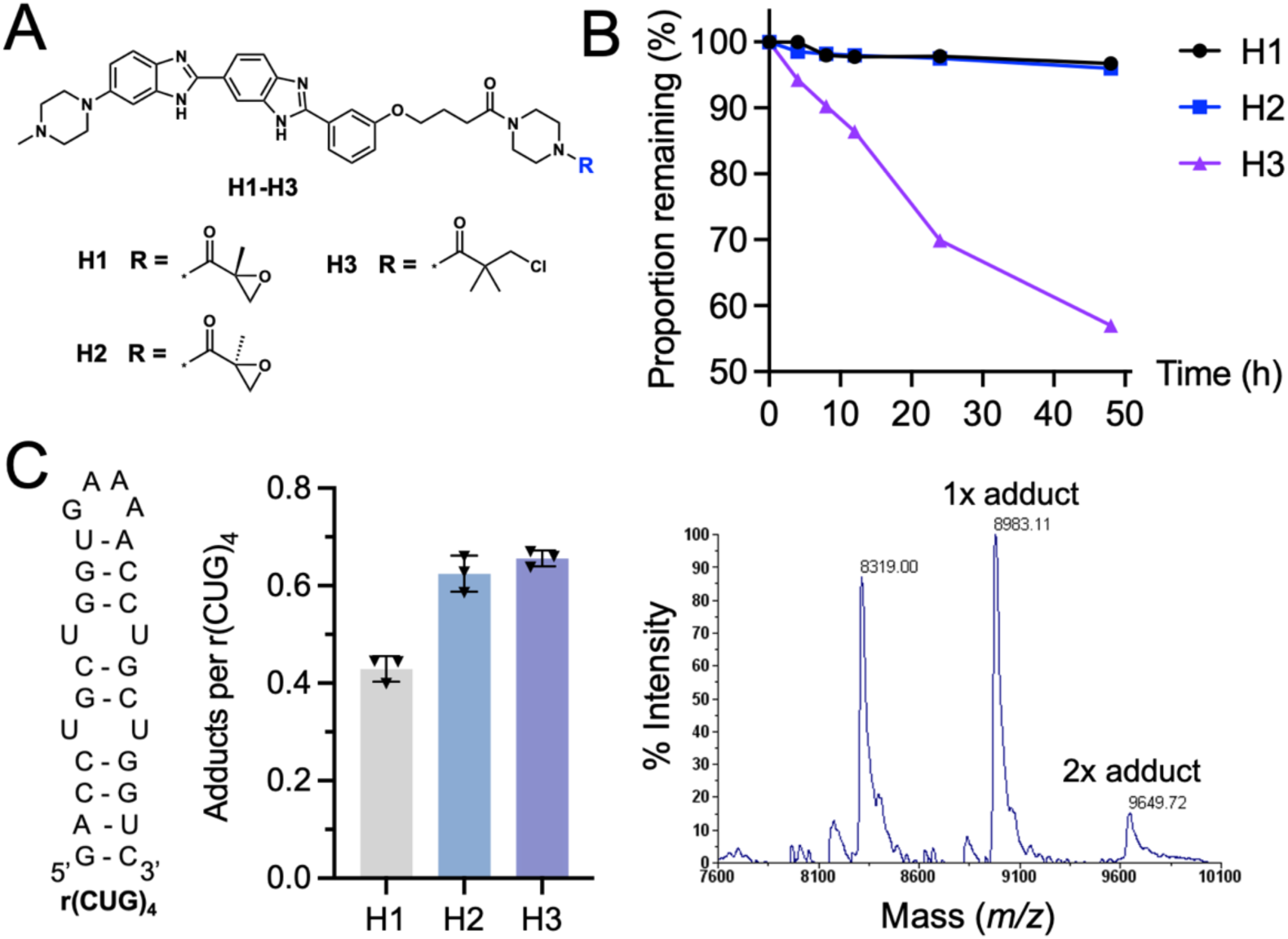
Developing covalent r(CUG)^exp^ ligands based on the epoxide-2-carboxamide warhead. A) Chemical structures of compounds **H1**-**H3**. B) Hydrolytic stabilities of compounds **H1**-**H3** in the r(CUG)_4_ folding buffer (pH 6.5) at 37 °C. C) Left: Secondary structure of r(CUG)_4_. Middle: Covalent modification of r(CUG)_4_ (2 μM) by compounds **H1**-**H3** (200 μM) at 37 °C for 12 h. Quantification was performed by integrating the MALDI-TOF peak areas of unmodified and modified RNA (n = 3). Right: Representative MALDI-TOF mass spectrum of r(CUG)_4_ (2 μM) reacted with **H2** (200 μM). Peaks are labeled with its corresponding mass-to-charge ratio (*m/z*).

The hydrolytic stability of compounds **H1**–**H3** was evaluated in reaction buffer (50 mM HEPES, pH 6.5, 100 mM KCl, and 15 mM MgCl_2_). Compounds **H1** and **H2** exhibited high stability, with more than 95% of each compound remaining after 48 h of incubation at 37 °C (Fig. 2B). No significant side products, including hydrolysis products, were detected. In contrast, the 3-chloropivalamide compound **H3** showed pronounced instability, with only 57% of the parent compound remaining under the same conditions (Fig. 2B). Hydrolysis of the 3-chloropivalamide moiety was identified as the predominant byproduct (Fig. S2A).

Covalent reactivity of compounds **H1**–**H3** toward r(CUG)^exp^ was assessed using an established matrix-assisted laser desorption/ionization time-of-flight mass spectrometry (MALDI-TOF-MS) assay.^12^ To facilitate MALDI system detection, a shorter r(CUG)_4_ was evaluated, which contains two unpaired 1×1 nucleotide U×U internal loops (Fig. 2C). The extent of covalent modification was quantified by integration of peak areas corresponding to modified and unmodified RNA species. After 12 h of incubation at 37 °C, robust covalent modification of r(CUG)_4_ was observed for the epoxide-2-carboxamide compounds **H1** and **H2** (Fig. 2C). Notably, **H1** and **H2** share identical intrinsic electrophilicity, but *S*-enantiomer **H2** (0.62 ± 0.04 adducts per RNA) exhibited higher reactivity than the *R*-enantiomer **H1** (0.43 ± 0.03 adducts per RNA) and displayed covalent efficiency comparable to that of the 3-chloropivalamide analogue **H3** (0.65 ± 0.02 adducts per RNA) at 200 µM (Fig. 2C, Fig. S2B). These results suggest that stereochemistry-dependent warhead orientation may contribute to covalent reactivity. In contrast, compound **P5**, which lacks the r(CUG)^exp^-binding scaffold, showed no detectable modification under the same conditions (Fig. S2C).

### Mechanistic Elucidation of Epoxide-2-carboxamide Electrophile

High-resolution LC–MS analysis was performed on enzymatically digested mononucleotides derived from r(CUG)_4_ modified with compound **H2** to assess the site of covalent modification. An **H2**–guanosine adduct, [A1]^+^, was detected with a molecular weight of 946.4300 (calculated for [A1]^+^, 946.4324), while no adducts corresponding to uridine or cytidine were observed (Fig. 3A). Prominent fragment ions corresponding to the **H2**–guanine adduct, [A2]^+^ (lacks ribose moiety), were observed with a molecular weight of 814.3905 (calculated for [A2]^+^, 814.3901), along with clear half-mass peaks for both the **H2**–guanosine ([A1+H^+^]^2+^) and **H2**–guanine ([A2+H^+^]^2+^) species (Fig. 3A). These data indicate selective modification of the guanine base.

**Figure 3.**
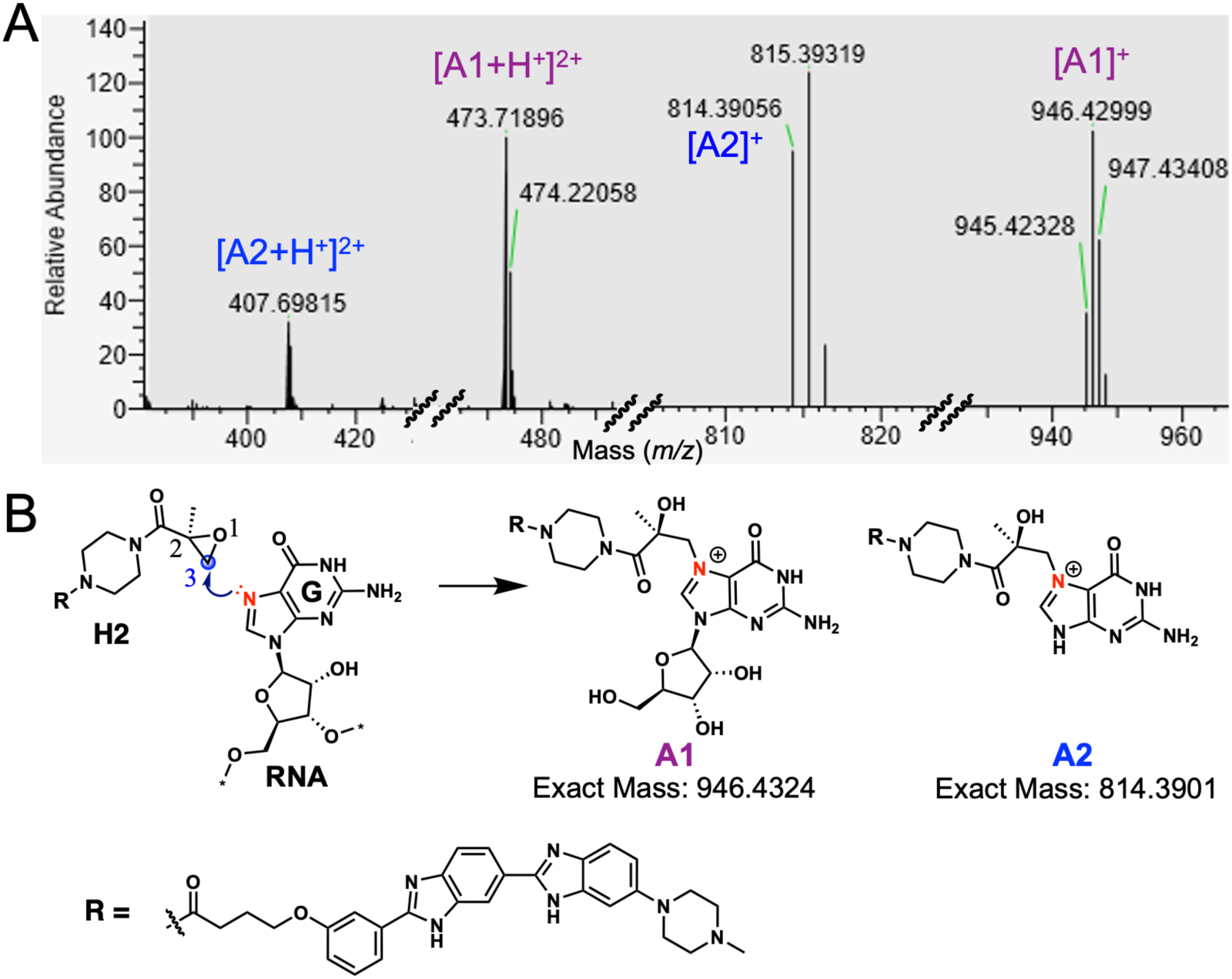
Adduct identification for compound H2 by high-resolution LC–MS. A) High-resolution LC–MS spectrum of enzymatically digested r(CUG)_4_ following **H2** treatment. B) The predicted covalent modification mechanism of epoxide-2-carboxamide warhead. Possible chemical structures of the **H2**-guanosine (A1) and **H2**-guanine (A2) adducts based on the high-resolution LC–MS.

Epoxide-mediated alkylation of DNA has been reported to occur predominantly at the N7 position of guanine.^18^ In RNA, the N7 nitrogen of guanine has been reported to react with a range of electrophilic warheads.^9, 11, 19^ Given that the 2-position of the epoxide-2-carboxamide warhead in **H2** is methyl-substituted, nucleophilic attack by guanine N7 is proposed to occur at the 3-position of the epoxide, leading to ring opening and formation of a primary alcohol adduct (Fig. 3B).

### Structure–Activity Relationships (SAR) of the Epoxide-2-carboxamide Electrophile

A SAR study was conducted to characterize the reactivity and stability of the epoxide-2-carboxamide warhead (Table 1, Fig. S3). Removal of the 2-position methyl substituent (**H4** and **H5**) resulted in increased covalent modification relative to **H1** and **H2** at 200 μM and retained substantial modification at 100 μM. However, **H4** and **H5** exhibited reduced stability, with more than 60% of each compound converted to inactive products after 48 h of incubation at 37 °C. LC–MS analysis identified two predominant degradation pathways: (i) hydrolytic opening of the epoxide ring and (ii) nucleophilic attack by chloride present in the buffer, yielding the corresponding chlorinated products (Fig. S4). In contrast, methyl substitution at the 3-position (**H6**) resulted in high stability but completely abolished covalent reactivity. These observations are consistent with modification occurring at the 3-position of the epoxide, in agreement with the covalent mechanism inferred for **H2** from high-resolution LC–MS analysis (Fig. 3). Methyl substitution at the 2-position of the epoxide-2-carboxamide warhead was associated with enhanced stability relative to the unsubstituted analogues.

**Table 1.** Covalent activity and stability of derivatives.

| Cmpd. | R | Adducts per r(CUG)<sub>4</sub> |  | Stability <sup>a</sup><br>(%) |
| --- | --- | --- | --- | --- |
| | | 100 μM | 200 μM | |
| H1 | | - | 0.43 ± 0.03 | 96.7 |
| H2 | | - | 0.62 ± 0.04 | 96.0 |
| H4 | | 0.51 ± 0.03 | 0.90 ± 0.04 | 35.6 |
| H5 | | 0.39 ± 0.03 | 0.91 ± 0.02 | 37.5 |
| H6 |  | 0 | 0 | 96.4 |
| H7 | | 0.65 ± 0.02 | 0.93 ± 0.06 | - |
| H8 | | 0.76 ± 0.04 | 0.95 ± 0.01 | 31.7 |
| H9 | | - | 0.08 ± 0.02 | - |
| H10 | | - | 0.12 ± 0.02 | 71.8 |
| H11 | | 0.18 ± 0.02 | 0.32 ± 0.03 | - |
| H12 | | 0.35 ± 0.02 | 0.55 ± 0.04 | 98.0 |
| H13 |  | 0 | 0 | - |
| H14 |  | - | - | - |

-, not determined. <sup>a</sup> Stability, the proportion remaining after 48 h incubation in FMN riboswitch folding buffer at 37 °C.

Compounds **H7** and **H8** exhibited increased covalent modification relative to **H4** and **H5** at both 100 and 200 μM, indicating that α-methylation adjacent to the warhead–piperazine linkage influences reactivity. Replacement of the piperazine moiety with an aniline group (**H9** and **H10**) resulted in markedly reduced covalent modification, suggesting a role for the piperazine in facilitating warhead reactivity. Similar reactivity trends have been reported for 3-chloropivalamides^12^ and chloroacetamides.^20^

Aziridine, an analogue of epoxide, has been widely employed as an electrophilic covalent warhead in protein-targeted systems.^21^ Compounds **H11** and **H12** bearing an aziridine-2-carboxamide warhead also exhibited covalent modification of r(CUG)_4_. Compound **H12** displayed high stability, with less than 2% degradation observed after 48 h of incubation at 37 °C. Although lower covalent modification was observed for **H12** relative to the epoxide-2-carboxamide analogue **H5**, substantial stability was retained. Consistent with observations for epoxide-2-carboxamide **H6**, methyl substitution at the 3-position of the aziridine-2-carboxamide warhead (**H13**) eliminated detectable covalent reactivity, indicating a shared mechanistic requirement for the substitution pattern.

Across the compound series, comparison of enantiomeric pairs on r(CUG)_4_ showed that *S*-enantiomers consistently exhibited higher levels of covalent modification than the corresponding *R*-enantiomers at 200 μM. These results support that stereochemistry-dependent warhead orientation contributes to covalent reactivity, suggesting the potential value of chirality in applying epoxide- and aziridine-2-carboxamide warheads to develop stereoselective RNA-targeted covalent ligands.

### Disruption of the r(CUG)^exp^–MBNL1 Complex by H1–H2

The r(CUG)^exp^ sequesters MBNL1 protein into nuclear foci, resulting in loss of MBNL1 function and mis-splicing events associated with myotonic dystrophy type 1 (DM1).^13^ Disruption of the r(CUG)^exp^–MBNL1 complex by compounds **H1**–**H2** was evaluated using a previously developed *in vitro* time-resolved fluorescence resonance energy transfer (TR-FRET) assay.^22^ A biotinylated r(CUG)_12_ and His_6_-tagged MBNL1 were used in this assay. Upon the formation of complex, a FRET signal is generated through the streptavidin-XL665 and an anti-His_6_-terbium-labeled antibody. Consequently, inhibition of the r(CUG)^exp^–MBNL1 interaction by small molecules results in a decrease in the TR-FRET signal.

Compounds were preincubated with biotinylated r(CUG)_12_ at 37 °C for 12 h to allow covalent modification prior to assay initiation. Compound **H2** effectively disrupted the r(CUG)^exp^–MBNL1 complex in a dose-dependent manner over the range of 20-100 µM, with an inhibition of 72.9 ± 2.3% at 100 µM (Fig. S5). **H1** displayed a comparable inhibition at 100 µM but showed significantly reduced activity at 50 µM relative to **H2**, consistent with its lower *in vitro* covalent modification efficiency. In contrast, compound **H6**, which lacks covalent reactivity due to methyl substitution at the 3-position, and the noncovalent analogue **H14** (Table 1), which does not contain the epoxide motif, produced only partial inhibition of the TR-FRET signal (27.7 ± 0.5% and 40.3 ± 1.9% at 100 μM, respectively) (Fig. S5), underscoring the benefit of covalent engagement.

Cellular activity of compounds **H1** and **H2** was further evaluated using an established r(CUG)^exp^–MBNL1 nano-bioluminescence resonance energy transfer (NanoBRET) assay,^23^ which provides a quantitative readout of small-molecule disruption of the r(CUG)^exp^–MBNL1 interaction in cells. In this assay, r(CUG)^exp^ templates the formation of a complex between two different MBNL1 fusion proteins–one with NanoLuciferase and the other with HaloTag to generate a NanoBRET signal (Fig. S6A).

Compounds **H1** and **H2** produced dose-dependent decreases in NanoBRET signal, with IC_50_ values of 5 ± 2 μM and 3 ± 1 μM, respectively (Fig. S6B). In contrast, compound **H6** (analogue that does not react with r(CUG) repeats *in vitro*) and the noncovalent analogue **H14** reduced NanoBRET signal by about 40% at 20 μM (Fig. S6B), the highest dose tested. Cell viability remained above 80% for all compounds across the tested concentration range (40 nM – 20 μM; Fig. S6C), indicating that reductions in NanoBRET signal were not attributable to cytotoxic effects. Collectively, both *in vitro* TR-FRET and cellular NanoBRET results suggest that covalent modification of r(CUG)^exp^ enhances disruption of the r(CUG)^exp^–MBNL1 interaction relative to noncovalent analogues.

### Structure-Guided Design of Epoxide- and Aziridine-2-carboxamide Covalent Ligands for the FMN Riboswitch

The FMN riboswitch, a well-characterized RNA element located in the 5′ leader sequence of bacterial mRNAs,^24^ was selected for structure-guided design of covalent ligands incorporating epoxide- and aziridine-2-carboxamide warheads. Examination of the crystal structure of FMN bound to the *Fusobacterium nucleatum* FMN riboswitch (PDB ID: 2YIE) showed that guanine residues G10 and G11 adopt conformations with accessible N7 positions within the FMN binding pocket (Fig. 4A).^25^

**Figure 4.**
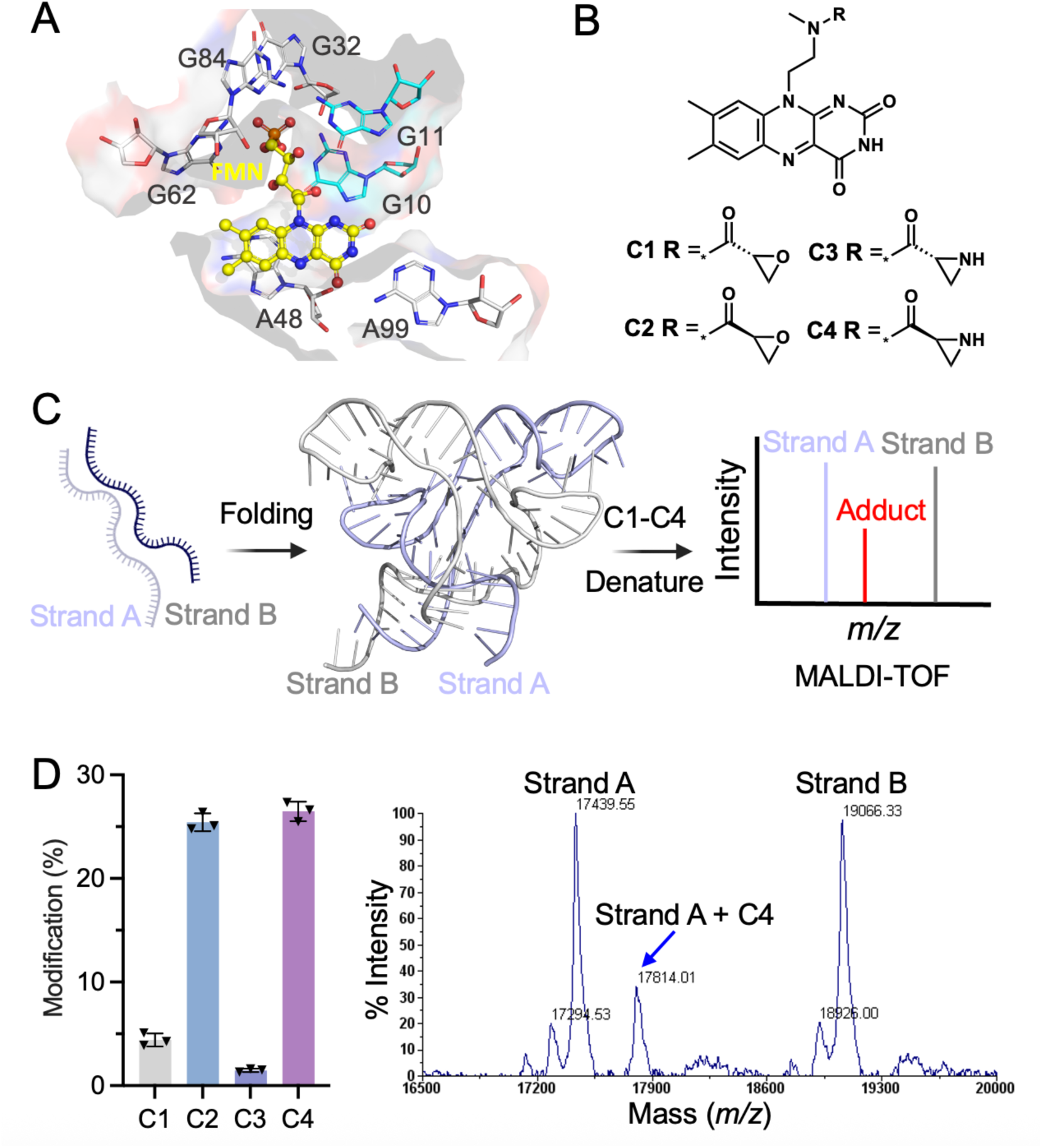
Structure-guided design of epoxide- and aziridine-2-carboxamide covalent ligands for the FMN riboswitch. A) Crystal structure of the *Fusobacterium nucleatum* FMN riboswitch bound to FMN (yellow). Residues G10 and G11 are highlighted as cyan stick (PDB ID: 2YIE). B) Chemical structures of designed compound **C1**-**C4**. C) Schematic of the MALDI-TOF-MS-based workflow for evaluating covalent modification of the FMN riboswitch. D) Left: Modification of strand A of the FMN riboswitch by **C1**-**C4** (n = 3). Right: Representative MALDI-TOF mass spectrum for the reaction of **C4** (200 μM) with FMN riboswitch (10 μM) at 37 °C for 12 h. The **C4**–strand A adduct peak is indicated with a blue arrow.

Two pairs of enantiomeric compounds were designed to assess stereochemical effects on covalent modification using epoxide- (**C1** and **C2**) and aziridine-2-carboxamide warheads (**C3** and **C4**) (Fig. 4B). A flavin core was used as the binding scaffold, and the distance between G10/G11 and the flavin core guided selection of an ethyl linker to connect the warheads to the scaffold. A methyl substituent was introduced at the amide linkage to restrict warhead conformation and orient the electrophile toward the N7 positions of G10 and G11.

Compounds **C1**–**C4** were synthesized and evaluated for the stability in FMN riboswitch folding buffer (50 mM HEPES, pH 6.5, 100 mM KCl, and 15 mM MgCl_2_). Epoxide-2-carboxamide derivatives **C1** and **C2** showed moderate instability, with approximately 78% of the parent compound remaining after 60 h of incubation at 37 °C (Fig. S7A). In contrast, aziridine-2-carboxamide derivatives **C3** and **C4** exhibited higher stability, with approximately 95% of the parent compound remaining under the same conditions.

Subsequently, covalent modification of the *F. nucleatum* FMN riboswitch was measured using a previously reported MALDI-TOF-MS workflow.^3^ In brief, the FMN riboswitch is reconstituted by co-folding two RNA strands, dubbed strand A and strand B (Fig. 4C). The *S*-enantiomers, epoxide-2-carboxamide **C2** and aziridine-2-carboxamide **C4**, showed comparable levels of covalent modification when 200 μM of the compound was incubated with 10 μM of the folded FMN riboswitch, achieving 25.4 ± 0.8% and 26.5 ± 0.9% selective modification of strand A after a 12 h incubation at 37 °C (Fig. 4D, Fig. S7B-D). In contrast, the corresponding *R*-enantiomers, **C1** and **C3**, showed less than 5% modification under the same conditions. These findings indicate that stereochemical control can strongly influence covalent engagement.

High-resolution LC–MS analysis was performed on enzymatically digested mononucleotides derived from FMN riboswitch samples modified with compounds **C2** and **C4** (Fig. S8). In both cases, only **C2**–guanosine and **C4**–guanosine adducts were detected. Corresponding fragment ions assigned to the guanine adducts (lacks ribose moiety) were also observed, indicating selective modification of the guanine base. Consistent with results obtained for r(CUG)_4_, the observed adducts are compatible with N7-directed covalent modification (Fig. S8).

### Crosslinking Site Identification of Compound C4 in the FMN Riboswitch

Compound **C4** was selected for crosslinking site identification in the FMN riboswitch based on its stability and covalent activity. Covalent modification of RNA can generate reverse transcriptase (RT) stops or induce RT-mutations at the site of crosslinking, an approach widely used in glyoxal-,^6^ DMS-,^26^ and SHAPE-based^27^ sequencing methodologies. Because **C4** selectively modified strand A of the FMN riboswitch, RT-stop and RT-mutation assays were performed using a strand A specific primer following previously established procedures (Fig. 5A).^3, 5^

**Figure 5.**
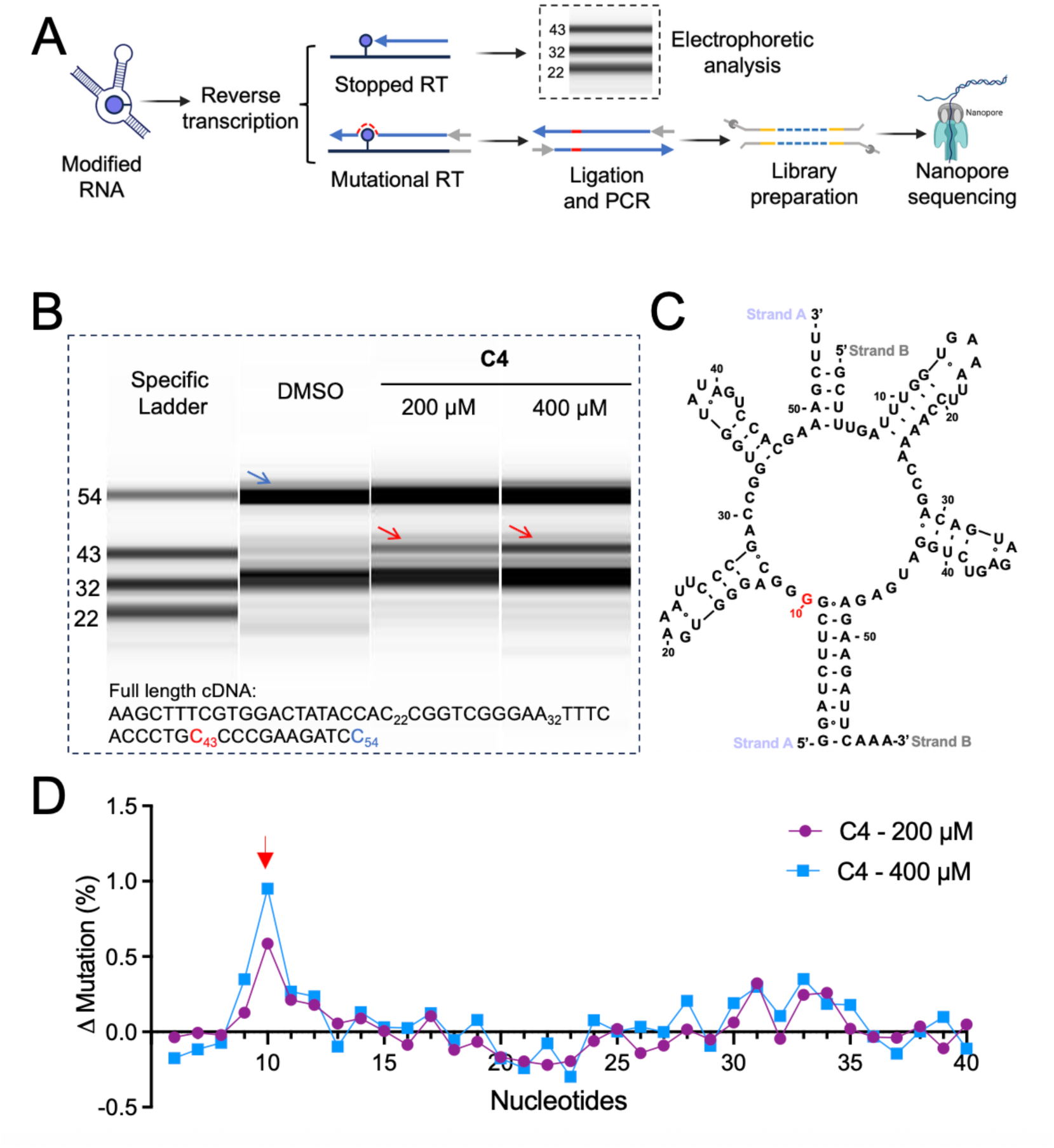
Investigation of the covalent modification site in the FMN riboswitch by C4. A) Schematic of RT stop analysis and RT-mutational mapping workflow. B) Representative fragment analyzer electropherograms of cDNA products generated from strand A of the FMN riboswitch. The full-length cDNA (54 nt) and the **C4**-induced RT-stop cDNA product (∼43 nt) were assigned using a custom DNA ladder (“Specific Ladder”) and are indicated by blue and red arrows, respectively. C) Mapping of the RT-stop site onto the FMN riboswitch secondary structure. D) Representative RT-mutation profile of strand A after reaction with **C4**, obtained by nanopore sequencing.

Complementary DNA (cDNA) products generated from the RT-stop assay were analyzed by fragment analyzer capillary electrophoresis using customized ladders to enable quantitative mapping of stop sites. Relative to the DMSO control, treatment with **C4** resulted in the appearance of a prominent truncated cDNA species of approximately 43 nucleotides (Fig. 5B), corresponding to a stop site proximal to G10 on strand A of the FMN riboswitch (Fig. 5C). The intensity of this truncated cDNA species increased at 400 μM relative to 200 μM, indicating dose-dependent formation of the RT-stop (Fig. 5B).

An RT-mutation assay coupled with nanopore sequencing was additionally employed to identify the **C4** crosslinking site. Relative to the DMSO control, **C4** treatment produced a pronounced mutation signal at G10 on strand A, which also exhibited clear dose dependence (Fig. 5D). Together, the RT-stop and RT-mutation analyses identify G10 on strand A as the site of covalent modification by compound **C4**, as designed.

### Bioactivity Evaluation of C1–C4 in *Bacillus subtilis* PY79

Biological activity of compounds **C1**–**C4** was evaluated using a LacZ reporter assay in *Bacillus subtilis* PY79 following previously established procedures.^3^ In this system, LacZ expression is regulated by an IPTG-inducible promoter and the *B. subtilis ribD* FMN riboswitch, and reporter activity is quantified by monitoring the cleavage of *o*-nitrophenyl-*β*-D-galactopyranoside (ONPG) to *o*-nitrophenol (ONP).^28^ In the absence of FMN, the riboswitch adopts an antiterminated state that permits LacZ expression, whereas in the presence of FMN, transcriptional termination represses LacZ expression (Fig. 6A).^24^ In the presence of IPTG, strong LacZ expression was observed (red, Fig. 6B,C), and repression of LacZ expression occurred upon addition of 100 μM riboflavin to the growth medium (green, Fig. 6B,C).

**Figure 6:**
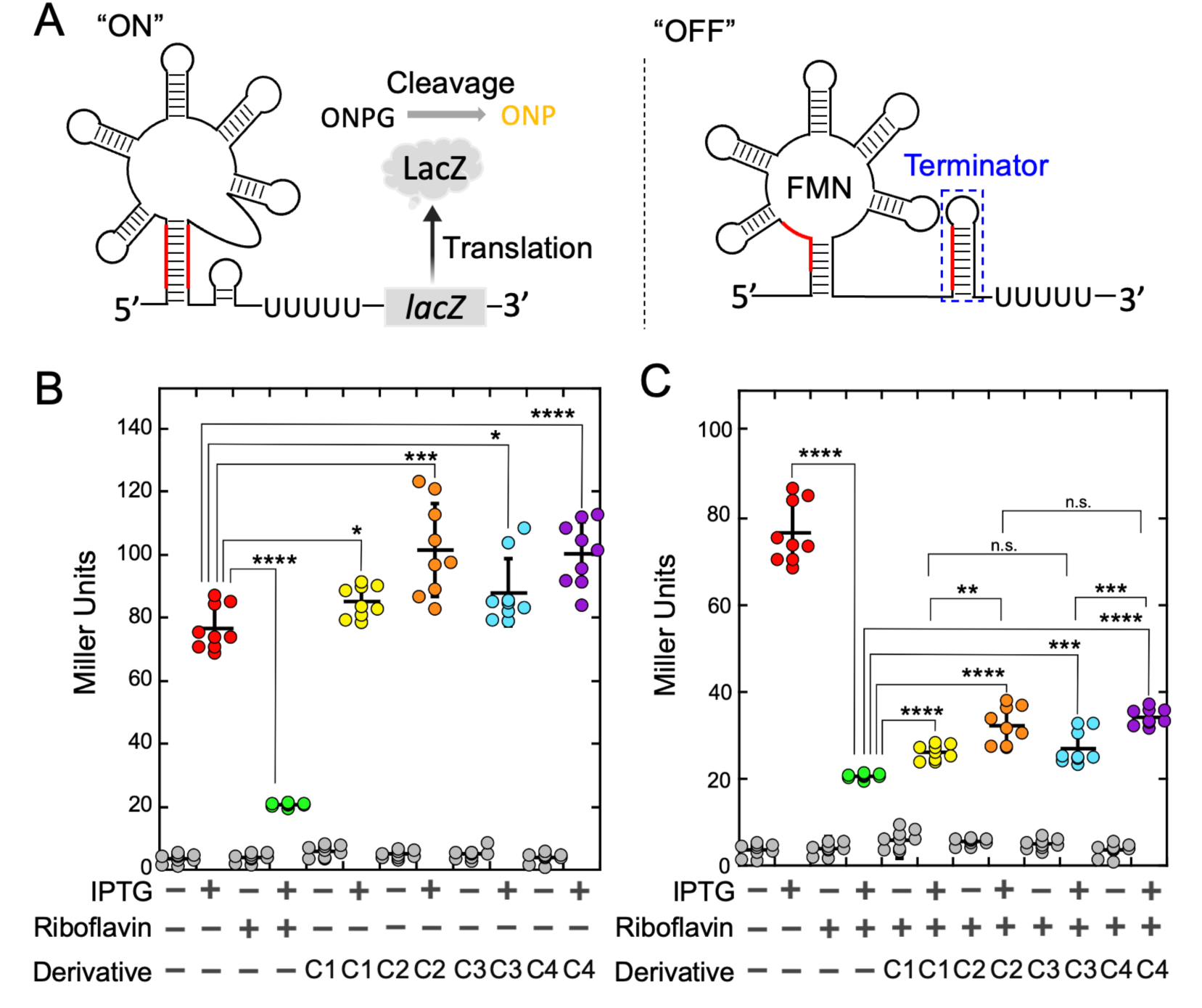
Bioactivity of compounds C1–C4. A) Schematic diagram of the FMN-regulated *B. subtilis ribD* riboswitch controlling expression of an mRNA encoding a LacZ reporter protein. Left: In the absence of bound FMN, the riboswitch enables read-through transcription and expression of LacZ, which cleaves *o*-nitrophenyl-*β*-D-galactopyroanoside (OPNG) to *o*-nitrophenol (ONP; yellow), quantified by using the Miller assay.^28^ Right: In the presence of riboflavin added to the growth medium, the riboswitch forms a terminator element, causing RNA polymerase to abort mRNA synthesis. B) LacZ activity of the *ribD*/*lacZ* reporter. LacZ expression is induced by IPTG (1 mM) and repressed by the addition of 100 µM riboflavin to the growth medium. Compounds **C2** and **C4** display statistically significant levels of *lacZ* expression above that induced by IPTG alone; 200 µM of **C1** and **C3** display levels of *lacZ* expression similar to IPTG alone. C) Compounds **C2** and **C4** exhibit competitive inhibition of *lacZ* repression by riboflavin, while **C1** and **C3** show weak competitive inhibition of *lacZ* repression. Error bars in panels B and C represent one standard deviation (s.d.); asterisks denote *P* values calculated using a two-sample *t*-test assuming unequal variances (\**P*<0.05, \*\**P*<0.01, \*\*\**P*<0.001, and \*\*\*\**P*<0.0001).

Compounds **C1**–**C4** produced modulation of FMN riboswitch activity consistent with *in vitro* assays. In the presence of 200 μM of **C2** or **C4**, a modest but reproducible increase in LacZ expression was observed (orange and purple, Fig. 6B). A similar stimulatory effect has been reported for a phenylglyoxal derivative acting as an inverse agonist of the FMN riboswitch.^3^ In contrast, **C1** and **C3** produced significantly lower effects on FMN riboswitch activity (yellow and cyan, Fig. 6B), consistent with their reduced covalent modification observed *in vitro* (Fig. 4C).

The ability of **C1**–**C4** to compete with riboflavin was further evaluated in the presence of 100 μM riboflavin in the growth medium (Fig. 6C). Under these conditions, LacZ expression in the presence **C2** and **C4** showed less attenuation than that observed for **C1** and **C3**. No appreciable difference was observed between **C1** and **C3** in their ability to compete with riboflavin, nor between **C2** and **C4**. These observations are consistent with a correlation between *in vitro* covalent modification and cellular modulation of FMN riboswitch activity.

## DISCUSSION

Progress in the development of RNA-targeted covalent small molecules has been limited in part by the scarcity of electrophiles that combine predictable reactivity with stability under biological conditions. In this study, epoxide- and aziridine-2-carboxamides were identified as tunable, mechanistically defined electrophiles capable of covalently modifying RNA. Structure–activity relationship studies delineated how substitution pattern and stereochemistry govern reactivity and stability, while mechanistic analyses indicated selective modification at guanine N7. Together, these findings introduce a new class of electrophiles that addresses a key limitation in covalent RNA ligand development.

A general framework for the rational design of RNA-targeted covalent small molecules remains underdeveloped. Here, such a framework is illustrated through application of epoxide- and aziridine-2-carboxamide warheads to two distinct structured RNA targets. Covalent ligands targeting the pathogenic r(CUG)^exp^ repeat RNA were generated by rational design and shown to modify the RNA and disrupt RNA–protein interactions in cells. In parallel, structure-guided installation of these warheads on a flavin scaffold yielded covalent modulators of the FMN riboswitch with validated reaction sites and cellular activity. Together, these examples demonstrate the applicability of this chemistry across disparate RNA architectures and biological contexts.

A notable feature of both systems is the pronounced stereoselectivity observed in covalent modification, underscoring the importance of chirality in RNA-targeted covalent ligand design. While stereochemical effects are well established in noncovalent RNA recognition, their role in covalent RNA modification has received little attention. The results presented here indicate that stereochemical control can strongly influence covalent engagement and may represent an important design parameter for future RNA-reactive electrophiles.

The chemical behavior of these warheads parallels that of electrophiles successfully employed in protein-targeted covalent inhibitors. Epoxide-containing motifs are well precedented for covalent protein modification, including in the FDA-approved proteasome inhibitor carfilzomib,^29^ and aziridines have been widely used to target nucleophilic amino acid residues.^21^ These similarities suggest that epoxide- and aziridine-2-carboxamides constitute a chemically versatile class of electrophiles whose selectivity for RNA or protein targets may be dictated by scaffold context, positioning, and molecular recognition rather than intrinsic reactivity alone.

In summary, this work defines epoxide- and aziridine-2-carboxamides as a coherent class of RNA-reactive electrophiles with tunable stability, predictable reactivity, and stereochemical control. Although demonstrated here with two RNA targets, the mechanistic clarity and compatibility with structure-guided placement suggest that this chemistry may be extendable to other structured RNAs presenting accessible guanine residues. These findings expand the toolkit for covalent RNA ligand design and highlight opportunities at the interface of RNA- and protein-directed covalent chemistry.

## ASSOCIATED CONTENT

### Supporting Information

The following data can be found in the Supporting Information. (i) Figures S1−S8; (ii) Materials and Methods; and (iii) synthetic methods and characterization.

### Author Contributions

The manuscript was written through contributions of all authors. All authors have given approval to the final version of the manuscript.

### Notes

The authors declare no competing financial interest.

## ACKNOWLEDGMENT

This work was supported by the U.S. National Institutes of Health R35 NS116846 (to M.D.D.), R01 P0377752 (to M.D.D. and N.C.M.) and R35 GM152029 (to R.T.B.).

## ABBREVIATIONS

cDNA: complementary DNA
DM1: myotonic dystrophy type 1
DMS: dimethyl sulfate
FMN: flavin mononucleotide
IPTG: isopropyl-*β*-D-1-thiogalactopyranoside
MALDI-TOF-MS: matrix-assisted laser desorption/ionization-time of flight-based mass spectrometry
MBNL1: muscleblind-like 1 protein
m/z: mass-to-charge ratio
NanoBRET: nano-bioluminescence resonance energy transfer
ONPG: *o*-nitrophenyl-*β*-D-galactopyroanoside
ONP: *o*-nitrophenol
RT: reverse transcriptase
SAR: structure-activity relationships
TAMRA: tetramethylrhodamine
TR-FRET: time-resolved fluorescence resonance energy transfer.

## TOC Graphic

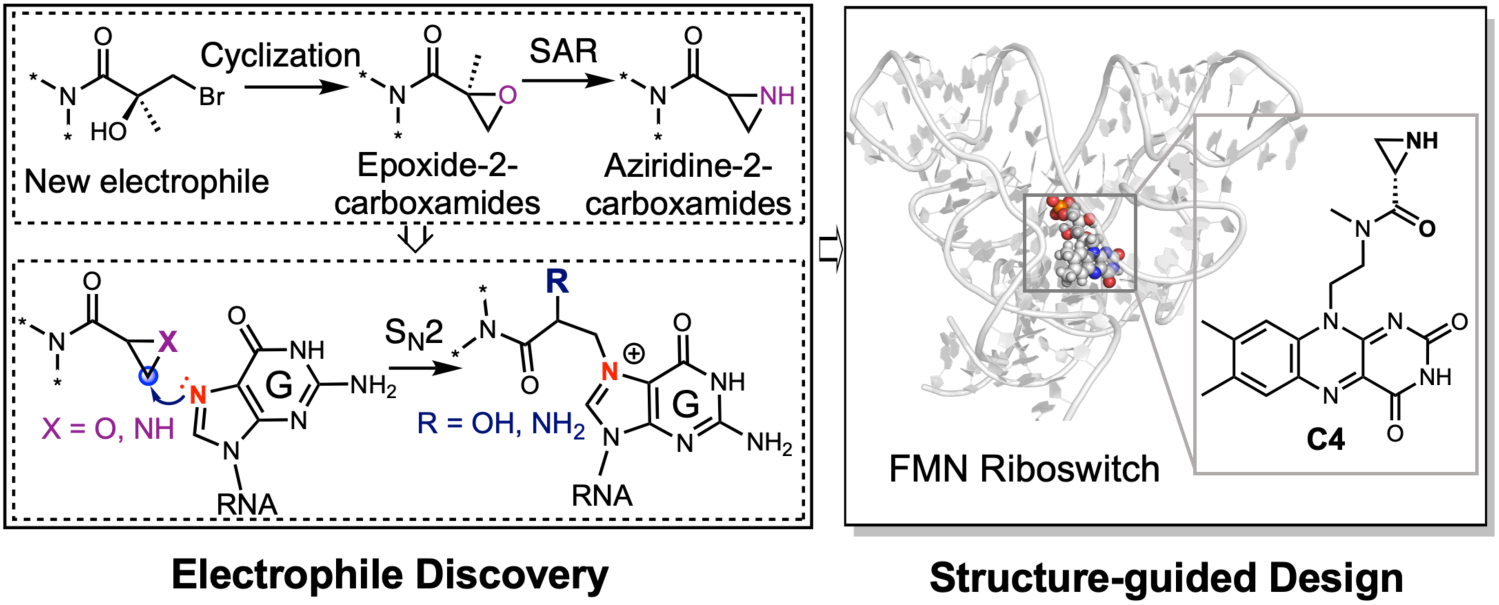

