## Supporting information for "Mechanistically Defined Epoxide- and Aziridine-2-carboxamide Electrophiles Enable Stereoselective Covalent RNA Modulation"

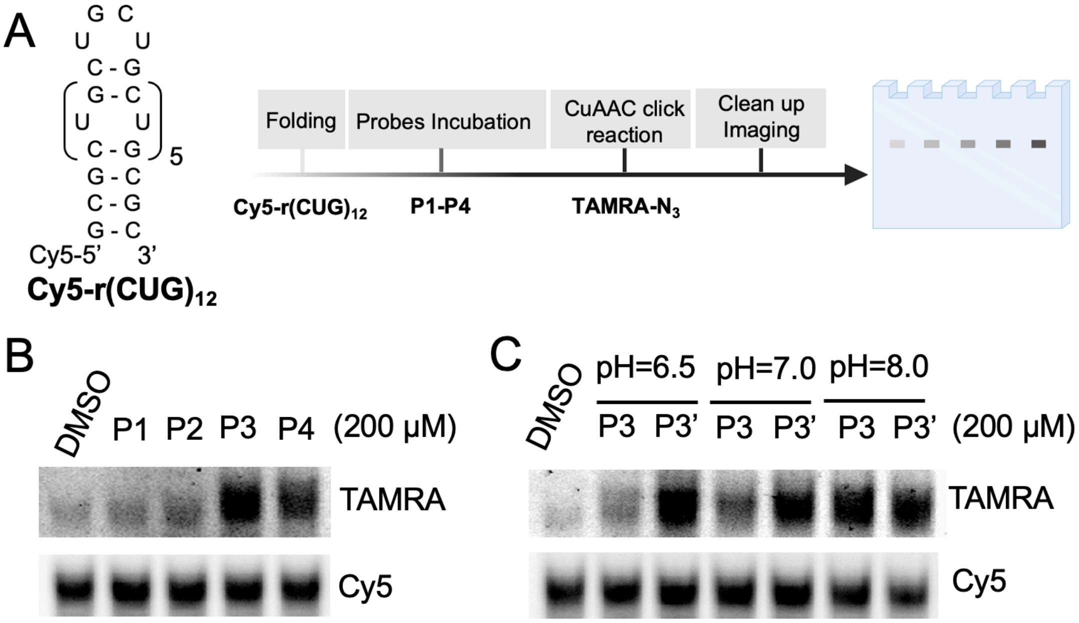


**Figure S1**. **Characterization of covalent RNA modification by TAMRA labeling assay. A**) Left: Secondary structure of Cy5-labeled r(CUG)$_{12}$. Right: Schematic representation of the TAMRA labeling assay workflow. **B)** TAMRA labeling of Cy5-labeled r(CUG)$_{12}$ (2 μM) by **P1-P4** (200 μM) at 37 °C for 12 h in a buffer containing 50 mM HEPES, pH 7.0, 100 mM KCl, and 15 mM $MgCl_2$. **C)** TAMRA labeling of Cy5-labeled r(CUG)$_{12}$ (2 μM) by **P3** (200 μM) and **P3'** (200 μM) in buffer ranging from pH 6.5 to 8.0.

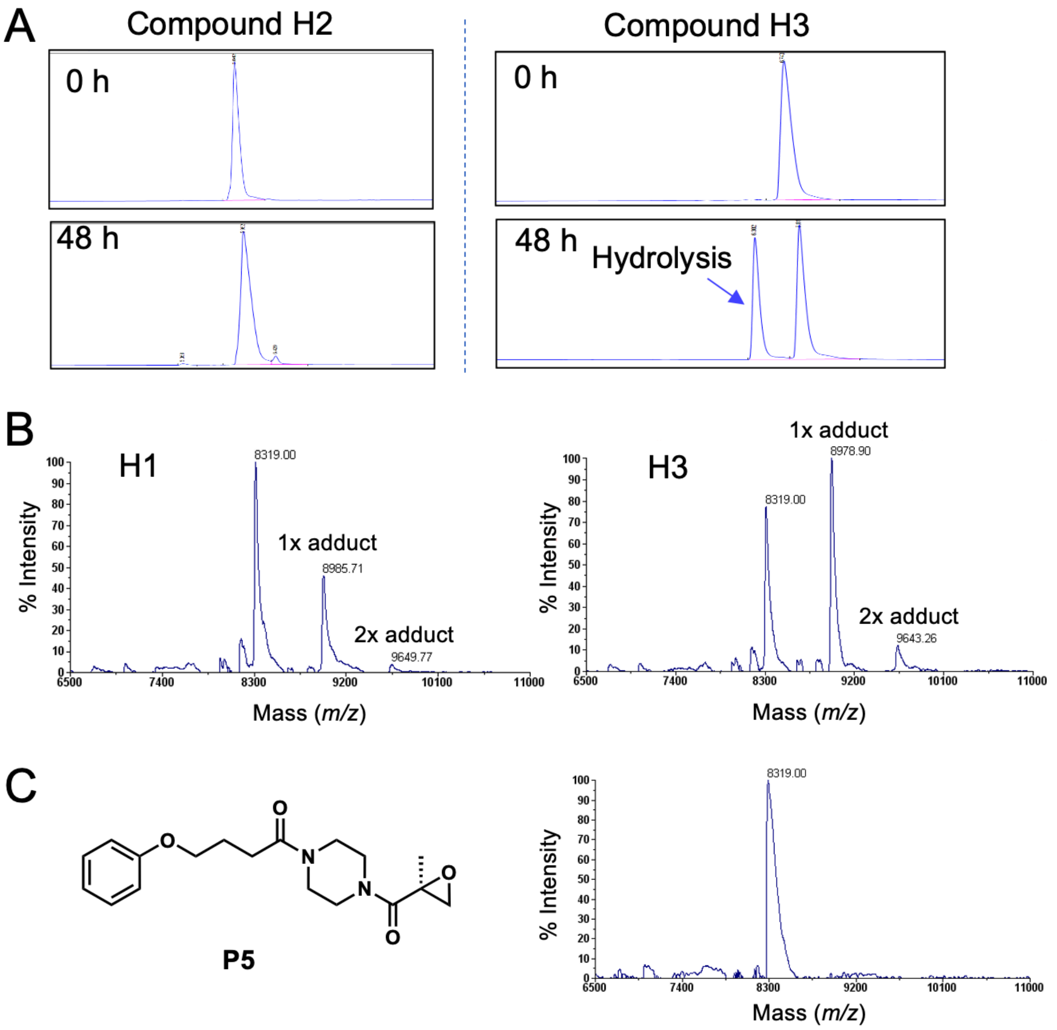


**Figure S2. A)** Representative LC-MS traces of **H2** and **H3** showing their hydrolytic stability in the reaction buffer (50 mM HEPES, pH 6.5,100 mM KCl, and 15 mM $MgCl_2$) at 37 °C. The hydrolysis product of **H3** is indicated by a blue arrow. **B)** Left: Representative MALDI-TOF mass spectrum of $r(CUG)_4$ (2 μM) after reaction with **H1** (200 μM) at 37 °C for 12 h. Right: Representative MALDI-TOF mass spectrum of $r(CUG)_4$ (2 μM) after reaction with **H3** (200 μM) at 37 °C for 12 h. **C)** Left: Chemical structure of compound **P5.** Right: Representative MALDI-TOF mass spectrum of $r(CUG)_4$ (2 μM) after reaction with **P5** (200 μM) at 37 °C for 12 h.

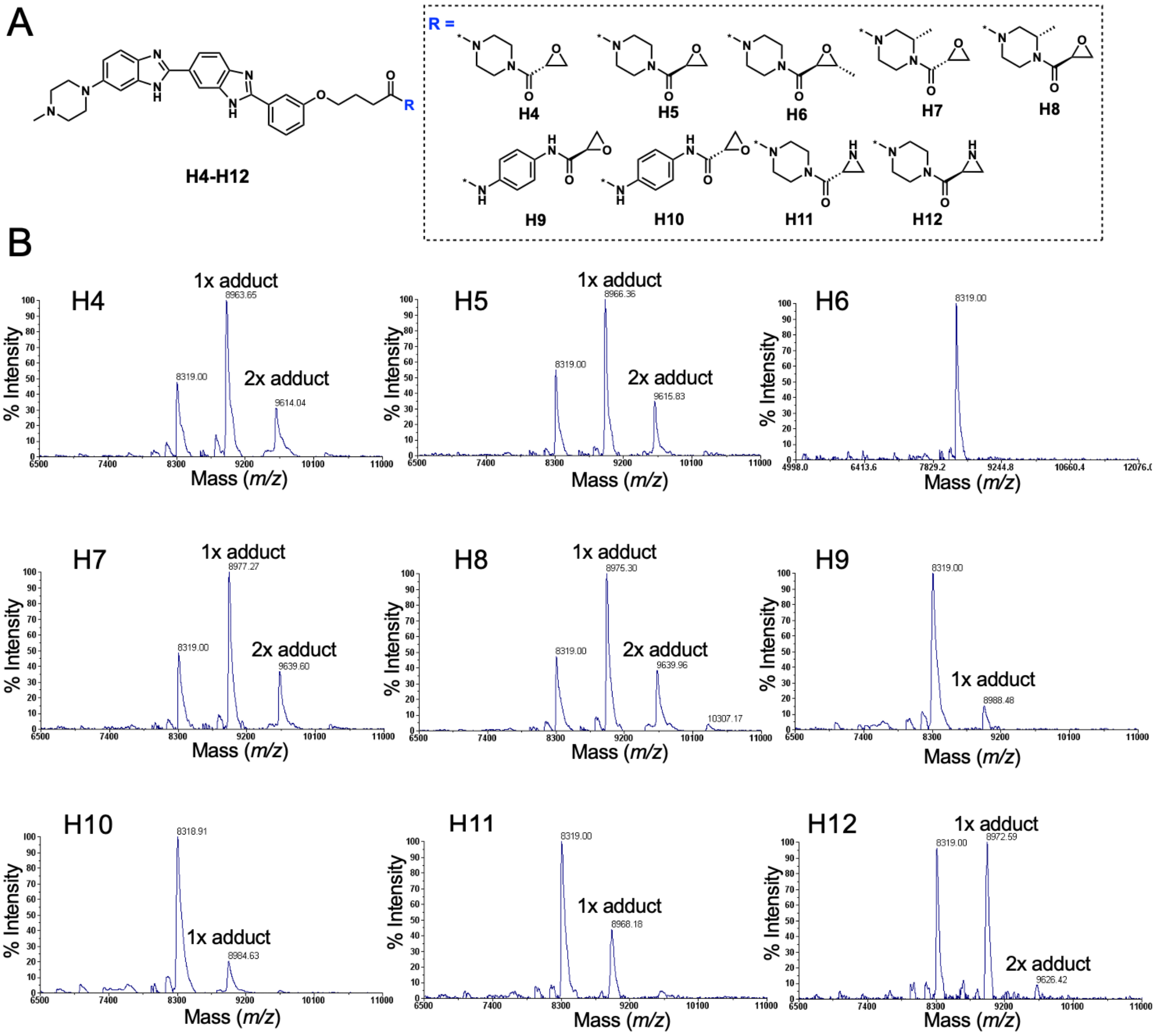


**Figure S3. A)** Chemical structure of compounds **H4-H12**. **B)** Representative MALDI-TOF mass spectrum of r(CUG)$_4$ (2 μM) after incubation with compounds **H4-H12** (200 μM) at 37 °C for 12 h.

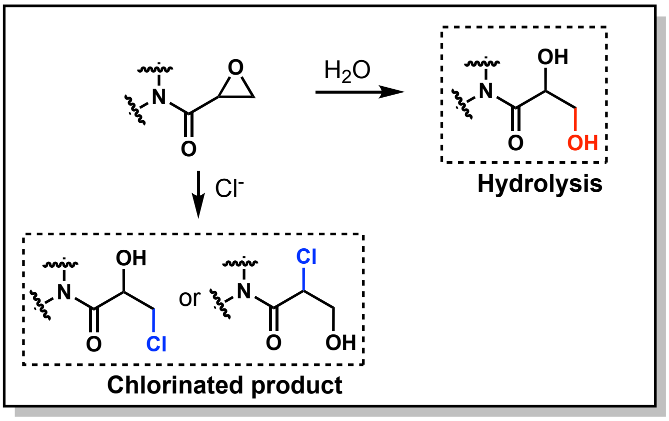


**Figure S4.** Proposed degradation pathways of epoxide-2-carboxamide warheads, as supported by LC–MS analysis. Epoxide ring opening occurs *via* hydrolysis to generate the corresponding glycol product (red), and through nucleophilic attack by chloride in buffer to yield chlorinated adducts (blue).

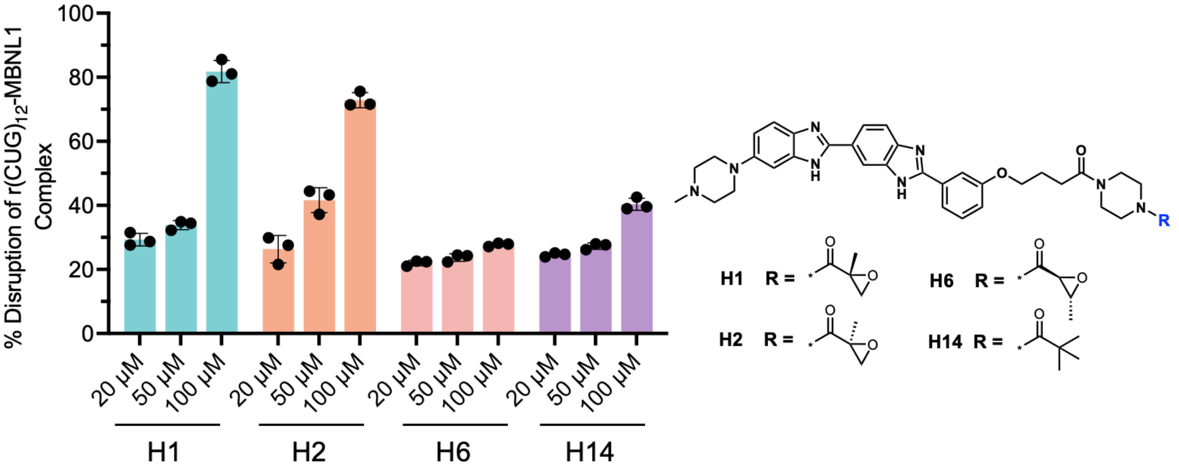


**Figure S5.** *In vitro* disruption of r(CUG)$_{12}$–MBNL1 complex by compounds **H1** and **H2** using a TR-FRET assay.[1] Compounds (20 μM – 100 μM) were preincubated with biotinylated r(CUG)$_{12}$ (320 nM) at 37 $^{o}$C for 12 h to allow covalent modification. Left: The percentage disruption of the r(CUG)$_{12}$–MBNL1 complex by the indicated compounds were calculated relative to DMSO control. Data are shown as mean ± SD (n = 3). Right: Chemical structures of the tested compounds.

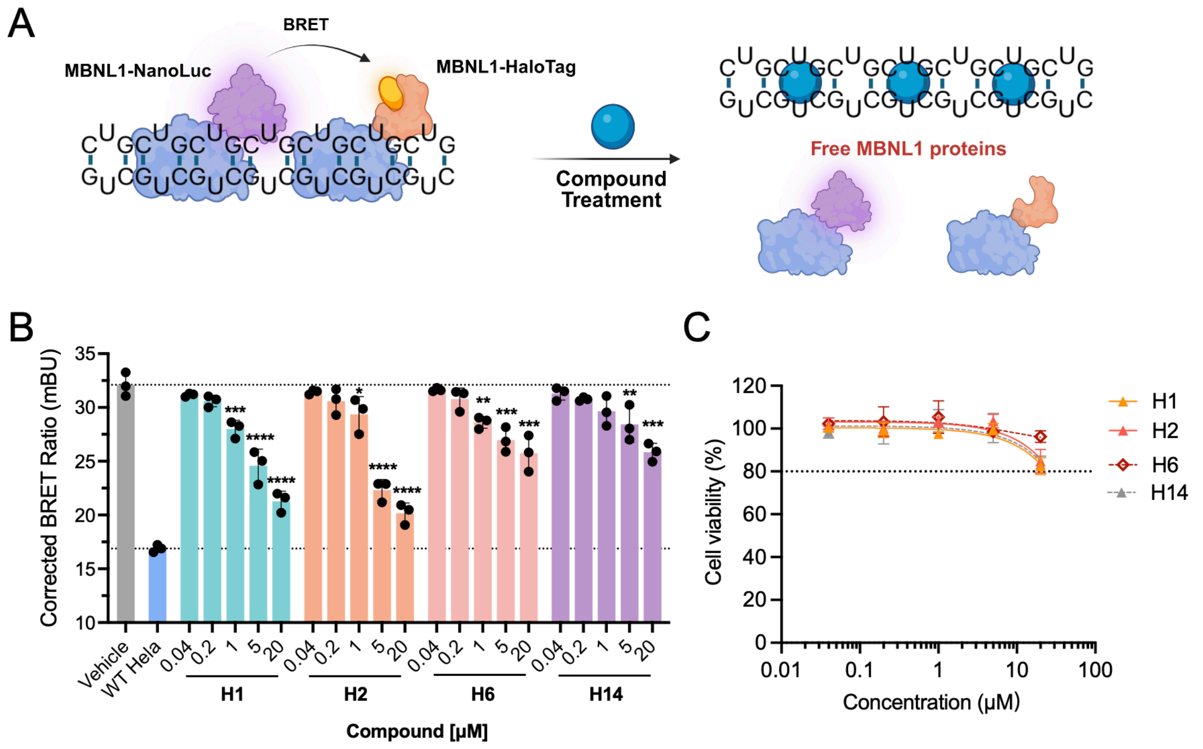


**Figure S6. Cellular activity and viability of covalent compounds H1 and H2. A)** Schematic of nano-bioluminescence resonance energy transfer (NanoBRET) assay in HeLa480 cells, which stably express 480 r(CUG) repeats.[2] The proximity of MBNL1-NanoLuc and MBNL1-HaloTag upon binding to r(CUG)$^{exp}$ enables resonance energy transfer. Treatment with compounds targeting r(CUG)$^{exp}$ dissociates the MBNL1 proteins from the RNA, thereby quenching the BRET signal. **B)** Dose−response analysis of **H1** and **H2** that disrupt r(CUG)$^{exp}$−MBNL1 complex formation. **H6** (analogue that does not react with r(CUG) repeats *in vitro*) and **H14** (analogue lacking a reactive electrophile) were selected as control compounds that do not (**H6**) or cannot (**H14**) covalently modify r(CUG)$^{exp}$ but can reduce NanoBRET by a simple binding mode of action. **B)** Cell viability of HeLa480 cells treated under the same conditions. All compounds maintained >80% viability across the tested concentration range. Data are shown as mean ± SD (n = 3). Statistical significance was determined using two-tailed unpaired Student's *t*-test relative to the vehicle. Significance is indicated as $p < 0.05$ (*), $p < 0.01$ (**), $p < 0.001$ (***), $p < 0.0001$ (****).

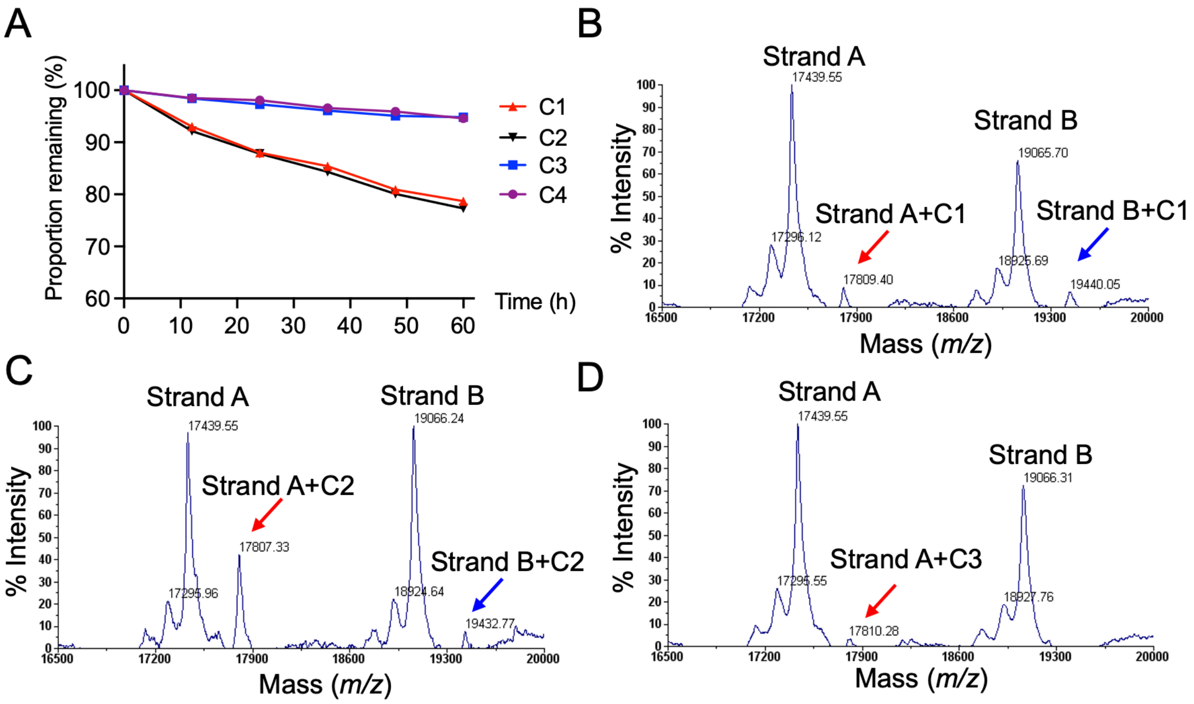


**Figure S7. A**) Stability of compounds **C1-C4** in reaction buffer (50 mM HEPES, pH 6.5, 100 mM KCl, and 15 mM $MgCl_2$). **B-D**) Representative MALDI-TOF mass spectrum of FMN riboswitch (10 μM) treated with **C1**, **C2**, and **C3** (200 μM, 37 °C for 12 h). Peaks corresponding to the covalently modified strand A and strand B are indicated by red and blue arrows, respectively.

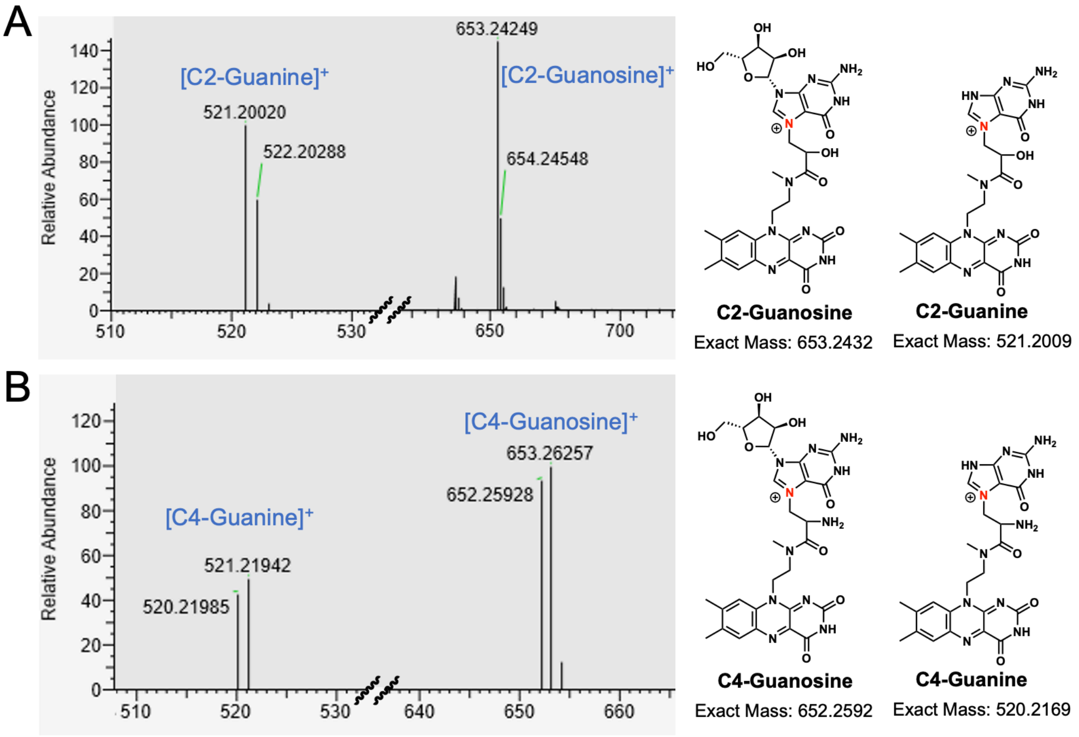


**Figure S8. High-resolution LC–MS identification of guanine adducts formed upon reaction of C2 and C4 with the FMN riboswitch. A)** Left: Extracted high-resolution MS spectrum of enzymatically digested nucleotides derived from **C2**-modified FMN riboswitch. Right: Possible chemical structure of the **C2**-guanosine and **C2**-guanine adducts. **B)** Left: Extracted high-resolution MS spectrum of enzymatically digested nucleotides derived from **C4**-modified FMN riboswitch. Right: Possible chemical structure of the **C4**-guanosine and **C4**-guanine adducts.

**Table S1.** Sequences of oligonucleotides used in this study. [α]

| Name | Sequence | Assay |
|---|---|---|
| Cy5-r(CUG)$_{12}$ | 5'-GCGCUGCUGCUGCUGCUGCUGCUGCUGCUGCU GCUGCUGCGC-3' | TAMRA |
| r(CUG)$_4$ | 5'-GACCUGCUGGUGAAAACCUGCUGGUC-3' | MALDI-TOF |
| Riboswitch strand A | 5'-GGAUCUUCGGGGCAGGGUGAAAUUCCCGACCG GUGGUAUAGUCCACGAAAGCUU-3' | MALDI-TOF |
| Riboswitch strand B | 5'-GCUUUGAUUUGGUGAAAUUCCAAAACCGACAGU AGAGUCUGGAUGAGAGAAGAUUCAAA-3' | MALDI-TOF |
| Riboswitch strand A RT primer | 5'-AAGCTTTCGTGGAC-3' | RT-STOP |
| ssDNA ladder | 5'-AAGCTTTCGTGGACTATACCAC-3'<br>5'-AAGCTTTCGTGGACTATACCACCGGTCGGGAAT-3'<br>5'-AAGCTTTCGTGGACTATACCACCGGTCGGGAATTT CACCCTGC-3'<br>5'-AAGCTTTCGTGGACTATACCACCGGTCGGGAATTT CACCCTGCCCCGAAGATCC-3' | Fragment analyzer |
| Riboswitch strand A RT primer | 5'-GCGAGCACAGAATTAATACGACTCACTATA GGTTT TTTTTTTTTAAGCT-3' | RT-mutation |
| Ligation adaptor | 5'-/5phos/AGATCGGAAGAGCGTCGTGTAG/3spc/-3' | Nanopore Sequencing |
| PCR primer of strand A | Forward: 5'-CTACACGACGCTCTTCCGATCT-3'<br>Reverse: 5'-GCGAGCACAGAATTAATACGAC-3' | Nanopore Sequencing |

[α] All RNA constructs were custom obtained from Horizon Discovery, and DNA constructs were ordered from Integrated DNA Technologies, Inc. (IDT). All oligonucleotides were purified by the vendor using a standard HPLC method.

### MATERIALS AND METHODS

#### RNA folding

*Cy5-r(CUG)$_{12}$ and r(CUG)$_4$*

The RNAs were dissolved in Nanopure water to a 100 μM stock concentration. For folding, RNA was diluted to 2 μM in 1× folding buffer (50 mM HEPES, and 100 mM KCl). Folding of Cy5-r(CUG)$_{12}$ was carried out at pH 6.5, 7.0, or 8.0, while r(CUG)$_4$ was folded at pH 6.5. The RNAs solution was heated at 95 °C for 3 min, snap cooled on ice for 3 min, and then supplemented with $MgCl_2$ to final concentration of 15 mM. The mixture was incubated on ice for additional 5 min, followed by incubation at 37 °C for 10 min prior to use.

*FMN riboswitch*

Two chemically synthesized strand A (10 μM) and Strand B (10 μM) were mixed in 1× folding buffer (50 mM HEPES, pH 6.5, and 100 mM KCl). The mixture was heated at 95 °C for 3 min, cooled down to 25 °C by 3 °C/min, and then supplemented with $MgCl_2$ to final concentration of 15 mM. The mixture was incubated at 37 °C for 30 min prior to use.

#### TAMRA labeling assay

The TAMRA labeling assay was performed as previously described.[3] Folded Cy5-r(CUG)$_{12}$ (2 μM) samples were incubated with probe compounds (200 μM) at 37 °C for 12 h in a final reaction volume of 20 μL, then cleaned up using RNAClean XP beads (Beckman Coulter, catalog #A63987), and eluted in Nanopure $H_2O$ (10 μL; see "*RNA clean-up*" below). The eluted, cleaned-up RNA was then incubated with 5 μL of a "click mix" containing tris(3-hydroxypropyltriazolylmethyl)amine (THPTA, 20 mM, 0.5 μL), copper sulfate ($CuSO_4$, 20 mM, 0.1 μL), sodium ascorbate (100 mM, 0.4 μL), TAMRA-$N_3$ (20 mM, 0.1 μL) and Nanopure $H_2O$ (3.9 μL) at 37 °C for 1 h. The RNA was cleaned up using RNAClean XP beads and eluted in Nanopure $H_2O$ (10 μL). RNA concentrations were determined using a NanoDrop-2000 spectrophotometer (Thermo

Fisher Scientific). The TAMRA-labeled RNA was analyzed using a urea denaturing acrylamide gel (11%, w/v), which was imaged by using a ChemiDoc™ MP System (TAMRA: excitation 550 nm, emission 565 nm; Cy5: excitation 651 nm, emission 670 nm).

**RNA clean-up**

RNAs were cleaned up using an optimized RNAClean XP bead protocol. RNAClean XP beads (1.8× the RNA sample volume) and isopropanol (2.5× the RNA sample volume) were added to the RNA solution, and the sample was incubated at room temperature for 10 min. The beads were pelleted using a magnetic rack, the supernatant was removed, and the beads were washed three times with 85% (v/v) ethanol. RNA was eluted with Nanopure $H_2O$ in a volume adjusted according to experimental requirements, followed by magnetic separation to remove the beads.

**Stability of compounds in reaction buffer**

Compounds (1 mM) were incubated at 37 °C in buffer containing 50 mM HEPES, pH 6.5, 100 mM KCl, and 15 mM $MgCl_2$. A 10 μL aliquot of the reaction mixture was withdrawn and diluted with 40 μL of MeOH/$H_2O$ (1:1, v/v) at the indicated time points. Each sample (10 μL) was immediately injected into an Agilent Infinity II liquid chromatography system coupled with an Agilent 6120 quadrupole LC/MS. The proportion of compound remaining was quantified by integrating the LC absorbance at 254 nm, and the identity of transformed products was confirmed by MS.

**Identification of covalent modification by MALDI-TOF-MS**

*r(CUG)$_4$ sample preparation*

Folded r(CUG)$_4$ (2 μM) was incubated with compound of interest (100 μM, 200 μM) in a total volume of 20 μL at 37 °C for 12 h. The RNA was then cleaned up using RNAClean XP beads, and eluted with 10 μL of Nanopure $H_2O$.

*FMN riboswitch sample preparation*

FMN riboswitch samples were prepared with an additional denaturation step as previously described.[3] Briefly, folded FMN riboswitch RNA (10 μM) was incubated with compounds (200 μM) in a total volume of 20 μL at 37 °C for 12 h and cleaned up using RNAClean XP beads. The RNA was eluted in 10 μL of Nanopure $H_2O$, followed by the addition of urea (8 M, 10 μL) and incubation at 37 °C for 10 min to dissociate noncovalently bound compounds. The mixture was then cleaned up using RNAClean XP beads, and the RNA was eluted in 10 μL of Nanopure $H_2O$ for analysis.

*MALDI-TOF-MS analysis*

Covalent RNA modification by small molecules was analyzed by MALDI-TOF mass spectrometry as previously described.[4] Briefly, purified RNA (1 μL) was spotted onto a MALDI plate and air-dried. Next, 2′,4′,6′-Trihydroxyacetophenone monohydrate (THAP, 1μL) matrix, prepared from 50 mg/mL diammonium citrate and 18 mg/mL THAP in $H_2O$/acetonitrile (1:1, v/v), was added to the dried sample spot and allowed to crystallize at room temperature. Mass spectra were collected on an Applied Biosystems 4800 Plus MALDI-TOF instrument (laser intensity: 7400), acquiring sub-spectra with a signal-to-noise threshold >15. Spectra were processed using Applied Biosystems/SciEX Data Explorer software. The degree of covalent modification was determined by integrating the peak areas corresponding to unmodified and modified RNA species.

Given most compounds formed a primary singly modified adduct (1× adduct) and a low-abundance doubly modified adduct (2× adduct) with $r(CUG)_4$, the extent of modification for the **H**-scaffold compounds (**H1**–**H13**) was calculated using the following equation:

$$\text{Adducts per r(CUG)}_4 = \frac{A_{1\times} + 2A_{2\times}}{A_{1\times} + A_{2\times} + A_{\text{unmodified}}}$$

where $A_{1x}$, $A_{2x}$, an $A_{unmodified}$ represent the integrated peak areas of the singly modified (1× adduct), doubly modified (2× adduct), and unmodified RNA species, respectively.

For the FMN riboswitch, the percentage of covalent modification was calculated using the following equation:

$$\text{Modification (\%)} = \frac{A_{\text{adduct}}}{A_{\text{adduct}} + A_{\text{unmodified}}} \times 100\%$$

**Nucleoside digestion to identify reaction mechanism by high-resolution LC–MS**

The nucleoside digestion was performed using the manufacturer's protocol for the Nucleoside Digestion Mix (New England Biolabs, catalog #M0649S). Covalently modified RNA (500 ng in 17 μL Nanopure $H_2O$), as estimated by NanoDrop UV absorbance after cleanup, was added the manufacturer-supplied reaction buffer (10×, 2 μL) and Nucleoside Digestion Mix (1 μL). The resulting mixture was incubated at 37 °C for 1 h. A 10 μL aliquot of the digestion mixture was diluted with 40 μL of MeOH/$H_2O$ (1:1, v/v) and analyzed by a Vanquish HPLC system (Thermo Fisher Scientific). Mass spectra of the resulting nucleosides were recorded in positive ESI mode on an Orbitrap Exploris 120 (Thermo Fisher Scientific).

**Evaluation of r(CUG)$_{12}$–MBNL1 disruption by an *in vitro* TR-FRET assay**

Disruption of r(CUG)$_{12}$–MBNL1 complex was evaluated by using a previously established time-resolved fluorescence resonance energy transfer (TR-FRET) assay.[1] Biotinylated r(CUG)$_{12}$ (320 nM) was folded by heating at 95 °C for 3 min in TR-FRET reaction buffer (20 mM HEPES, pH 7.5, 110 mM KCl, 10 mM NaCl) and then snap cooling on ice for 10 min. The folded RNA was incubated with compounds at 37 °C for 12 h to allow the covalent modification. Compound-treated r(CUG)$_{12}$ (320 nM, 10 μL) was combined with MBNL1 (267 nM, 9 μL) in assay buffer (20 mM HEPES, pH 7.5, 110 mM KCl, 10 mM NaCl, 4 mM $MgCl_2$, 4 mM $CaCl_2$, 10 mM dithiothreitol (DTT), 0.2% (w/v) BSA, and 0.1% (v/v) Tween-20) . The mixture was incubated for 15 min at room temperature, followed by addition of streptavidin-XL665 (3.2 μM, 0.5 μL; Revvity, catalog #610SAXLF) and terbium-Anti-His$_6$ antibody (35 ng/μL, 0.5 μL; Revvity, catalog #61HISTLF) to reach a total volume of 20 μL. Maximum TR-FRET controls contained r(CUG)$_{12}$ and MBNL1 with an equivalent volume of DMSO in place of test compounds, whereas minimum TR-FRET controls were prepared in the absence of MBNL1, with the protein volume replaced by assay buffer.

TR-FRET signals were recorded on a SpectraMax M5 plate reader in time-resolved mode (200 μs delay, 1500 μs integration). Emission at 545 nm and 665 nm was measured following excitation at 345 nm (420 nm cut-off). The TR-FRET ratio (545/665) was used to quantify $r(CUG)_{12}$–MBNL1 interaction, and inhibition was calculated using the following equation:

$$Inhibition\ rate = \frac{TR-FRET_{maximum} - TR-FRET_{sample}}{TR-FRET_{maximum} - TR-FRET_{minimum}} \times 100\%$$

**Cellular activity evaluation by a NanoBRET assay**

*Cell culture*

HeLa480[5] and wild type HeLa cells were cultured in Dulbecco's Modified Eagle Medium (DMEM; Corning, catalog #15-017-CV) supplemented with 10% (v/v) fetal bovine serum (FBS; Gibco, catalog #12676029), 1% (v/v) Antibiotic-Antimycotic Solution (Corning, catalog #30-004-CI), and 1% (v/v) Glutagro (Corning, catalog #25-015-CI). Cells were maintained at 37 °C in a humidified atmosphere with 5% $CO_2$ and used at passage numbers below 20. All cells were routinely tested and confirmed to be mycoplasma-free prior to use.

*NanoBRET: live-cell $r(CUG)^{exp}$-MBNL1 displacement assay*

Cellular formation of the $r(CUG)^{exp}$-MBNL1 complex was measured using a previously established NanoBRET assay.[2] HeLa480 and wild type HeLa cells were co-transfected in 6-well plate with a plasmid encoding MBNL1-HaloTag (2500 ng) and MBNL1-NanoLuc (50 ng) using FuGENE HD transfection reagent (8 μL, Promega, catalog #E2311) according to the manufacturer's protocol. After overnight incubation, cells were harvested and reseeded into white 96-well plates (Corning, catalog #3903) at $1.6 \times 10^4$ cells per well in assay medium (Opti-MEM supplemented with 4% (v/v) FBS) containing 200 nM HaloTag NanoBRET 618 ligand (Promega, catalog #N1662).

For each experiment, cells were also dispensed into three wells containing 100 µL of assay medium alone to serve as no–HaloTag ligand controls. The cells were allowed to adhere for 4 h at 37 °C prior to treatment with compounds (0.5% (v/v) DMSO). Following 16 h of incubation, NanoLuc substrate (1×, Promega, catalog #N1662) and extracellular NanoLuc inhibitor (20 µM, Promega, catalog #N2160) were added in 25 µL assay medium per well. The NanoBRET signal was measured using the GloMax Discover System (Promega, catalog #GM3000) with dual-filtered luminescence detection: a 460/80 nm bandpass filter for the NanoLuc donor and a 610 nm long-pass filter for HaloTag acceptor with an integration time of 500 ms. The corrected NanoBRET ratio, expressed in milliBRET units (mBU), was calculated using following equation:

$$\text{Corrected NanoBRET Ratio} = \frac{\text{Halotag ligand (Emission 610nm/LP)}}{\text{Donor Luminescence (Emission 450nm/BP 80nm)}} - \frac{\text{No Ligand Control (Emission 610nm/LP)}}{\text{Donor Luminescence (Emission 450nm/BP 80nm)}}$$

The $IC_{50}$ values were calculated by fitting dose-response curve to a four-parameter logistic (4PL) equation:

$$Inhibition\ rate = \frac{100}{1 + 10^{(\log IC50 - concentration)\times Hill\ Slope}}$$

*Cell viability*

Cell viability was measured using the CellTiter-Glo 2.0 Cell Viability Assay (Promega, catalog #G9242) following NanoBRET assay readout. After NanoBRET measurements, CellTiter-Glo 2.0 Reagent was equilibrated to room temperature and added to each well at a 1:1 ratio to the culture medium volume. The plates were shaken at 500–700 rpm for 30 min at room temperature to allow cell lysis and quench the NanoLuc signal. After incubation, total luminescence was recorded using the GloMax Discover System according to the manufacturer's protocol.

**Reverse transcription for RT-Stop analysis**

Reverse transcription of covalently modified RNA was performed using SuperScript III (Invitrogen, catalog #18-080-093) following the manufacturer's protocol. To compound **C4**-modified FMN riboswitch RNA (200 ng in 4.2 μL nuclease-free $H_2O$), a strand A specific RT primer (5 μM, 1 μL, Table S1) was added. The mixture was heated at 60 °C for 5 min and snap cooled to 4 °C for 5 min in a thermocycler to allow primer annealing. A master mix containing 10 mM dNTPs (1 μL), 5× First Strand Buffer (2 μL), 0.1 M dithiothreitol (DTT, 0.5 μL), DMSO (0.3 μL), RNaseOUT (0.5 μL; Invitrogen, catalog #10777019), and SuperScript III (0.5 μL) was then added to afford a final reaction volume of 10 μL. The reaction was incubated at 50 °C for 10 min, followed by enzyme inactivation at 85 °C for 5 min. After cooling to room temperature, RNase T1 (5 units, 1 μL; Thermo Scientific, catalog #EN0542) and RNase H (5 units, 1 μL; NEB, catalog #M0297L) were added, and the mixture was incubated at 37 °C for 30 min to degrade the RNA template. The resulting cDNA was purified using Ampure XP beads (Beckman Coulter, catalog #A63881) and eluted in 10 μL of Nanopure $H_2O$.

**Analyzing RT-cDNA products using a fragment analyzer**

cDNA products generated from RT were analyzed on an Agilent 5300 Fragment Analyzer using the manufacturer's protocol for the Small RNA Kit (Agilent, catalog #DNF-470-0275). Prior to analysis, cDNA was denatured at 70 °C for 10 min and diluted with 18 μL of Small RNA Diluent (provided with the kit). Electrophoretic separation was performed with a 30 s pre-run at 11.5 kV, followed by a 50 s injection at 8.0 kV and a 45 min separation at 11.5 kV. Fragment sizes were assigned using a custom ssDNA ladder (22–54 nt) composed of oligonucleotides matching the sequence of strand A of the FMN riboswitch (Table S1).

**RT-mutation and Nanopore sequencing analysis**

*Poly(A) tailing of modified RNA.*

Poly(A) tailing of **C4**-modified FMN riboswitch RNA was carried out using *E. coli* poly(A) polymerase following the manufacturer's protocol. Briefly, 200 ng of RNA was mixed with 1 μL of 10× *E. coli* Poly(A) Polymerase reaction buffer (provided with the kit), 1 μL of 10 mM ATP (final concentration 1 mM), 1 μL of *E. coli* Poly(A) Polymerase (5 units), and nuclease-free water to afford a final reaction volume of 10 μL. The reaction was incubated at 37 °C for 30 min and subsequently purified using RNAClean XP beads. The polyadenylated RNA was eluted in nuclease-free water and used for RT-mutation step.

*Reverse transcriptional mutation by SuperScript II.*

RT-mutational analysis was carried out according to a reported protocol.[6] Briefly, polyadenylated RNA (100 ng in 8 μL nuclease-free $H_2O$) was mixed with polyA RT primer for strand A (5 μM, 1 μL, Table S1), heated at 65 °C for 5 min, and snap cooled to 4 °C for 5 min to anneal the primer. A freshly prepared 2.22× RT buffer (111 mM Tris–HCl, pH 8.0, 167 mM KCl, 13.3 mM $MnCl_2$, 22 mM DTT, and 2.22 M betaine; 9 μL) and 10 mM dNTPs (1 μL) were added, and the samples were incubated at room temperature for 2 min. SuperScript II (200 units, 1 μL; Invitrogen, catalog #18064014) was then added, and reverse transcription was carried out as follows: 25 °C for 10 min; 42 °C for 90 min; 10 cycles of 42 °C for 2 min and 50 °C for 2 min; followed by 70 °C for 10 min. The reaction was treated with RNase T1 (5 units, 1 μL; Thermo Scientific, catalog #EN0542) and RNase H (5 units, 1 μL; NEB, catalog #M0297L) at 37 °C for 30 min, purified using RNAClean XP beads, and eluted in 10 μL of Nanopure $H_2O$.

*Library preparation and nanopore sequencing*

For PCR amplification, cDNA generated by reverse transcription was ligated to a single strand DNA adaptor (Table S1) using CircLigase ssDNA Ligase (Biosearch Technologies, catalog #CL4111K) according to the manufacturer's instructions. The ligated cDNA was subsequently amplified by using Phusion High-Fidelity DNA Polymerase (NEB, catalog #M0530S), following the manufacturer's protocol. Sequencing libraries were carried out by using the Native Barcoding Kit 96 V14 (Oxford Nanopore, catalog #SQK-NBD114.96) following the recommended protocol and sequenced using a MinION R10.4.1 flow cell (Oxford Nanopore, catalog #FLO-

MIN114).

*RNA sequencing data analysis*

Nanopore sequencing reads were first base-called and demultiplexed using Dorado's 'basecaller' and 'demux' subcommands, respectively.[7] The processed base-called reads were converted to FASTQ format with 'samtools fastq' and aligned to the target sequence using 'bowtie2' with the '--local' option. Alignment outputs were generated as SAM files, which were subsequently converted to BAM files and sorted using samtools.[8] The alignment start site of each read on FMN strand A was extracted from each BAM file using a customed script. The mutation rates were calculated as the number of mutations divided by the coverage at each nucleotide position.

***Bacillus subtilis* reporter assay**

The bioactivity assay for **C1**–**C4** was performed using a previously described approach.[3] The reporter strain comprises the *B. subtilis ribD* riboswitch upstream of the *lacZ* gene under control of the isopropyl *β*-D-1-thiogalactopyranoside (IPTG)-inducible Pspank promoter inserted into the genome of *B. subtilis* strain P79 at the *amyE* locus.[3] LacZ activity was quantified using a modified Miller assay.[9] In brief, for each assay, the cells were cultured overnight at 37 °C in chemical salts broth (CSB) medium supplemented with 100 μg/mL spectinomycin.[10] For each reaction condition, three separate colonies were isolated and outgrown to provide three biological replicates. This culture was used to inoculate (1:50 dilution) 1 mL CSB medium with 1 mM IPTG and appropriate ligands (100 μM riboflavin and/or 200 μM compounds **C1**–**C4**). After outgrowth for approximately 6 h, the cells were harvested by centrifugation, washed with fresh CSB medium and resuspended in 1.5 mL of working buffer (60 mM $Na_2HPO_4$, 40 mM $NaH_2PO_4$, pH 7.0, 10 mM KCl, 1 mM $MgSO_4$ and 20 mM *β*-mercaptoethanol). This cellular suspension was divided into three equal volumes, representing three technical replicates. The absorbance at 600 nm for each reaction was determined, followed by lysis by adding 6.25 μL 15 mg/mL lysozyme and incubation for 20 min at 37 °C. Subsequently, 93.8 μL of 4 mg/mL *ortho*-nitrophenyl-*β*-

D-galactopyranoside (ONPG) was added to each reaction, which were incubated for an additional 15 min. Reactions were quenched with 250 µL of 1 M $Na_2CO_3$ and the absorbance taken at 420 nm. The Miller units for each reaction was calculated using the following equation:

$$Miller\ Units = 1000 * \frac{A420}{Reaction\ time\ (minutes) * Reaction\ Volume\ (mL) * A600}$$

where the A values represent the absorbance at a specific wavelength.

### General Synthetic Methods

NMR spectra were collected on a Bruker UltraShield™ NMR spectrometer (600 MHz). Silica gel flash column chromatography was conducted using a Biotage Isolera One purification system. Preparative reverse-phase HPLC was operated on a Waters system (Pump: Waters 1525; Absorbance detector: Waters 2487; Column: Waters Sunfire C18 OBD 5 μm, 19 × 150 mm S-14). Purification was carried out using a linear gradient elution from 0% to 100% methanol (MeOH) in water over 60 min at a flow rate of 5 mL/min. Analytical HPLC was used to assess compound purity on a Waters Symmetry C18 column (5 μm, 4.6 × 150 mm) at a flow rate of 1 mL/min under the same gradient and solvent conditions, with UV absorbance monitored at 254 nm. High-resolution mass spectra (HRMS) were acquired using an Orbitrap Exploris 120 mass spectrometer (Thermo Fisher Scientific) operating in positive electrospray ionization (ESI) mode, coupled to a Vanquish HPLC system (Thermo Fisher Scientific).

### Chemical synthesis

3-Chloro-*N*-(3-ethynylphenyl)-2,2-dimethylpropanamide (**P1**)

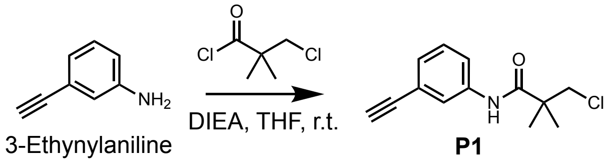


To a solution of 3-ethynylaniline (10 mg, 0.08 mmol) and *N,N*-diisopropylethylamine (DIEA, 13 mg, 0.10 mmol) in tetrahydrofuran (THF, 1 mL), was added 3-chloro-2,2-dimethylpropanoyl chloride (13 mg, 0.08 mmol) at 0 °C. The resulting mixture was stirred at 0 °C for 30 min and quenched by water (3 mL), following a purification by reverse-phase high-performance liquid chromatography (RP-HPLC, 0.1% (v/v) trifluoroacetic acid (TFA) in MeOH/$H_2O$) to obtain compound **P1** (14 mg, 74%). $^1$H NMR (600 MHz, DMSO-$d_6$) δ 9.44 (s, 1H), 7.78 (s, 1H), 7.63 (dd, *J* = 8.3, 1.2 Hz, 1H), 7.33 (t, *J* = 7.9 Hz, 1H), 7.18 (dt, *J* = 7.6, 1.3 Hz, 1H), 4.13 (s, 1H), 3.84 (s, 2H), 1.29 (s, 6H). $^{13}$C NMR (150 MHz, DMSO-$d_6$) δ 173.78, 139.52, 129.51, 127.29, 123.79, 122.23, 121.52, 83.76, 80.90, 52.83, 45.22, 23.46. HRMS: calculated for $C_{13}H_{14}ClNO$

[M + H]+ 236.0837, found 236.0836.

3-Bromo-*N*-(3-ethynylphenyl)-2,2-dimethylpropanamide (**P2**)

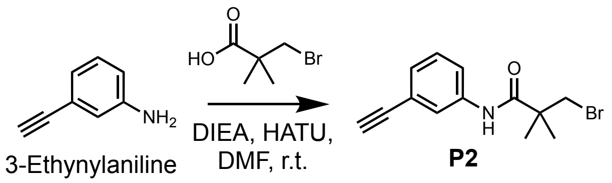


To a solution of 3-ethynylaniline (10 mg, 0.08 mmol) in *N,N*-dimethylformamide (DMF, 0.5 mL), was added 3-bromo-2,2-dimethylpropanoic acid (16 mg, 0.09 mmol), DIEA (31 mg, 0.24 mmol), and 2-(7-azabenzotriazol-1-yl)-*N',N',N'*-tetramethyluronium hexafluorophosphate (HATU, 38 mg, 0.10 mmol). The resulting mixture was stirred at room temperature for 3 h and then purified by RP-HPLC (0.1% (v/v) TFA in MeOH/$H_2O$) to obtain compound **P2** (11 mg, 48%). $^1$H NMR (600 MHz, DMSO-$d_6$) δ 9.42 (s, 1H), 7.78 (t, *J* = 1.9 Hz, 1H), 7.63 (ddd, *J* = 8.3, 2.2, 1.1 Hz, 1H), 7.32 (t, *J* = 7.9 Hz, 1H), 7.17 (dt, *J* = 7.6, 1.1 Hz, 1H), 4.16 (s, 1H), 3.77 (s, 2H), 1.32 (s, 6H). $^{13}$C NMR (150 MHz, DMSO-$d_6$) δ 173.51, 139.61, 129.48, 127.22, 123.76, 122.26, 121.44, 83.76, 80.98, 44.73, 43.16, 24.31. HRMS: calculated for $C_{13}H_{14}BrNO$ [M + H]+ 280.0332, found 280.0331.

(*S*)-3-Bromo-*N*-(3-ethynylphenyl)-2-hydroxy-2-methylpropanamide (**P3**)

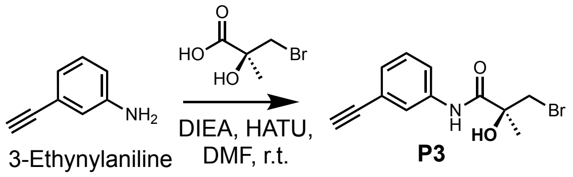


To a solution of 3-ethynylaniline (10 mg, 0.08 mmol) in DMF (0.5 mL), was added (*S*)-3-bromo-2-hydroxy-2-methylpropanoic acid (16 mg, 0.09 mmol), DIEA (31 mg, 0.24 mmol), and HATU (38 mg, 0.10 mmol). The resulting mixture was stirred at room temperature for 3 h and then purified by RP-HPLC (0.1% (v/v) TFA in MeOH/$H_2O$) to obtain compound **P3** (9 mg, 40%). $^1$H NMR (600 MHz, DMSO-$d_6$) δ 9.76 (s, 1H), 7.94

(t, *J* = 1.8 Hz, 1H), 7.75 (ddd, *J* = 8.2, 2.2, 1.0 Hz, 1H), 7.32 (t, *J* = 7.9 Hz, 1H), 7.19 (dt, *J* = 7.6, 1.2 Hz, 1H), 6.22 (brs, 1H), 4.17 (s, 1H), 3.82 (d, *J* = 10.4 Hz, 1H), 3.58 (d, *J* = 10.3 Hz, 1H), 1.47 (s, 3H). $^{13}$C NMR (150 MHz, DMSO-$d_6$) δ 172.89, 139.00, 129.47, 127.40, 123.32, 122.32, 121.16, 83.85, 81.03, 74.72, 42.34, 25.48. HRMS: calculated for $C_{12}H_{12}BrNO_2$ [M + H]$^+$ 282.0124, found 282.0124.

(*R*)-3-Bromo-*N*-(3-ethynylphenyl)-2-hydroxy-2-methylpropanamide (**P4**)

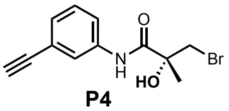

**P4**

Compound **P4** was prepared via a similar procedure to that of compound **P3**. $^1$H NMR (600 MHz, DMSO-$d_6$) δ 9.76 (s, 1H), 7.94 (t, *J* = 1.8 Hz, 1H), 7.75 (ddd, *J* = 8.3, 2.2, 1.1 Hz, 1H), 7.32 (t, *J* = 7.9 Hz, 1H), 7.19 (dt, *J* = 7.6, 1.1 Hz, 1H), 6.22 (brs, 1H), 4.18 (s, 1H), 3.82 (d, *J* = 10.3 Hz, 1H), 3.58 (d, *J* = 10.4 Hz, 1H), 1.47 (s, 3H). $^{13}$C NMR (150 MHz, DMSO-$d_6$) δ 172.89, 139.00, 129.47, 127.40, 123.32, 122.32, 121.16, 83.85, 81.03, 74.72, 42.34, 25.48. HRMS: calculated for $C_{12}H_{12}BrNO_2$ [M + H]$^+$ 282.0124, found 282.0125.

(*S*)-*N*-(3-Ethynylphenyl)-2-methyloxirane-2-carboxamide (**P3'**)

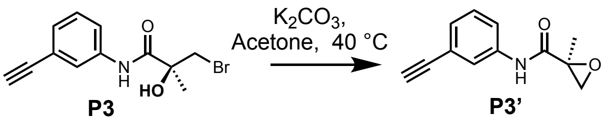


To a solution of **P3** (5 mg, 0.02 mmol) in acetone (0.5 mL), was added potassium carbonate ($K_2CO_3$, 5 mg, 0.03 mmol). The resulting mixture was stirred at 40 °C for 2 h and then purified by RP-HPLC (0.1% (v/v) TFA in MeOH/$H_2O$) to obtain compound **P3'** (3 mg, 88%). $^1$H NMR (600 MHz, DMSO-$d_6$) δ 9.56 (s, 1H), 7.85 (t, *J* = 1.9 Hz, 1H), 7.70 (ddd, *J* = 8.3, 2.3, 1.1 Hz, 1H), 7.31 (t, *J* = 7.9 Hz, 1H), 7.18 (dt, *J* = 7.6, 1.3 Hz, 1H), 4.17 (s, 1H), 3.02 (d, *J* = 5.2 Hz, 1H), 2.97 (d, *J* = 5.1 Hz, 1H), 1.53 (s, 3H). $^{13}$C NMR (150 MHz, DMSO-$d_6$) δ 169.64, 138.87, 129.43, 127.47, 123.59, 122.27, 121.36, 83.81, 81.06, 56.46, 53.35, 17.80. HRMS: calculated for $C_{12}H_{11}NO_2$ [M + H]$^+$ 202.0863, found 202.0862.

($R$)-1-(4-(2-Methyloxirane-2-carbonyl)piperazin-1-yl)-4-(3-(6-(4-methylpiperazin-1-yl)-1$H$,3'$H$-[2,5'-bibenzo[$d$]imidazol]-2'-yl)phenoxy)butan-1-one (**H1**)

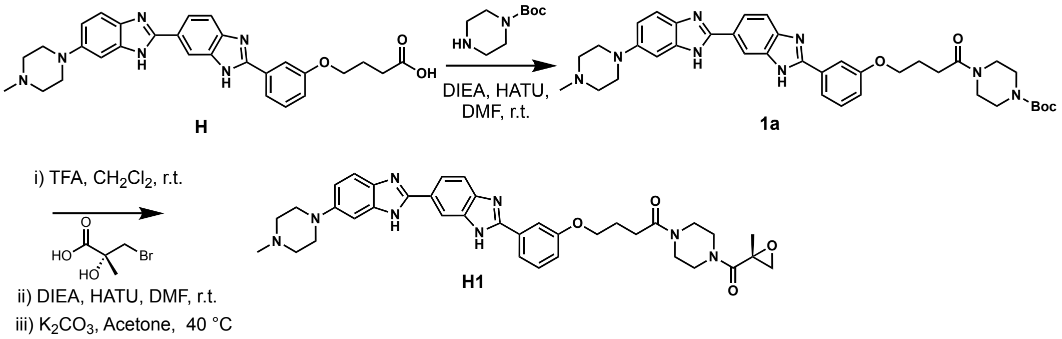


The intermediate **H** was prepared according to a reported procedure.[11] To a solution of intermediate **H** (100 mg, 0.20 mmol) in DMF (5 mL), was added *tert*-butyl piperazine-1-carboxylate (43 mg, 0.24 mmol), DIEA (77 mg, 0.60 mmol), and HATU (95 mg, 0.25 mmol). The resulting mixture was stirred at room temperature for 3 h, and then diluted with water (25 mL) and extracted with ethyl acetate (EtOAc, 20 mL). The organic layer was dried over anhydrous sodium sulfate ($Na_2SO_4$), filtered, and concentrated to afford a yellow solid, which was washed by dichloromethane ($CH_2Cl_2$)/hexanes (1:2, v/v) to get crude intermediate **1a**, which was used in the next step without further purification. LC-MS: calculated for $C_{38}H_{46}N_8O_4$ $[M + H]^+$ 679.4, found 679.3.

To a solution of intermediate **1a** (20 mg, 0.03 mmol) in dichloromethane (DCM, $CH_2Cl_2$, 3 mL), was added TFA (2 mL). The reaction mixture was stirred at room temperature for 3 h and then concentrated to remove solvent and excess TFA, affording a yellow solid, which was subsequently dissolved in DMF (0.5 mL). ($R$)-3-Bromo-2-hydroxy-2-methylpropanoic acid (7 mg, 0.04 mmol), DIEA (39 mg, 0.30 mmol), and HATU (15 mg, 0.04 mmol) were then added. The reaction mixture was stirred at room temperature for 3 h, diluted with $H_2O$ (15 mL) and extracted with EtOAc (10 mL). The organic layer was washed with brine, dried over anhydrous $Na_2SO_4$, filtered, and concentrated to give a yellow solid, which was dissolved in acetone (0.5 mL). $K_2CO_3$

(5 mg, 0.03 mmol) was then added. The resulting mixture was stirred at 40 °C for additional 3 h before filtration. The resulting solution was purified by RP-HPLC (0.1% TFA (v/v) in MeOH/$H_2O$) to afford compound **H1** (3 mg, 15% over three steps). $^1H$ NMR (600 MHz, DMSO-$d_6$) δ 9.91 (s, 2H), 8.47 (s, 1H), 8.05 (dd, $J$ = 8.5, 1.8 Hz, 1H), 7.90 (d, $J$ = 8.4 Hz, 1H), 7.83 (d, $J$ = 8.0 Hz, 1H), 7.81 (s, 1H), 7.71 (d, $J$ = 9.0 Hz, 1H), 7.52 (t, $J$ = 7.9 Hz, 1H), 7.31 (d, $J$ = 9.0 Hz, 1H), 7.22 (d, $J$ = 2.4 Hz, 1H), 7.16 (dd, $J$ = 8.3, 3.4 Hz, 1H), 4.14 (t, $J$ = 6.4 Hz, 2H), 3.92 (d, $J$ = 12.5 Hz, 2H), 3.64 – 3.32 (m, 12H), 3.06 (t, $J$ = 12.0 Hz, 2H), 2.91 (s, 3H), 2.88 (d, $J$ = 5.0 Hz, 1H), 2.83 (d, $J$ = 5.0 Hz, 1H), 2.57 (t, $J$ = 7.2 Hz, 2H), 2.03 (p, $J$ = 6.9 Hz, 2H), 1.45 (s, 3H). $^{13}C$ NMR (150 MHz, DMSO-$d_6$) δ 170.85, 167.92, 159.50, 154.47, 150.18, 148.69, 130.85, 122.34, 119.72, 117.61, 116.99, 113.03, 67.63, 56.79, 52.84, 52.59, 47.02, 45.38, 45.29, 45.01, 44.83, 42.57, 41.77, 41.48, 41.03, 29.11, 24.89, 19.92. HRMS (ESI) calculated for $C_{37}H_{42}N_8O_4$ [M + H]$^+$ 663.3402, found 663.3400. HPLC purity 97.97%.

(*S*)-1-(4-(2-Methyloxirane-2-carbonyl)piperazin-1-yl)-4-(3-(6-(4-methylpiperazin-1-yl)-1*H*,3'*H*-[2,5'-bibenzo[*d*]imidazol]-2'-yl)phenoxy)butan-1-one (**H2**)

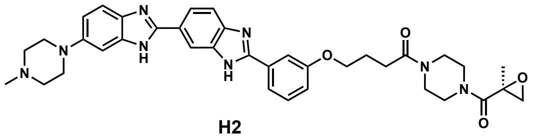

**H2**

Compound **H2** was prepared via a similar procedure to that of compound **H1**. $^1H$ NMR (600 MHz, DMSO-$d_6$) δ 10.00 (s, 2H), 8.48 (s, 1H), 8.07 (dd, $J$ = 8.4, 1.8 Hz, 1H), 7.92 (d, $J$ = 8.4 Hz, 1H), 7.85 (d, $J$ = 7.9 Hz, 1H), 7.82 (s, 1H), 7.73 (d, $J$ = 9.0 Hz, 1H), 7.53 (t, $J$ = 7.9 Hz, 1H), 7.33 (dd, $J$ = 9.0, 2.3 Hz, 1H), 7.24 (s, 1H), 7.17 (dd, $J$ = 8.3, 3.4 Hz, 1H), 4.15 (t, $J$ = 6.4 Hz, 2H), 3.94 (d, $J$ = 12.6 Hz, 2H), 3.66 – 3.37 (m, 12H), 3.08 (t, $J$ = 11.9 Hz, 2H), 2.92 (s, 3H), 2.90 (d, $J$ = 5.4 Hz, 1H), 2.84 (d, $J$ = 5.0 Hz, 1H), 2.58 (t, $J$ = 7.3 Hz, 2H), 2.04 (p, $J$ = 6.9 Hz, 2H), 1.46 (s, 3H). $^{13}C$ NMR (150 MHz, DMSO-$d_6$) δ 170.31, 167.36, 158.96, 153.97, 149.56, 148.23, 130.31, 121.83, 119.19, 117.10, 116.57, 112.49, 67.09, 56.25, 52.28, 52.06, 46.45, 44.83, 44.47, 44.30, 42.03, 41.15, 40.93, 28.57, 24.35, 19.38. HRMS (ESI) calculated for $C_{37}H_{42}N_8O_4$ [M + H]$^+$

663.3402, found 663.3397. HPLC purity 97.89%.

1-(4-(3-Chloro-2,2-dimethylpropanoyl)piperazin-1-yl)-4-(3-(6-(4-methylpiperazin-1-yl)-1*H*,3'*H*-[2,5'-bibenzo[*d*]imidazol]-2'-yl)phenoxy)butan-1-one (**H3**)

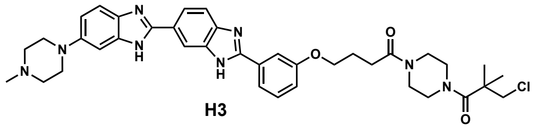

H3

Compound **H3** was prepared from intermediate **1a** following a procedure analogous to that used for **P1**. $^1$H NMR (600 MHz, DMSO-$d_6$) δ 10.07 (brs, 1H), 8.48 (s, 1H), 8.06 (dd, *J* = 8.4, 1.8 Hz, 1H), 7.92 (d, *J* = 8.5 Hz, 1H), 7.84 (d, *J* = 8.0 Hz, 1H), 7.81 (s, 1H), 7.72 (d, *J* = 9.0 Hz, 1H), 7.52 (t, *J* = 7.9 Hz, 1H), 7.33 (dd, *J* = 9.0, 2.3 Hz, 1H), 7.24 (d, *J* = 2.4 Hz, 1H), 7.16 (dd, *J* = 8.3, 1.7 Hz, 1H), 4.14 (t, *J* = 6.4 Hz, 2H), 3.93 (d, *J* = 13.0 Hz, 2H), 3.77 (s, 2H), 3.64 – 3.52 (m, 6H), 3.52 – 3.44 (m, 4H), 3.31 – 3.19 (m, 2H), 3.16 – 3.02 (m, 2H), 2.91 (s, 3H), 2.56 (t, *J* = 7.2 Hz, 2H), 2.03 (p, *J* = 6.9 Hz, 2H), 1.28 (s, 6H). $^{13}$C NMR (150 MHz, DMSO-$d_6$) δ 176.58, 172.69, 170.85, 159.50, 154.54, 150.01, 148.85, 130.85, 130.73, 122.41, 119.76, 117.69, 117.20, 114.97, 113.05, 99.83, 67.64, 53.96, 52.81, 52.79, 46.95, 45.25, 44.95, 43.96, 42.56, 41.60, 29.09, 24.87, 24.03. HRMS (ESI) calculated for $C_{38}H_{45}ClN_8O_3$ $[M + H]^+$ 697.3376, found 697.3373. HPLC purity 98.00%.

(*S*)-1-(4-(2-methyloxirane-2-carbonyl)piperazin-1-yl)-4-phenoxybutan-1-one (**P5**)

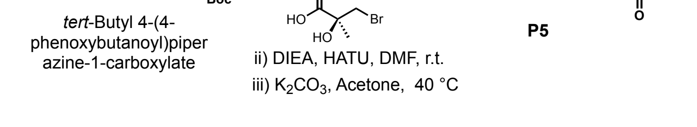


Compound **P5** was prepared from *tert*-butyl 4-(4-phenoxybutanoyl)piperazine-1-carboxylate following a procedure analogous to that used for **H1**. $^1$H NMR (600 MHz, $CDCl_3$) δ 7.28 (dd, *J* = 8.7, 7.4 Hz, 2H), 6.94 (t, *J* = 7.4 Hz, 1H), 6.89 (d, *J* = 7.7 Hz, 2H), 4.04 (t, *J* = 5.8 Hz, 2H), 3.88 – 3.35 (m, 8H), 2.96 (d, *J* = 4.8 Hz, 1H), 2.80 (d, *J*

= 4.9 Hz, 1H), 2.57 (t, *J* = 7.3 Hz, 2H), 2.20 – 2.08 (m, 2H), 1.58 (s, 3H). $^{13}C$ NMR (150 MHz, $CDCl_3$) δ 171.16, 168.47, 168.29, 158.77, 129.53, 120.80, 114.41, 66.63, 57.21, 57.10, 53.16, 53.07, 45.69, 45.28, 45.14, 45.01, 41.91, 41.84, 41.17, 29.42, 24.83, 19.95. HRMS (ESI) calculated for $C_{18}H_{24}N_2O_4$ $[M + H]^+$ 333.1809, found 333.1808.

(*R*)-4-(3-(6-(4-Methylpiperazin-1-yl)-1*H*,3'*H*-[2,5'-bibenzo[*d*]imidazol]-2'-yl)phenoxy)-1-(4-(oxirane-2-carbonyl)piperazin-1-yl)butan-1-one (**H4**)

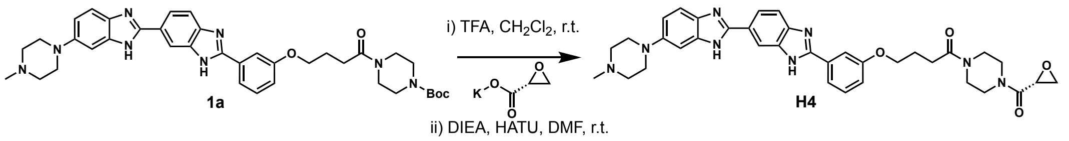


To a solution of intermediate **1a** (10 mg, 0.01 mmol) in $CH_2Cl_2$ (3 mL), was added TFA (2 mL). The reaction mixture was stirred at room temperature for 3 h and then concentrated to remove solvent and excess TFA, affording a yellow solid, which was subsequently dissolved in DMF (0.5 mL). Potassium (*R*)-oxirane-2-carboxylate (3 mg, 0.02 mmol), DIEA (39 mg, 0.30 mmol), and HATU (11 mg, 0.03 mmol) were then added. The reaction mixture was stirred at room temperature for 3 h and then purified by RP-HPLC (MeOH/$H_2O$ containing 0.1% ammonium bicarbonate ($NH_4HCO_3$)) to afford compound **H4** (4 mg, 42% over two steps). $^1H$ NMR (600 MHz, DMSO-$d_6$) δ 8.32 (s, 1H), 8.03 (d, *J* = 8.5 Hz, 1H), 7.80 (d, *J* = 7.9 Hz, 2H), 7.71 (d, *J* = 8.5 Hz, 1H), 7.48 (t, *J* = 7.9 Hz, 2H), 7.09 (dd, *J* = 8.2, 2.9 Hz, 1H), 6.94 (d, *J* = 9.0 Hz, 2H), 4.13 (t, *J* = 6.4 Hz, 2H), 3.88 (s, 1H), 3.77 – 3.44 (m, 10H), 3.13 (s, 3H), 2.90 (dd, *J* = 6.3, 4.4 Hz, 1H), 2.77 (d, *J* = 9.1 Hz, 1H), 2.63 – 2.54 (m, 2H), 2.24 (s, 2H), 2.03 (p, *J* = 6.8 Hz, 2H). $^{13}C$ NMR (150 MHz, DMSO-$d_6$) δ 170.89, 166.29, 159.44, 153.08, 148.13, 131.72, 130.64, 125.21, 121.40, 119.38, 116.98, 113.82, 112.61, 67.57, 50.34, 47.55, 47.47, 46.28, 46.02, 45.34, 44.84, 44.57, 42.06, 41.76, 41.66, 41.10, 29.16, 29.12, 24.90. HRMS (ESI) calculated for $C_{36}H_{40}N_8O_4$ $[M + H]^+$ 649.3245, found 649.3239.

(*S*)-4-(3-(6-(4-Methylpiperazin-1-yl)-1*H*,3'*H*-[2,5'-bibenzo[*d*]imidazol]-2'-yl)phenoxy)-1-(4-(oxirane-2-carbonyl)piperazin-1-yl)butan-1-one (**H5**)

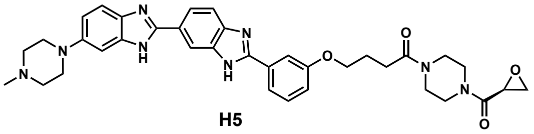

**H5**

Compound **H5** was prepared via a similar procedure to that of compound **H4.** [1]H NMR (600 MHz, DMSO-$d_6$) δ 8.30 (s, 1H), 8.01 (d, *J* = 8.4 Hz, 1H), 7.84 – 7.76 (m, 2H), 7.69 (d, *J* = 8.4 Hz, 1H), 7.51 – 7.38 (m, 2H), 7.07 (dd, *J* = 8.0, 3.0 Hz, 1H), 7.05 – 6.96 (m, 1H), 6.93 (dd, *J* = 8.8, 2.3 Hz, 1H), 4.11 (t, *J* = 6.4 Hz, 2H), 3.88 (s, 1H), 3.79 – 3.45 (m, 10H), 3.12 (s, 3H), 2.90 (dd, *J* = 6.3, 4.4 Hz, 1H), 2.82 – 2.75 (m, 1H), 2.61 – 2.53 (m, 2H), 2.24 (s, 2H), 2.02 (p, *J* = 6.8 Hz, 2H). [13]C NMR (150 MHz, DMSO-$d_6$) δ 170.87, 166.28, 159.41, 153.65, 148.14, 132.19, 130.56, 124.92, 121.19, 119.39, 116.81, 114.01, 112.57, 67.54, 50.45, 47.55, 47.47, 46.28, 46.02, 45.34, 44.84, 44.57, 42.06, 41.76, 41.66, 41.10, 29.16, 29.12, 24.91. HRMS (ESI) calculated for $C_{36}H_{40}N_8O_4$ $[M + H]^+$ 649.3245, found 649.3240.

1-(4-((2*S*,3*R*)-3-Methyloxirane-2-carbonyl)piperazin-1-yl)-4-(3-(6-(4-methylpiperazin-1-yl)-1*H*,3'*H*-[2,5'-bibenzo[*d*]imidazol]-2'-yl)phenoxy)butan-1-one (**H6**)

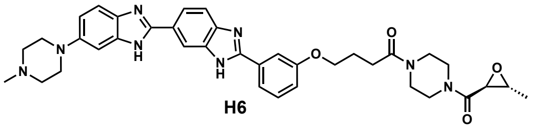

**H6**

Compound **H6** was prepared via a similar procedure to that of compound **H4.** [1]H NMR (600 MHz, DMSO-$d_6$) δ 9.99 (s, 2H), 8.41 (s, 1H), 7.99 (dd, *J* = 8.4, 1.8 Hz, 1H), 7.85 (d, *J* = 8.4 Hz, 1H), 7.77 (d, *J* = 8.0 Hz, 1H), 7.75 (s, 1H), 7.65 (d, *J* = 9.0 Hz, 1H), 7.45 (t, *J* = 7.9 Hz, 1H), 7.26 (dd, *J* = 9.0, 2.4 Hz, 1H), 7.17 (d, *J* = 2.4 Hz, 1H), 7.09 (dd, *J* = 8.2, 2.7 Hz, 1H), 4.07 (t, *J* = 6.4 Hz, 2H), 3.86 (d, *J* = 14.3 Hz, 2H), 3.60 – 3.30 (m, 11H), 3.25 – 3.11 (m, 2H), 3.05 – 2.95 (m, 3H), 2.84 (s, 3H), 2.51 (q, *J* = 7.0 Hz, 2H), 1.97 (t, *J* = 6.9 Hz, 2H), 1.23 (d, *J* = 5.2 Hz, 3H). [13]C NMR (150 MHz, DMSO-$d_6$) δ 170.87, 166.30, 159.50, 154.53, 150.04, 148.83, 130.85, 130.75, 122.40, 119.75, 117.71, 117.17, 114.99, 113.04, 99.86, 67.64, 54.11, 53.99, 53.83, 52.80, 46.96, 45.35,

44.86, 44.60, 42.57, 41.98, 41.67, 41.11, 29.09, 24.88, 17.45. HRMS (ESI) calculated for $C_{37}H_{42}N_8O_4$ $[M + H]^+$ 663.3402, found 663.3400.

1-((*S*)-3-Methyl-4-((*R*)-oxirane-2-carbonyl)piperazin-1-yl)-4-(3-(6-(4-methylpiperazin-1-yl)-1*H*,3'*H*-[2,5'-bibenzo[*d*]imidazol]-2'-yl)phenoxy)butan-1-one (**H7**)

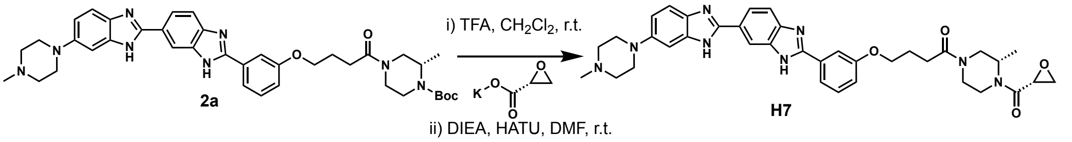


Intermediate **2a** was prepared following a procedure analogous to that used for intermediate **1a**. Specifically, **H** was coupled with *tert*-butyl-(*S*)-2-methylpiperazine-1-carboxylate by an amide coupling reaction. Intermediate **2a** was then converted to **H7** using a procedure analogous to that used for the synthesis of **H4**. $^1$H NMR (600 MHz, DMSO-$d_6$) δ 12.64 (brs, 2H), 8.33 (s, 1H), 8.03 (d, *J* = 8.4 Hz, 1H), 7.91 – 7.78 (m, 2H), 7.71 (d, *J* = 8.4 Hz, 1H), 7.53 – 7.40 (m, 2H), 7.08 (t, *J* = 5.5 Hz, 1H), 7.00 (s, 1H), 6.93 (d, *J* = 5.4 Hz, 1H), 4.58 – 4.06 (m, 5H), 4.02 – 3.68 (m, 3H), 3.61 – 3.16 (m, 6H), 3.12 (s, 3H), 2.99 – 2.54 (m, 6H), 2.24 (s, 3H), 2.02 (s, 2H), 1.37 – 0.93 (m, 3H). $^{13}$C NMR (150 MHz, DMSO-$d_6$) δ 171.35, 171.23, 166.20, 159.42, 153.39, 148.18, 131.96, 130.60, 125.09, 121.33, 119.40, 116.89, 114.10, 112.56, 67.57, 67.48, 50.44, 49.45, 48.79, 48.43, 46.27, 45.99, 45.65, 45.45, 45.20, 44.91, 41.75, 41.23, 36.88, 36.43, 28.97, 28.76, 24.93, 16.84, 16.47, 15.33, 14.90. HRMS (ESI) calculated for $C_{37}H_{42}N_8O_4$ $[M + H]^+$ 663.3402, found 663.3400.

1-((*S*)-3-Methyl-4-((*S*)-oxirane-2-carbonyl)piperazin-1-yl)-4-(3-(6-(4-methylpiperazin-1-yl)-1*H*,3'*H*-[2,5'-bibenzo[*d*]imidazol]-2'-yl)phenoxy)butan-1-one (**H8**)

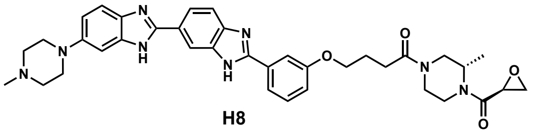

**H8**

Compound **H8** was prepared via a similar procedure to that of compound **H7**. $^1$H NMR (600 MHz, DMSO-$d_6$) δ 13.06 (s, 1H), 12.51 (s, 1H), 8.24 (s, 1H), 7.95 (d, *J* = 8.5 Hz,

1H), 7.73 (d, $J$ = 6.8 Hz, 2H), 7.63 (d, $J$ = 8.6 Hz, 1H), 7.40 (t, $J$ = 6.9 Hz, 2H), 7.05 – 6.98 (m, 2H), 6.87 (d, $J$ = 9.7 Hz, 1H), 4.53 – 3.99 (m, 6H), 3.91 – 3.32 (m, 6H), 3.06 (s, 3H), 2.95 – 2.56 (m, 5H), 2.54 – 2.46 (m, 2H), 2.18 (s, 3H), 1.97 (t, $J$ = 6.9 Hz, 2H), 1.04 (ddd, $J$ = 64.9, 47.9, 6.5 Hz, 3H). $^{13}$C NMR (150 MHz, DMSO-$d_6$) δ 171.38, 171.25, 166.47, 166.11, 159.42, 153.28, 148.22, 131.94, 130.60, 125.07, 121.30, 119.38, 116.88, 112.62, 112.57, 67.59, 67.50, 53.18, 50.45, 49.25, 48.72, 48.16, 47.74, 46.28, 46.01, 45.29, 44.87, 41.85, 37.02, 36.56, 28.98, 28.77, 24.96, 21.25, 21.18, 16.99, 16.60, 15.54, 15.11. HRMS (ESI) calculated for $C_{37}H_{42}N_8O_4$ [M + H]$^+$ 663.3402, found 663.3401.

($R$)-$N$-(4-(4-(3-(6-(4-Methylpiperazin-1-yl)-1$H$,3'$H$-[2,5'-bibenzo[$d$]imidazol]-2'-yl)phenoxy)butanamido)phenyl)oxirane-2-carboxamide (**H9**)

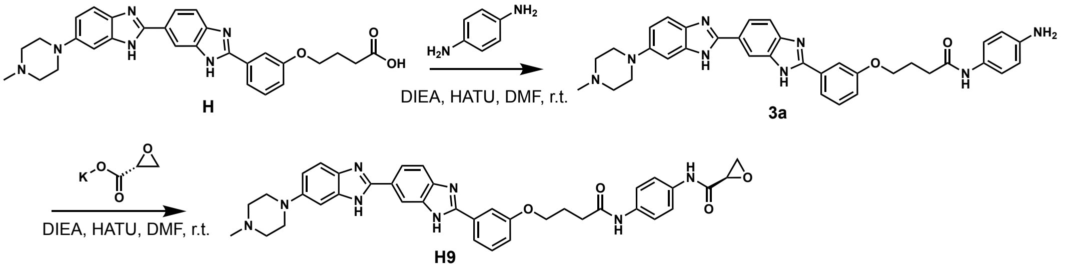


To a solution of compound **H** (50 mg, 0.10 mmol) in DMF (1 mL), was added benzene-1,4-diamine (32 mg, 0.30 mmol), DIEA (39 mg, 0.30 mmol), and HATU (46 mg, 0.12 mmol). The reaction mixture was stirred at room temperature for 3 h, diluted with $H_2O$ (10 mL), and extracted with EtOAc (10 mL). The organic layer was washed with brine, dried over anhydrous $Na_2SO_4$, filtered, and concentrated to afford the crude intermediate **3a** as a dark solid, which was used directly in the next step without further purification. LC-MS: calculated for $C_{35}H_{36}N_8O_2$ [M + H]$^+$ 601.3, found 601.3.

To a solution of crude intermediate **3a** (10 mg, ~0.02 mmol) in DMF (0.5 mL), was added potassium ($R$)-oxirane-2-carboxylate (4 mg, 0.03 mmol), DIEA (8 mg, 0.06 mmol), and HATU (15 mg, 0.04 mmol). The reaction mixture was stirred at room temperature for 3 h, before purifying by RP-HPLC (0.1% TFA in MeOH/$H_2O$) to afford compound **H9** (3 mg, ~20%). [1]H NMR (600 MHz, DMSO-$d_6$) δ 10.11 (s, 1H), 9.97 (s, 1H), 9.87 (brs, 1H), 8.46 (s, 1H), 8.04 (dd, *J* = 8.4, 1.8 Hz, 1H), 7.90 (d, *J* = 8.5 Hz, 1H), 7.84 – 7.79 (m, 2H), 7.70 (d, *J* = 8.9 Hz, 1H), 7.52 (t, *J* = 7.9 Hz, 4H), 7.29 (d, *J* = 9.0 Hz, 1H), 7.21 (d, *J* = 2.4 Hz, 1H), 7.15 (dd, *J* = 7.9, 3.0 Hz, 1H), 4.16 (t, *J* = 6.4 Hz, 2H), 3.92 (d, *J* = 13.3 Hz, 2H), 3.61 – 3.56 (m, 2H), 3.53 (dd, *J* = 4.3, 2.5 Hz, 1H), 3.24 (brs, 2H), 3.05 (t, *J* = 11.6 Hz, 2H), 2.96 (dd, *J* = 6.3, 4.3 Hz, 1H), 2.91 (s, 3H), 2.89 (dd, *J* = 6.4, 2.5 Hz, 1H), 2.54 (t, *J* = 7.4 Hz, 2H), 2.14 – 2.07 (m, 2H). [13]C NMR (150 MHz, DMSO-$d_6$) δ 170.89, 166.53, 159.52, 154.45, 150.28, 148.62, 135.75, 134.01, 130.85, 122.31, 120.53, 119.87, 113.00, 67.71, 52.85, 49.26, 47.05, 46.16, 42.57, 33.10, 25.14. HRMS (ESI) calculated for $C_{38}H_{38}N_8O_4$ $[M + H]^+$ 671.3089, found 671.3086.

(*S*)-*N*-(4-(4-(3-(6-(4-Methylpiperazin-1-yl)-1*H*,3'*H*-[2,5'-bibenzo[*d*]imidazol]-2'-yl)phenoxy)butanamido)phenyl)oxirane-2-carboxamide (**H10**)

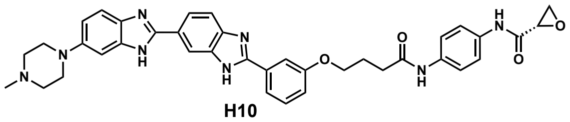

**H10**

Compound **H10** was prepared via a similar procedure to that of compound **H9**. [1]H NMR (600 MHz, DMSO-$d_6$) δ 10.12 (s, 1H), 9.97 (s, 1H), 9.88 (brs, 1H), 8.46 (s, 1H), 8.05 (dd, *J* = 8.4, 1.8 Hz, 1H), 7.90 (d, *J* = 8.4 Hz, 1H), 7.87 – 7.80 (m, 2H), 7.70 (d, *J* = 8.9 Hz, 1H), 7.55 (s, 4H), 7.29 (dd, *J* = 8.8, 2.0 Hz, 1H), 7.21 (d, *J* = 2.4 Hz, 1H), 7.15 (dd, *J* = 8.2, 1.6 Hz, 1H), 4.16 (t, *J* = 6.4 Hz, 2H), 3.95 – 3.89 (m, 2H), 3.62 – 3.56 (m, 2H), 3.53 (dd, *J* = 4.4, 2.5 Hz, 1H), 3.24 (brs, 2H), 3.09 – 3.01 (m, 2H), 2.98 – 2.94 (m, 1H), 2.91 (s, 3H), 2.89 – 2.87 (m, 1H), 2.54 (t, *J* = 7.3 Hz, 2H), 2.15 – 2.06 (m, 2H). [13]C NMR (150 MHz, DMSO-$d_6$) δ 170.89, 166.53, 159.52, 154.42, 150.26, 148.62, 135.76, 134.01, 130.84, 122.31, 120.53, 119.87, 119.74, 118.06, 116.88, 116.08, 115.08, 113.00, 67.71, 52.85, 49.26, 47.05, 46.16, 42.57, 33.10, 25.14. HRMS (ESI) calculated for $C_{38}H_{38}N_8O_4$ $[M + H]^+$ 671.3089, found 671.3088.

(*R*)-1-(4-(Aziridine-2-carbonyl)piperazin-1-yl)-4-(3-(6-(4-methylpiperazin-1-yl)-1*H*,3'*H*-

[2,5'-bibenzo[*d*]imidazol]-2'-yl)phenoxy)butan-1-one (**H11**)

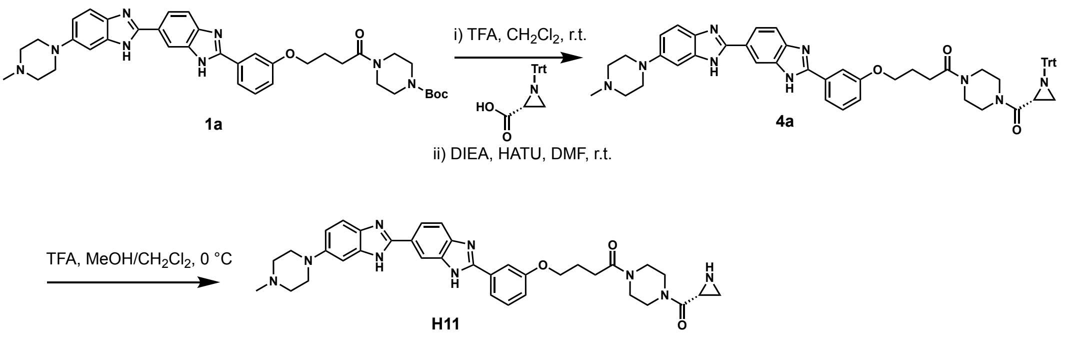

To a solution of intermediate **1a** (20 mg, 0.02 mmol) in $CH_2Cl_2$ (3 mL), was added TFA (2 mL). The reaction mixture was stirred at room temperature for 3 h and then concentrated to afford a yellow solid, which was subsequently dissolved in DMF (0.5 mL). (*R*)-1-Tritylaziridine-2-carboxylic acid (10 mg, 0.03 mmol), DIEA (39 mg, 0.30 mmol), and HATU (15 mg, 0.04 mmol) were then added. The reaction mixture was stirred at room temperature for 3 h, diluted with $H_2O$ (15 mL), and extracted with EtOAc (10 mL). The organic layer was washed with brine, dried over anhydrous $Na_2SO_4$, filtered, and concentrated to afford the crude intermediate **4a** as a yellow solid, which was used directly in the next step without further purification. LC-MS: calculated for $C_{55}H_{55}N_9O_3$ $[M + H]^+$ 890.4, found 890.0.

To a solution of intermediate **4a** (10 mg, ~0.01 mmol) in MeOH/$CH_2Cl_2$ (1:1, v/v, total 1mL), was added TFA (8 mg, 0.06 mmol) at 0 °C. The resulting mixture was stirred at 0 °C for 1 h and then quenched by DIEA (0.2 mL), concentrated and purified by RP-HPLC (MeOH/$H_2O$ containing 0.1% $NH_4HCO_3$) to afford compound **H11** (2 mg, ~31% over two steps). $^1$H NMR (600 MHz, DMSO-*d*6) δ 13.04 (brs, 1H), 12.69 – 12.37 (m, 1H), 8.31 – 8.13 (m, 1H), 7.97 (d, *J* = 37.1 Hz, 1H), 7.75 – 7.52 (m, 3H), 7.41 (t, *J* = 8.0 Hz, 2H), 7.03 (d, *J* = 8.3 Hz, 1H), 6.88 (d, *J* = 14.9 Hz, 2H), 4.07 (t, *J* = 6.5 Hz, 2H), 3.76 – 3.38 (m, 10H), 3.17 – 2.98 (m, 4H), 2.74 (s, 1H), 2.51 (d, *J* = 7.6 Hz, 2H), 2.18 (s, 3H), 1.97 (p, *J* = 6.9 Hz, 2H), 1.56 (s, 2H), 1.19 (d, *J* = 5.8 Hz, 1H). $^{13}$C NMR (150 MHz, DMSO-*d*6) δ 170.89, 169.95, 159.45, 148.36, 145.31, 138.58, 136.38, 131.63, 130.67, 119.37, 119.12, 117.02, 113.81, 112.62, 111.42, 105.51, 97.64, 67.60, 55.43,

55.35, 50.73, 50.27, 46.28, 45.34, 44.88, 44.74, 44.48, 42.57, 42.28, 41.66, 41.17, 29.13, 27.56, 26.43, 24.91, 21.17. HRMS (ESI) calculated for $C_{36}H_{41}N_9O_3$ [M + H]$^+$ 648.3405, found 648.3403.

(*S*)-1-(4-(Aziridine-2-carbonyl)piperazin-1-yl)-4-(3-(6-(4-methylpiperazin-1-yl)-1*H*,3'*H*-[2,5'-bibenzo[*d*]imidazol]-2'-yl)phenoxy)butan-1-one (**H12**)

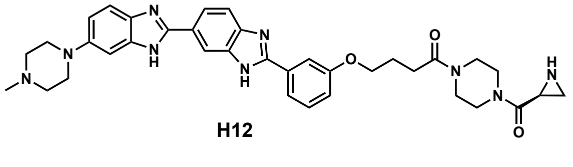

**H12**

Compound **H12** was prepared via a similar procedure to that of compound **H11**. $^1$H NMR (600 MHz, DMSO-$d_6$) δ 13.04 (brs, 1H), 12.60 (brs, 1H), 8.43 – 8.15 (m, 1H), 8.03 (s, 1H), 7.83 – 7.61 (m, 3H), 7.48 (t, *J* = 7.9 Hz, 2H), 7.13 – 7.06 (m, 1H), 6.99 – 6.88 (m, 2H), 4.14 (t, *J* = 6.4 Hz, 2H), 3.81 – 3.40 (m, 10H), 3.12 (d, *J* = 17.6 Hz, 4H), 2.81 (s, 1H), 2.58 (q, *J* = 7.5 Hz, 2H), 2.24 (s, 3H), 2.03 (p, *J* = 6.9 Hz, 2H), 1.71 – 1.55 (m, 2H), 1.26 (d, *J* = 5.6 Hz, 1H). $^{13}$C NMR (150 MHz, DMSO-$d_6$) δ 170.88, 169.95, 159.45, 152.97, 148.36, 138.60, 136.41, 131.64, 130.66, 119.37, 119.12, 116.99, 113.77, 112.62, 105.51, 97.64, 67.60, 55.35, 50.73, 50.27, 46.29, 45.34, 44.88, 44.74, 44.48, 42.57, 42.27, 41.66, 41.16, 29.13, 27.55, 26.43, 24.91, 21.17. HRMS (ESI) calculated for $C_{36}H_{41}N_9O_3$ [M + H]$^+$ 648.3405, found 648.3401.

1-(4-((2*S*,3*S*)-3-Methylaziridine-2-carbonyl)piperazin-1-yl)-4-(3-(6-(4-methylpiperazin-1-yl)-1*H*,3'*H*-[2,5'-bibenzo[*d*]imidazol]-2'-yl)phenoxy)butan-1-one (**H13**)

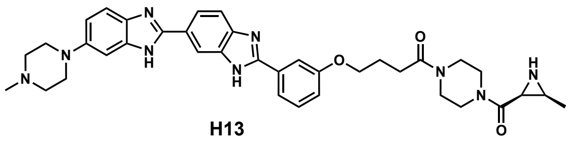

**H13**

Compound **H13** was prepared via a similar procedure to that of compound **H11**. $^1$H NMR (600 MHz, DMSO-$d_6$) δ 13.09 (s, 1H), 12.61 (s, 1H), 8.38 (s, 1H), 8.25 (s, 1H),

8.07 (d, $J$ = 10.0 Hz, 1H), 8.00 (d, $J$ = 10.0 Hz, 1H), 7.84 – 7.76 (m, 2H), 7.64 (d, $J$ = 8.4 Hz, 1H), 7.48 (td, $J$ = 7.9, 2.6 Hz, 1H), 7.10 (d, $J$ = 8.1 Hz, 1H), 6.95 (s, 1H), 4.13 (t, $J$ = 6.4 Hz, 2H), 3.74 – 3.39 (m, 10H), 3.17 – 3.12 (m, 4H), 2.76 (d, $J$ = 7.5 Hz, 1H), 2.57 (d, $J$ = 4.9 Hz, 5H), 2.30 (s, 3H), 2.12 (s, 1H), 2.03 (p, $J$ = 6.6 Hz, 2H), 1.03 (s, 3H). $^{13}$C NMR (150 MHz, DMSO-$d_6$) δ 170.89, 167.97, 159.45, 153.07, 152.84, 145.22, 144.42, 136.41, 135.76, 131.61, 130.67, 125.51, 124.98, 122.04, 121.03, 119.54, 119.35, 116.99, 112.64, 112.19, 67.58, 55.18, 50.18, 45.96, 45.59, 45.18, 44.82, 44.53, 41.91, 41.45, 35.18, 29.15, 24.92.

HRMS (ESI) calculated for $C_{37}H_{43}N_9O_3$ [M + H]$^+$ 662.3562, found 662.3558.

4-(3-(6-(4-Methylpiperazin-1-yl)-1*H*,3'*H*-[2,5'-bibenzo[*d*]imidazol]-2'-yl)phenoxy)-1-(4-pivaloylpiperazin-1-yl)butan-1-one (**H14**)

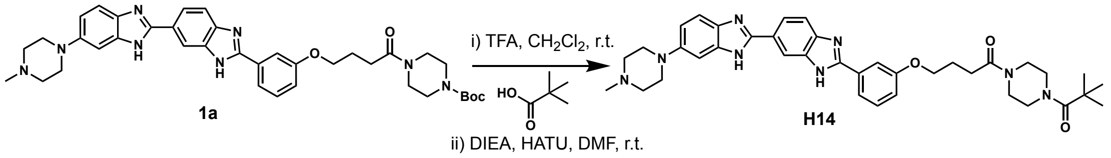


To a solution of intermediate **1a** (10 mg, 0.01 mmol) in $CH_2Cl_2$ (3 mL), was added TFA (2 mL). The reaction mixture was stirred at room temperature for 3 h and then concentrated to afford a yellow solid, which was subsequently dissolved in DMF (0.5 mL). Pivalic acid (2 mg, 0.02 mmol), DIEA (13 mg, 0.10 mmol) and HATU (7 mg, 0.02 mmol) were then added. The reaction mixture was stirred at room temperature for 3 h, and then purified by RP-HPLC (0.1% TFA in MeOH/$H_2O$) to afford compound **H14** (3 mg, 45% over two steps). $^1$H NMR (600 MHz, DMSO-$d_6$) δ 10.08 (s, 1H), 8.48 (s, 1H), 8.06 (dd, $J$ = 8.4, 1.8 Hz, 1H), 7.92 (d, $J$ = 8.5 Hz, 1H), 7.84 (d, $J$ = 8.2 Hz, 1H), 7.81 (d, $J$ = 1.0 Hz, 1H), 7.72 (d, $J$ = 9.0 Hz, 1H), 7.52 (t, $J$ = 8.0 Hz, 1H), 7.33 (dd, $J$ = 9.0, 2.3 Hz, 1H), 7.24 (d, $J$ = 2.4 Hz, 1H), 7.16 (dd, $J$ = 8.8, 2.1 Hz, 1H), 4.14 (t, $J$ = 6.4 Hz, 2H), 3.93 (d, $J$ = 11.3 Hz, 2H), 3.63 – 3.50 (m, 6H), 3.47 (dd, $J$ = 6.6, 3.8 Hz, 4H), 3.24 (brs, 2H), 3.07 (brs, 2H), 2.91 (s, 3H), 2.55 (t, $J$ = 7.2 Hz, 2H), 2.03 (p, $J$ = 6.9 Hz, 2H), 1.18 (s, 9H). $^{13}$C NMR (150 MHz, DMSO-$d_6$) δ 175.72, 170.82, 159.50, 154.54, 149.99, 148.87, 130.85, 130.71, 122.42, 119.76, 117.70, 117.23, 114.96, 113.05, 99.83, 67.63,

52.79, 46.94, 45.34, 44.98, 42.56, 41.67, 38.54, 29.10, 28.46, 24.89. HRMS (ESI) calculated for $C_{38}H_{46}N_8O_3$ [M + H]+ 663.3766, found 663.3764. HPLC purity 98.01%.

(*R*)-*N*-(2-(7,8-Dimethyl-2,4-dioxo-3,4-dihydrobenzo[*g*]pteridin-10(2*H*)-yl)ethyl)-*N*-methyloxirane-2-carboxamide (**C1**)

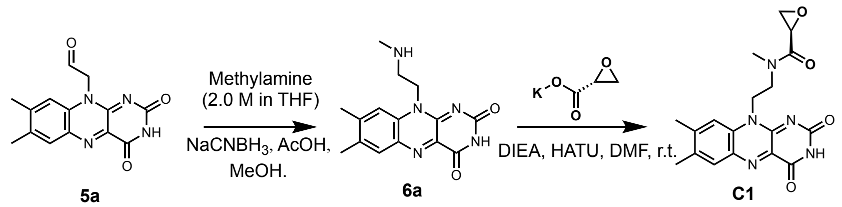


Intermediate **5a** was prepared according to a reported procedure.[3] To a solution of **5a** (100 mg, 0.35 mmol) in anhydrous MeOH (20 mL), was added methylamine solution (2.0 M in THF, 0.3 mL) and acetic acid (AcOH, 2 mL). The resulting mixture was stirred at 40 °C in the dark for 5 h and cooled to room temperature. Sodium cyanoborohydride ($NaCNBH_3$, 44 mg, 0.70 mmol) was then added into the reaction, which was stirred at room temperature for additional 12 h. The reaction was quenched by water (0.2 mL), and then was concentrated *in vacuo* to yield a yellow residue, which was purified by chromatography on a silica gel column (0.1% triethylamine in $CH_2Cl_2$/MeOH) to give yellow solid **6a** (50 mg, 47%). $^1H$ NMR (400 MHz, DMSO-$d_6$) δ 7.90 (s, 1H), 7.85 (s, 1H), 4.70 (t, *J* = 6.8 Hz, 2H), 2.92 (t, *J* = 6.6 Hz, 2H), 2.50 (s, 3H), 2.40 (s, 3H), 2.36 (s, 3H). LC-MS: calculated for $C_{15}H_{17}N_5O_2$ [M + H]+ 300.1, found 300.1.

To a solution of **6a** (10 mg, 0.03 mmol) in DMF (0.5 mL), was added potassium (*R*)-oxirane-2-carboxylate (6 mg, 0.04 mmol), DIEA (12 mg, 0.09 mmol), and HATU (19 mg, 0.05 mmol). The reaction mixture was stirred at room temperature for 3 h and then purified by RP-HPLC (0.1% (v/v) TFA in MeOH/$H_2O$) to afford compound **C1** (6 mg, 54%) as a *trans*/*cis* mixture (*trans*/*cis* = 2/1), yellow solid. $^1H$ NMR (*trans*, 600 MHz, DMSO-$d_6$) δ 11.36 (s, 1H), 7.89 (s, 1H), 7.85 (s, 1H), 4.79 (t, *J* = 6.0 Hz, 2H), 3.89 (t, *J* = 6.8 Hz, 1H), 3.81 (t, *J* = 6.2 Hz, 1H), 3.67 (t, *J* = 5.8 Hz, 1H), 3.63 (dd, *J* = 4.4, 2.6 Hz, 1H), 3.17 (s, 3H), 2.77 (dd, *J* = 6.5, 4.4 Hz, 1H), 2.51 (s, 3H), 2.40 (s, 3H), 2.27

(dd, *J* = 6.5, 2.6 Hz, 1H). $^{13}$C NMR (*trans*, 150 MHz, DMSO-$d_6$) δ 168.42, 160.33, 155.98, 151.02, 147.01, 137.22, 136.29, 134.25, 131.69, 131.53, 116.41, 47.16, 45.62, 45.47, 42.02, 35.60, 21.19, 19.22. HRMS (ESI) calculated for $C_{18}H_{19}N_5O_4$ [M + H]$^+$ 370.1510, found 370.1507. HPLC purity 98.36%.

(*S*)-*N*-(2-(7,8-Dimethyl-2,4-dioxo-3,4-dihydrobenzo[*g*]pteridin-10(2*H*)-yl)ethyl)-*N*-methyloxirane-2-carboxamide (**C2**)

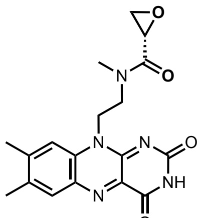

**C2**

Compound **C2** was prepared via a similar procedure to that of **C1** as a *trans*/*cis* mixture (*trans*/*cis* = 2/1), yellow solid. $^1$H NMR (*trans*, 600 MHz, DMSO-$d_6$) δ 11.29 (s, 1H), 7.83 (s, 1H), 7.78 (s, 1H), 4.73 (t, *J* = 6.1 Hz, 2H), 3.82 (t, *J* = 6.8 Hz, 1H), 3.76 – 3.73 (m, 1H), 3.61 (t, *J* = 5.8 Hz, 1H), 3.56 (dd, *J* = 4.4, 2.5 Hz, 1H), 3.10 (s, 3H), 2.71 (dd, *J* = 6.5, 4.4 Hz, 1H), 2.44 (s, 3H), 2.34 (s, 3H), 2.20 (dd, *J* = 6.5, 2.6 Hz, 1H). $^{13}$C NMR (*trans*, 150 MHz, DMSO-$d_6$) δ 168.41, 160.34, 155.99, 151.02, 147.01, 137.23, 136.28, 134.25, 131.69, 131.53, 116.41, 47.16, 45.62, 45.47, 42.03, 35.60, 21.19, 19.22. HRMS (ESI) calculated for $C_{18}H_{19}N_5O_4$ [M + H]$^+$ 370.1510, found 370.1509. HPLC purity 97.80%.

(*R*)-*N*-(2-(7,8-Dimethyl-2,4-dioxo-3,4-dihydrobenzo[*g*]pteridin-10(2*H*)-yl)ethyl)-*N*-methylaziridine-2-carboxamide (**C3**)

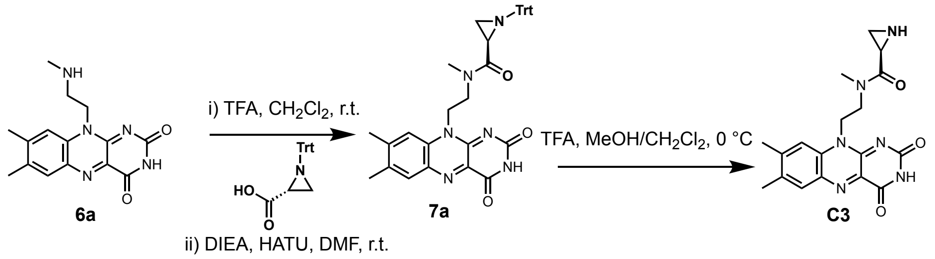

To a solution of intermediate **6a** (20 mg, 0.06 mmol) in DMF (1 mL), was added (*R*)-1-tritylaziridine-2-carboxylic acid (20 mg, 0.06 mmol), DIEA (15 mg, 0.12 mmol), and HATU (38 mg, 0.10 mmol). The reaction mixture was stirred at room temperature for 3 h, diluted with $H_2O$ (15 mL), and extracted with EtOAc (10 mL). The organic layer was washed with brine, dried over anhydrous $Na_2SO_4$, filtered, and concentrated to afford the crude intermediate **7a** as a yellow solid, which was used directly in the next step without further purification. LC-MS: calculated for $C_{37}H_{34}N_6O_3$ $[M + H]^+$ 611.3, found 611.3.

To a solution of intermediate **7a** (13 mg, ~0.02 mmol) in MeOH/$CH_2Cl_2$ (1:1, v/v, total 1mL), was added TFA (11 mg, 0.10 mmol) at 0 °C. The resulting mixture was stirred at 0 °C for 2 h and then quenched by DIEA (0.2 mL), concentrated and purified by RP-HPLC (MeOH/$H_2O$ containing 0.1% $NH_4HCO_3$) to afford **C3** (4 mg, ~50%) as a *trans*/*cis* mixture (*trans*/*cis* = 3/1), yellow solid. $^1$H NMR (*trans*, 600 MHz, DMSO-$d_6$) δ 11.37 (s, 1H), 7.90 (s, 1H), 7.84 (s, 1H), 5.04 – 4.84 (m, 2H), 4.01 – 3.93 (m, 1H), 3.60 (dt, *J* = 14.0, 5.1 Hz, 1H), 3.20 (s, 3H), 2.51 (s, 3H), 2.41 (s, 3H), 1.60 – 1.40 (m, 1H), 1.04 – 0.81 (m, 2H). $^{13}$C NMR (*trans*, 150 MHz, DMSO-$d_6$) δ 172.09, 160.34, 155.95, 151.01, 147.01, 137.13, 136.28, 134.26, 131.80, 131.51, 116.40, 45.81, 42.18, 35.67, 27.37, 26.02, 21.20, 19.21. HRMS (ESI) calculated for $C_{18}H_{20}N_6O_3$ $[M + H]^+$ 369.1664, found 369.1667. HPLC purity 98.12%.

(*S*)-*N*-(2-(7,8-dimethyl-2,4-dioxo-3,4-dihydrobenzo[*g*]pteridin-10(2*H*)-yl)ethyl)-*N*-methylaziridine-2-carboxamide (**C4**)

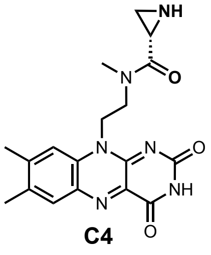


**C4**

Compound **C4** was prepared via a similar procedure to that of compound **C3** as a *trans*/*cis* mixture (*trans*/*cis* = 3/1), yellow solid. $^1$H NMR (*trans*, 600 MHz, DMSO-$d_6$) δ

11.36 (s, 1H), 7.88 (s, 1H), 7.83 (s, 1H), 4.99 – 4.83 (m, 2H), 3.99 – 3.93 (m, 1H), 3.59 (dd, *J* = 14.1, 5.1 Hz, 1H), 3.19 (s, 3H), 2.50 (s, 3H), 2.40 (s, 3H), 1.49 – 1.43 (m, 1H), 1.03 – 0.90 (m, 2H). $^{13}$C NMR (*trans,* 150 MHz, DMSO-$d_6$) δ 172.09, 160.34, 155.95, 151.01, 147.01, 137.13, 136.28, 134.25, 131.79, 131.51, 116.40, 45.80, 42.18, 35.67, 27.37, 26.02, 21.20, 19.21. HRMS (ESI) calculated for $C_{18}H_{20}N_6O_3$ [M + H]$^+$ 369.1664, found 369.1666. HPLC purity 97.96%.

**NMR spectra of final compounds**

Compound **P1**, [1]H NMR, DMSO-$d_6$, 600 MHz.

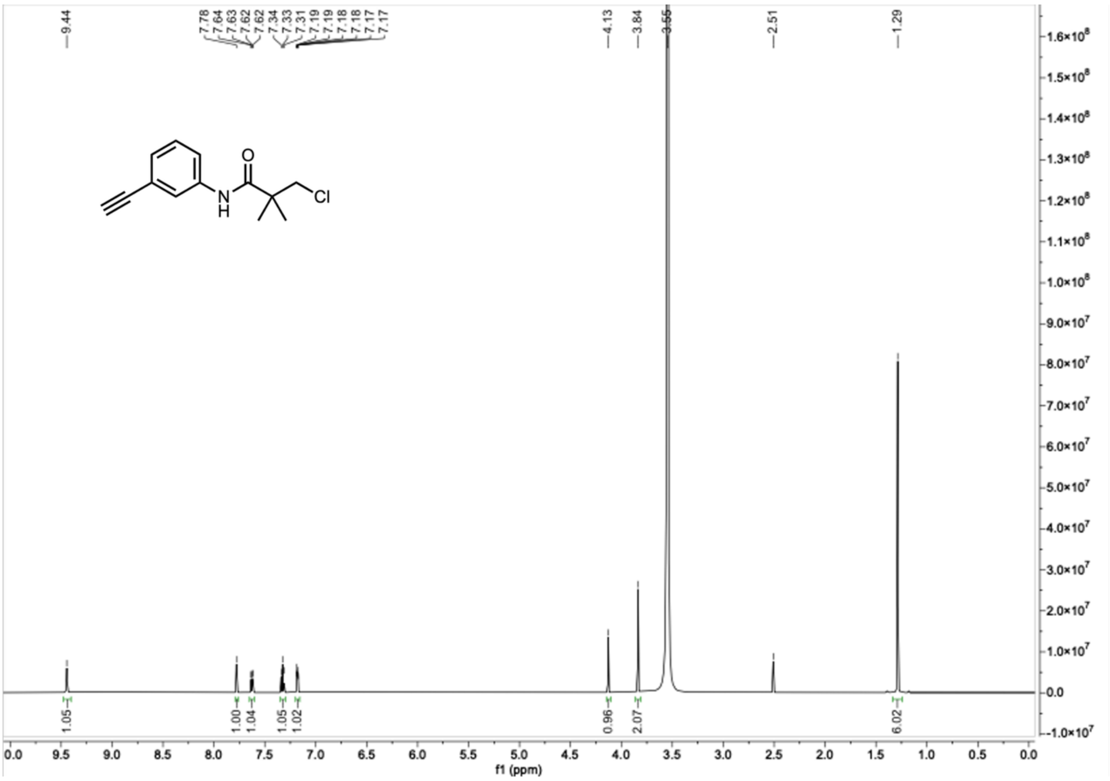


Compound **P1**, [13]C NMR, DMSO-$d_6$, 150 MHz.

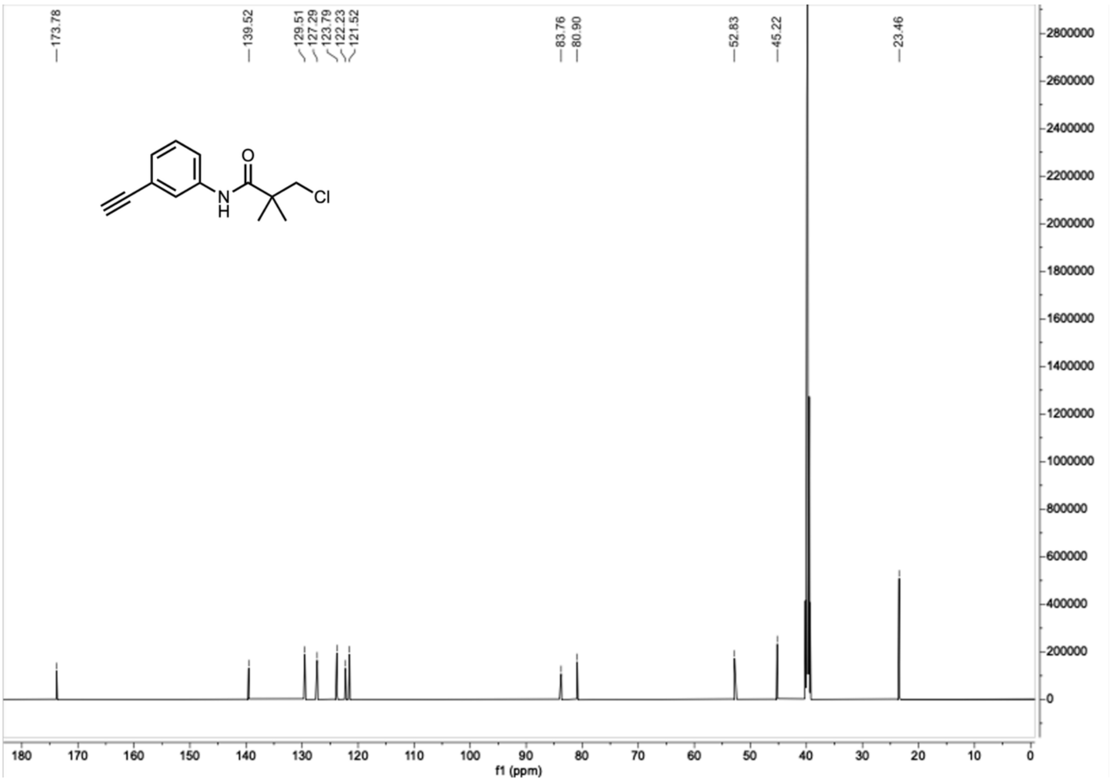

Compound **P2**, $^{1}H$ NMR, DMSO-$d_6$, 600 MHz.

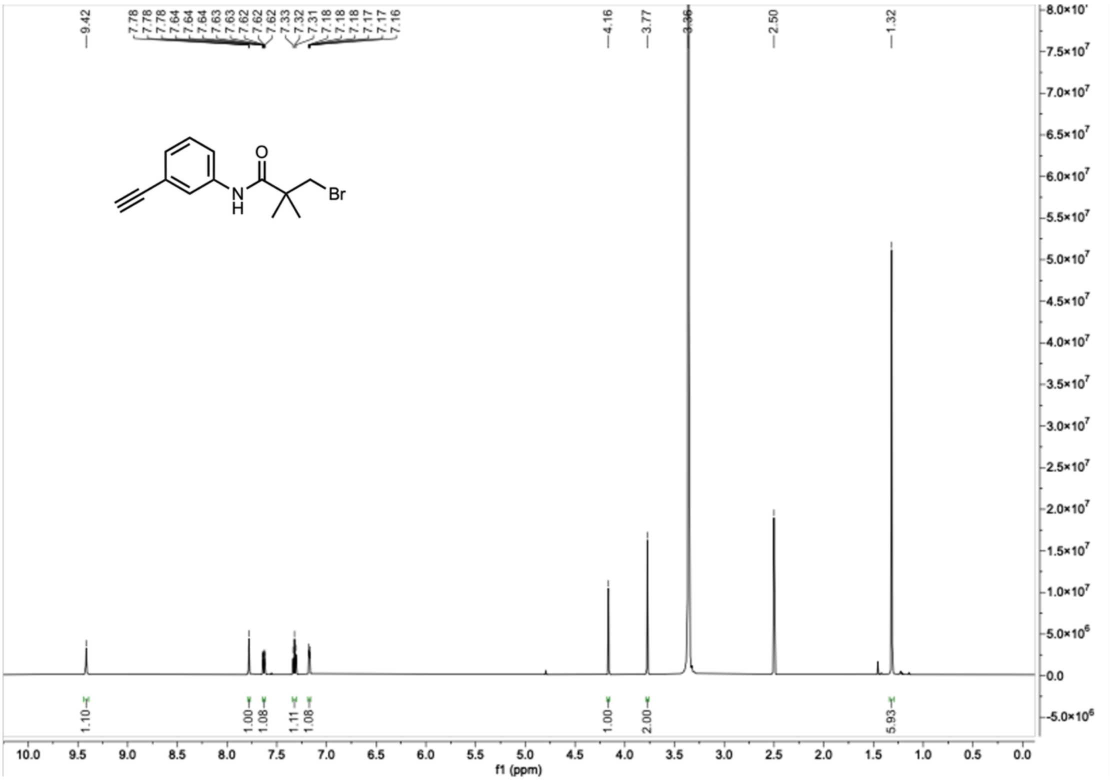

Compound **P2**, $^{13}C$ NMR, DMSO-$d_6$, 150 MHz.

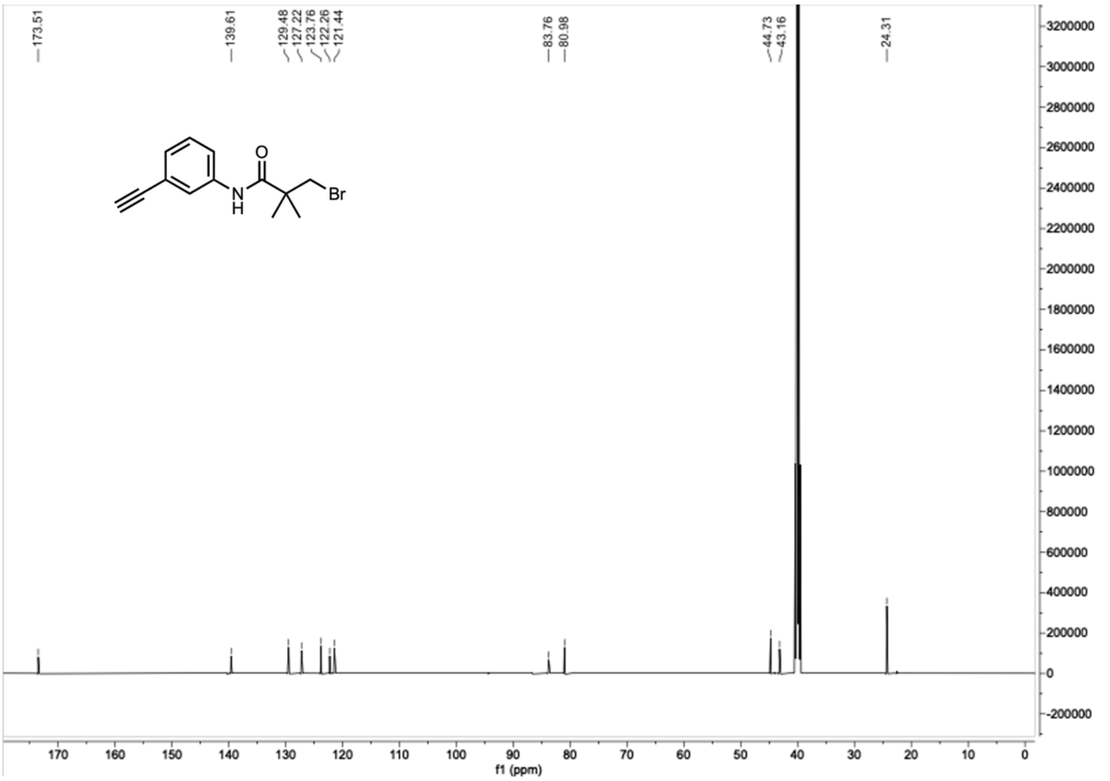

Compound **P3**, $^{1}H$ NMR, DMSO-$d_6$, 600 MHz.

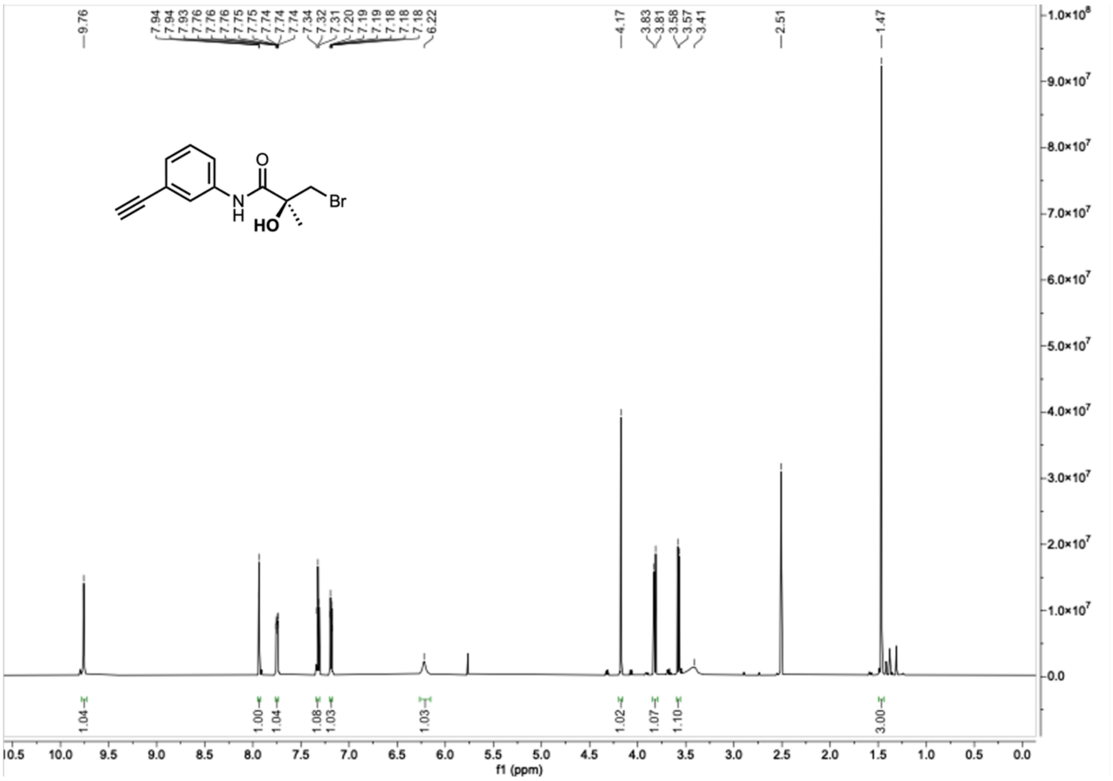

Compound **P3**, $^{13}C$ NMR, DMSO-$d_6$, 150 MHz.

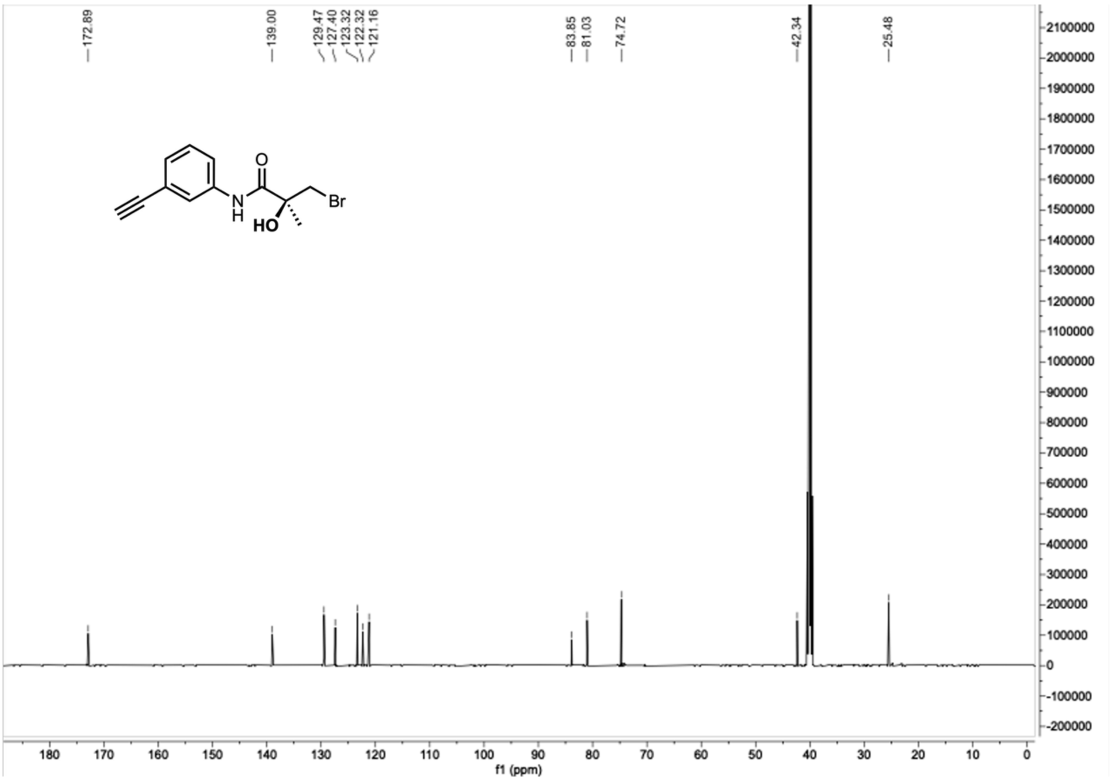

Compound **P4**, $^{1}H$ NMR, DMSO-$d_6$, 600 MHz.

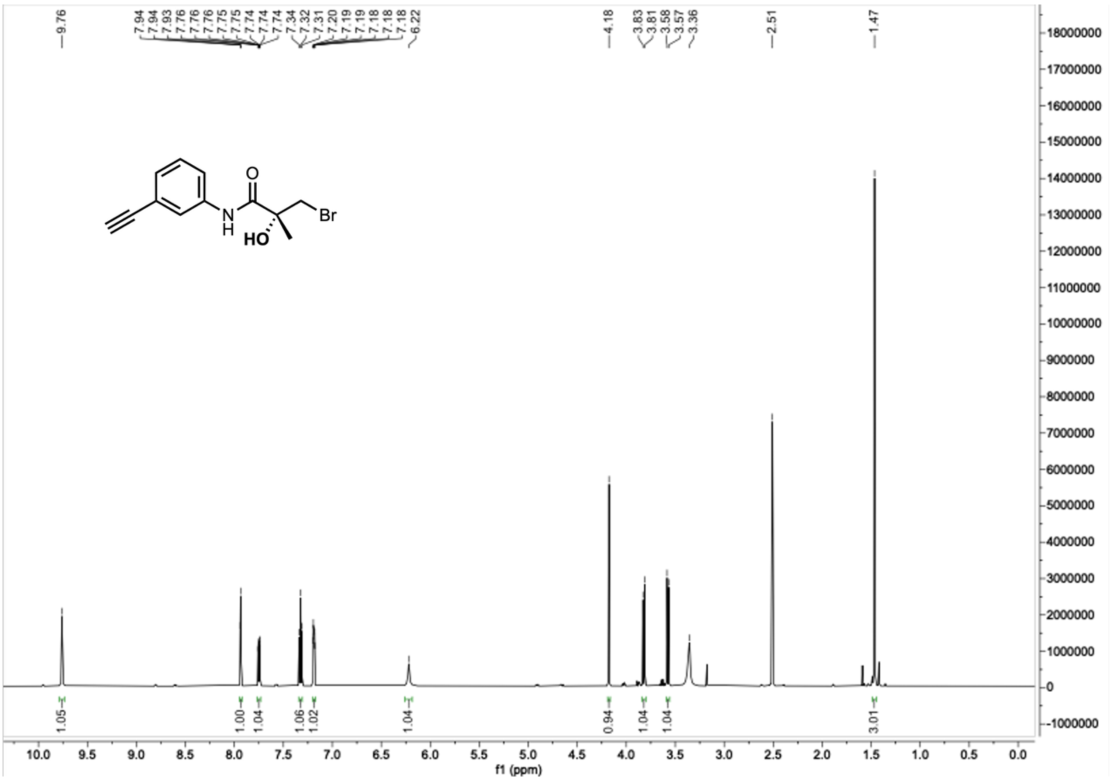

Compound **P4**, $^{13}C$ NMR, DMSO-$d_6$, 150 MHz.

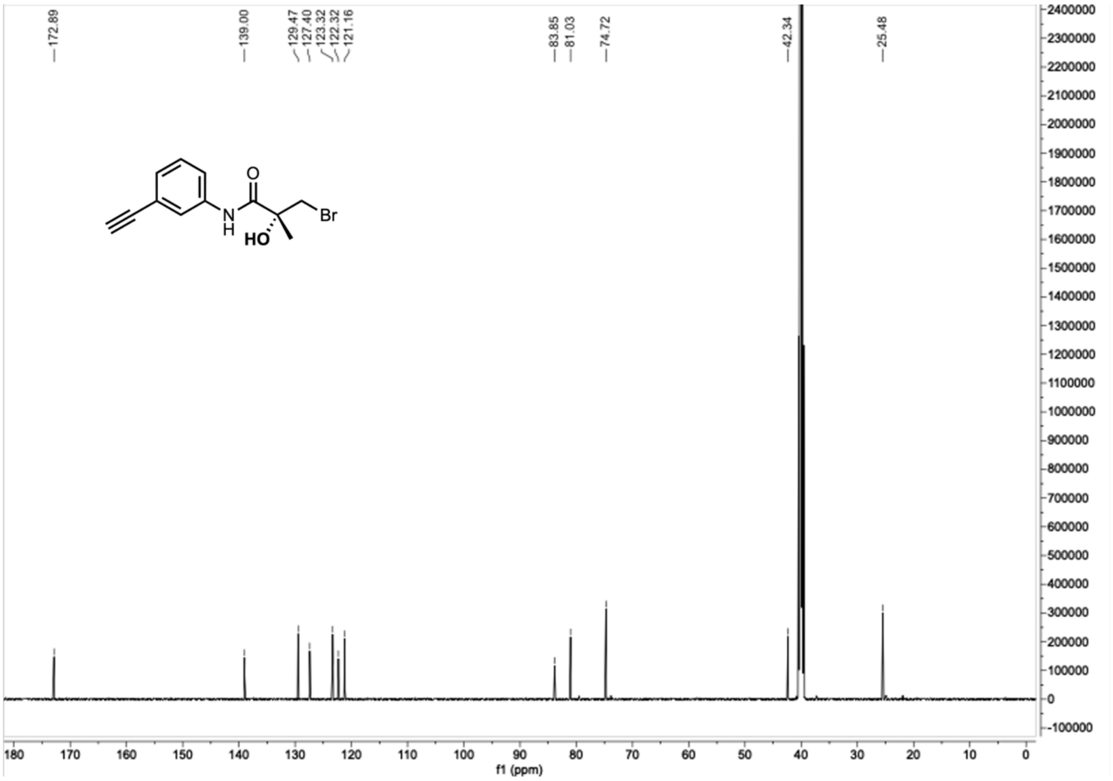

Compound **P3'**, $^{1}$H NMR, DMSO-$d_6$, 600 MHz.

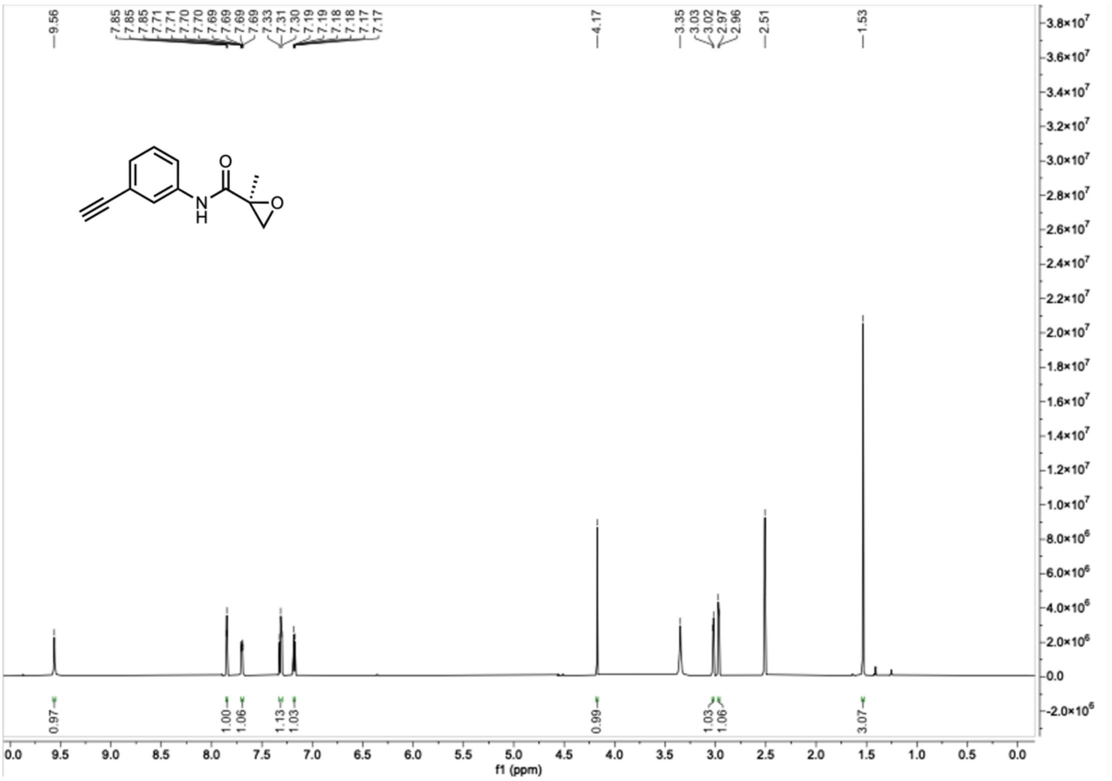

Compound **P3'**, $^{13}$C NMR, DMSO-$d_6$, 150 MHz.

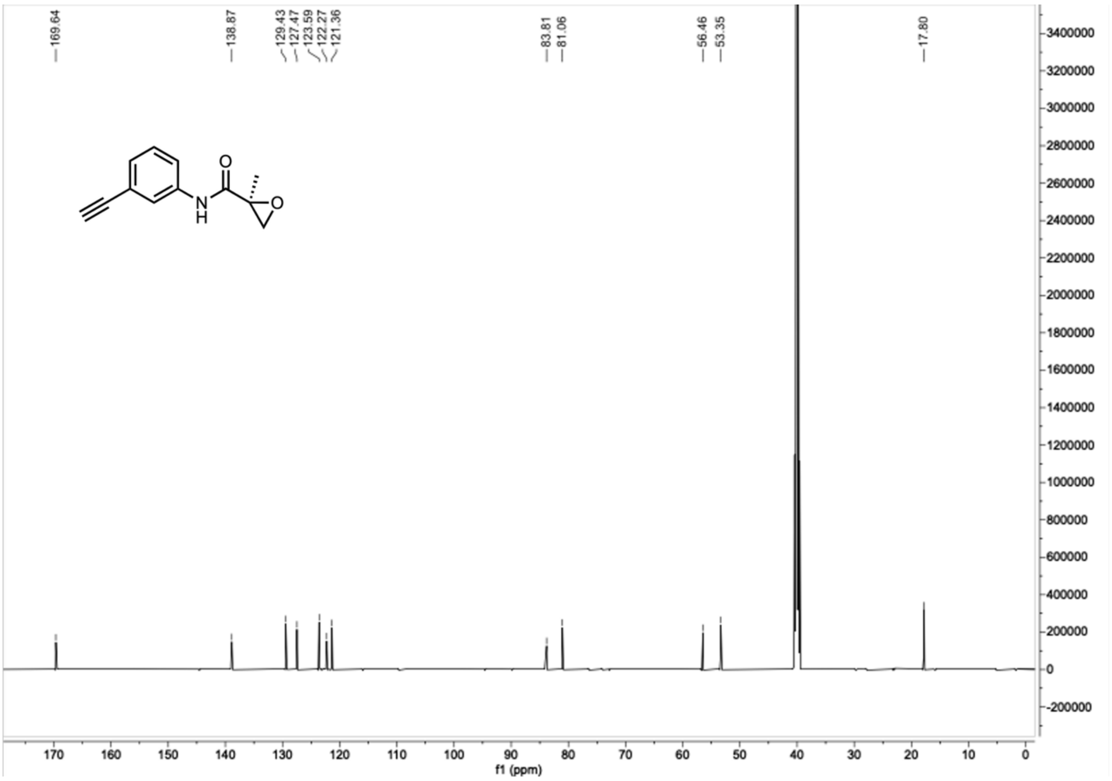

Compound **H1**, $^{1}H$ NMR, DMSO-$d_6$, 600 MHz.

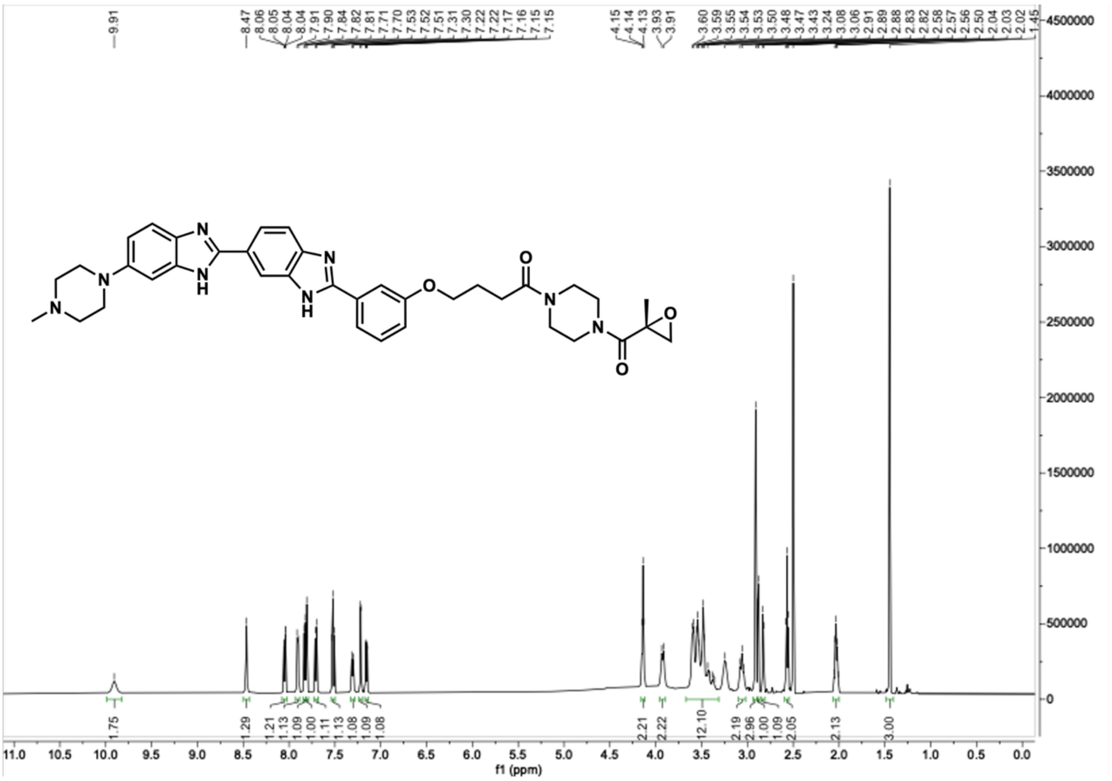

Compound **H1**, $^{13}C$ NMR, DMSO-$d_6$, 150 MHz.

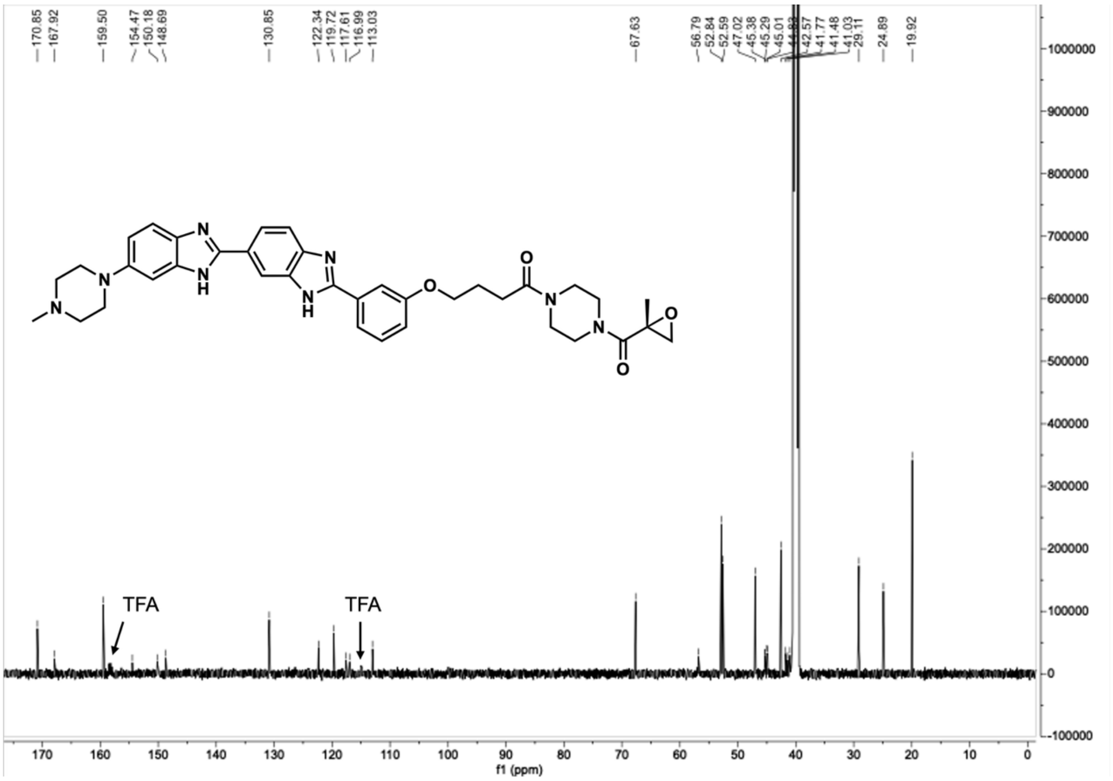

Compound **H2**, $^1$H NMR, DMSO-$d_6$, 600 MHz.

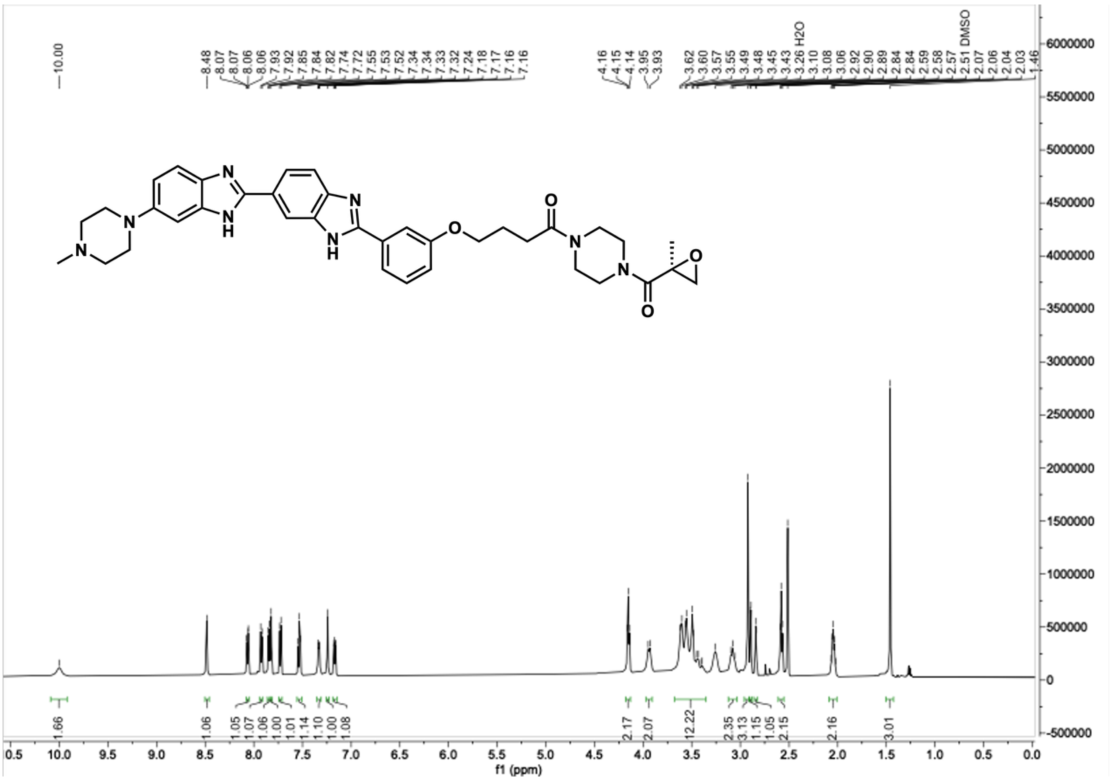

Compound **H2**, $^{13}$C NMR, DMSO-$d_6$, 150 MHz.

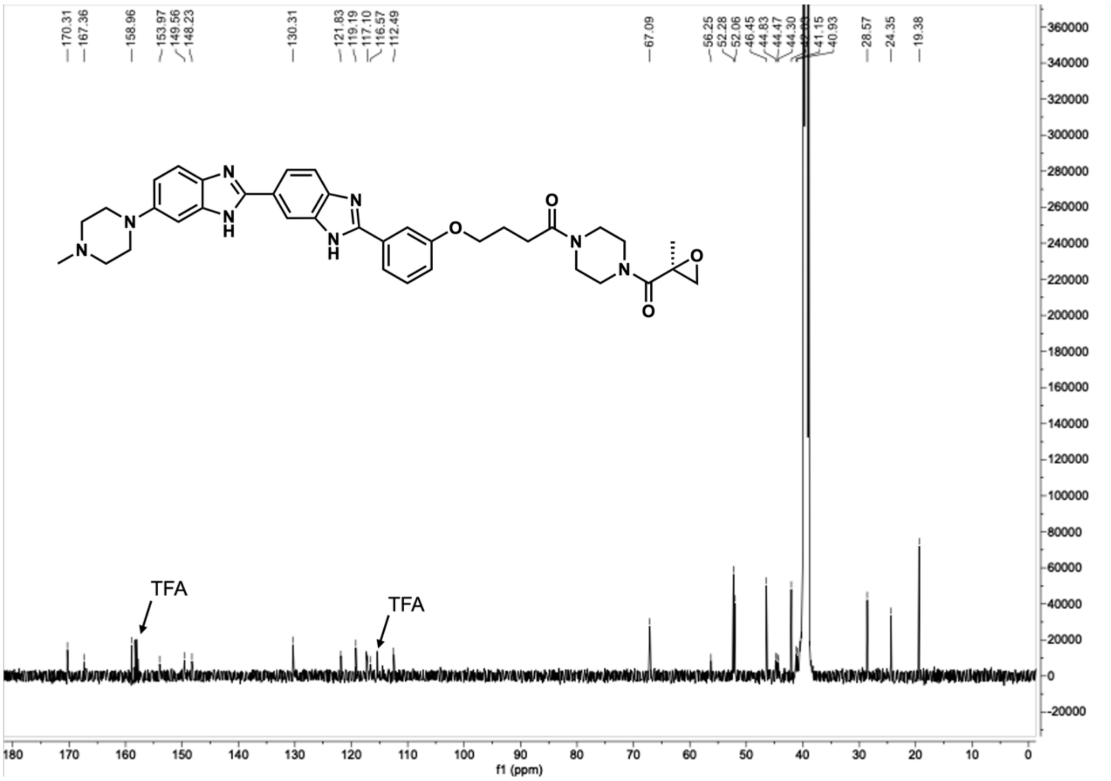

Compound **H3**, $^{1}$H NMR, DMSO-$d_6$, 600 MHz.

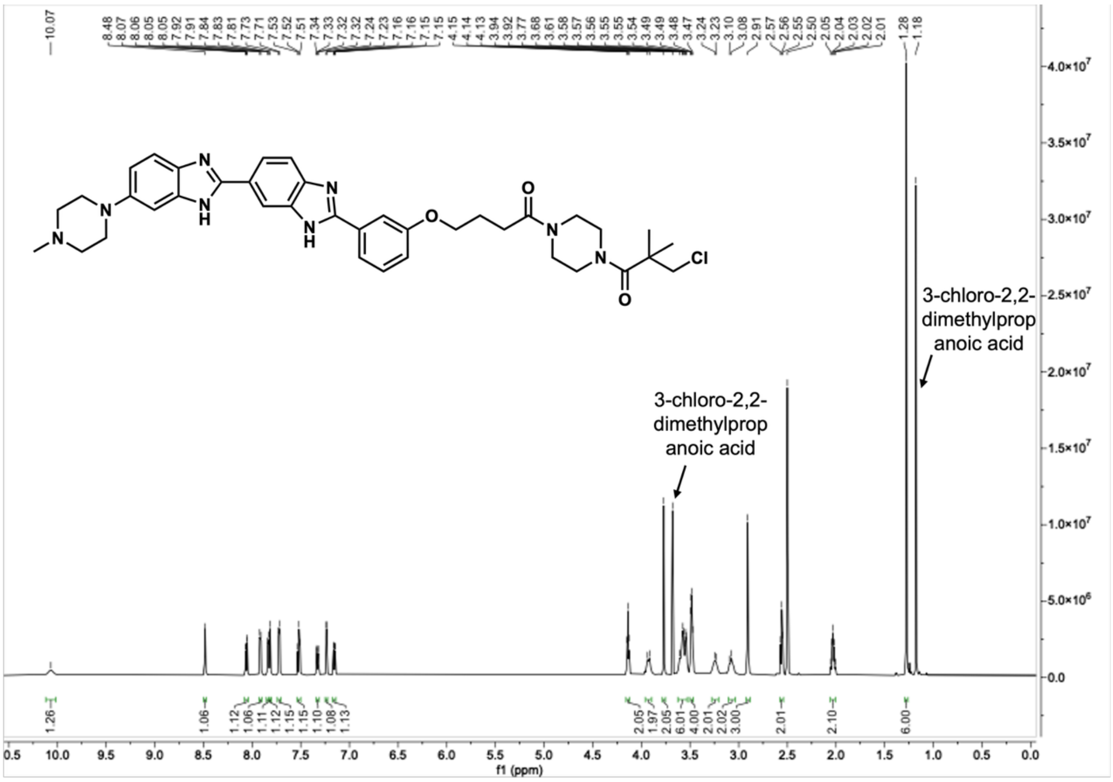


Compound **H3**, $^{13}$C NMR, DMSO-$d_6$, 150 MHz.

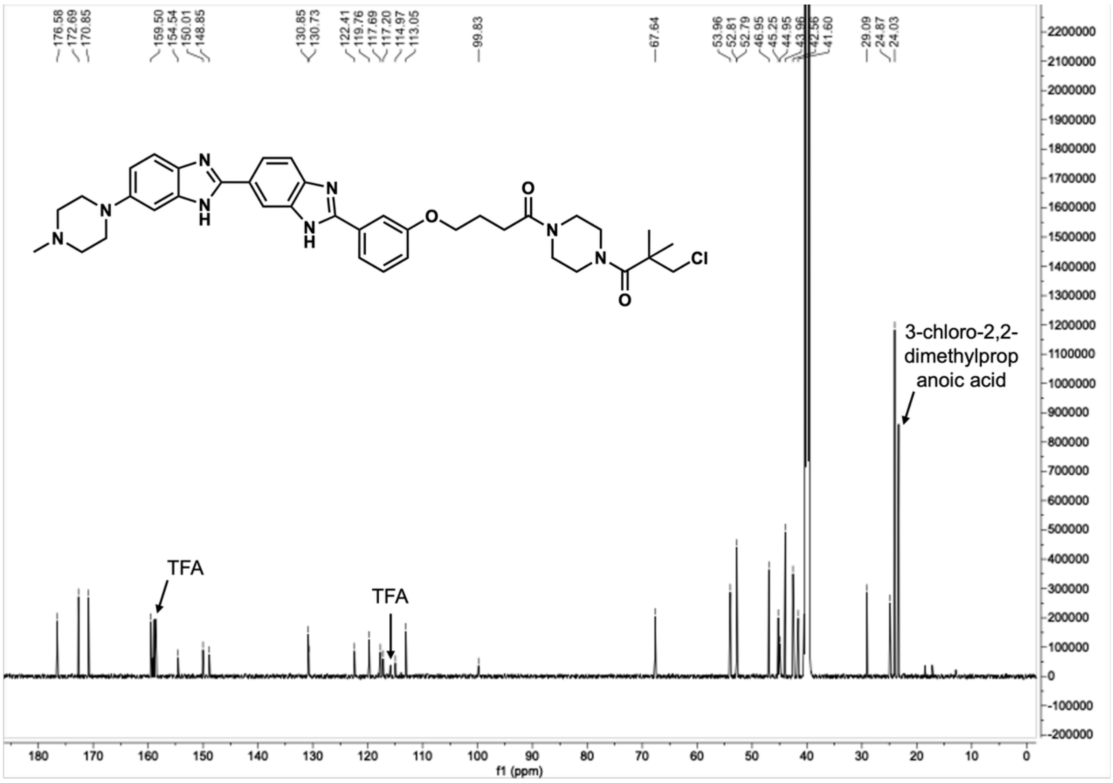

Compound **P5**, $^{1}$H NMR, $CDCl_3$, 600 MHz.

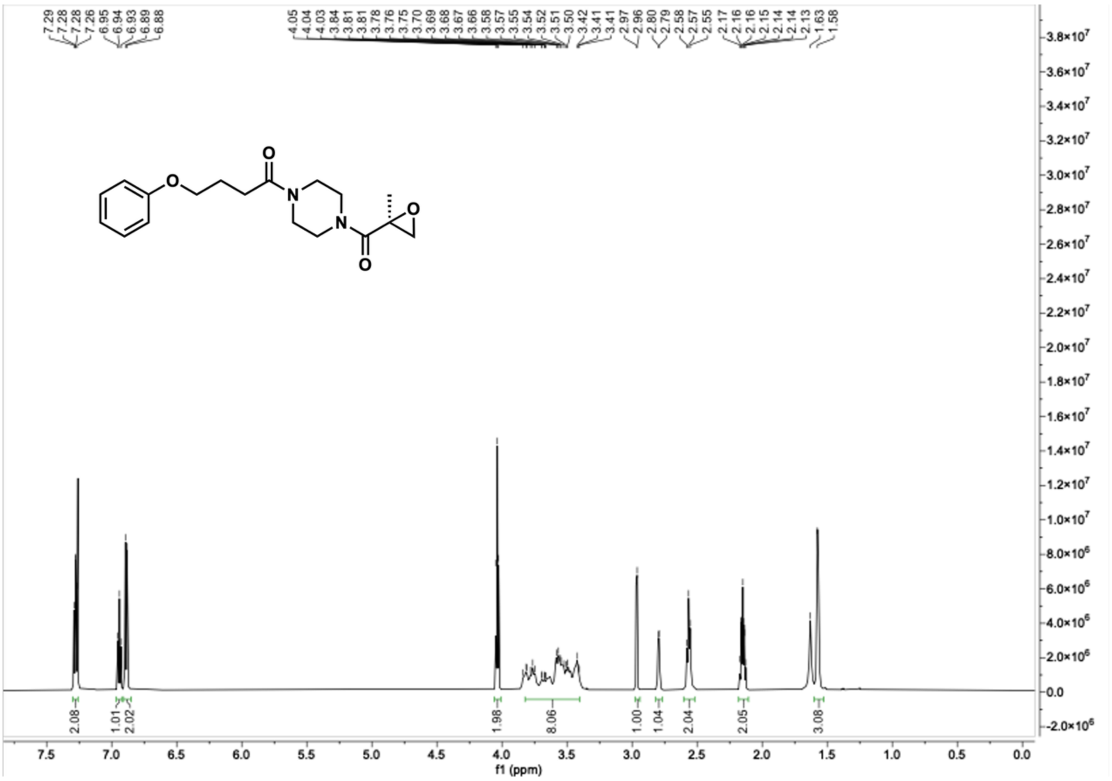

Compound **P5**, $^{13}$C NMR, $CDCl_3$, 150 MHz.

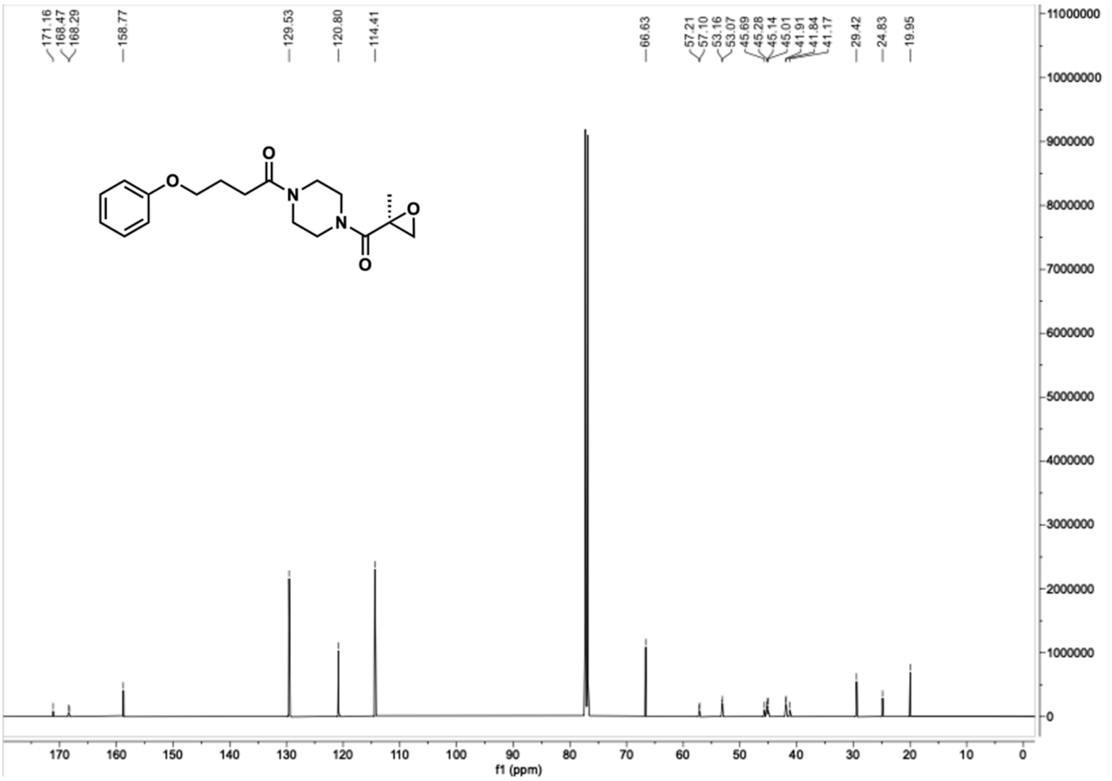

Compound **H4**, $^{1}$H NMR, DMSO-$d_6$, 600 MHz.

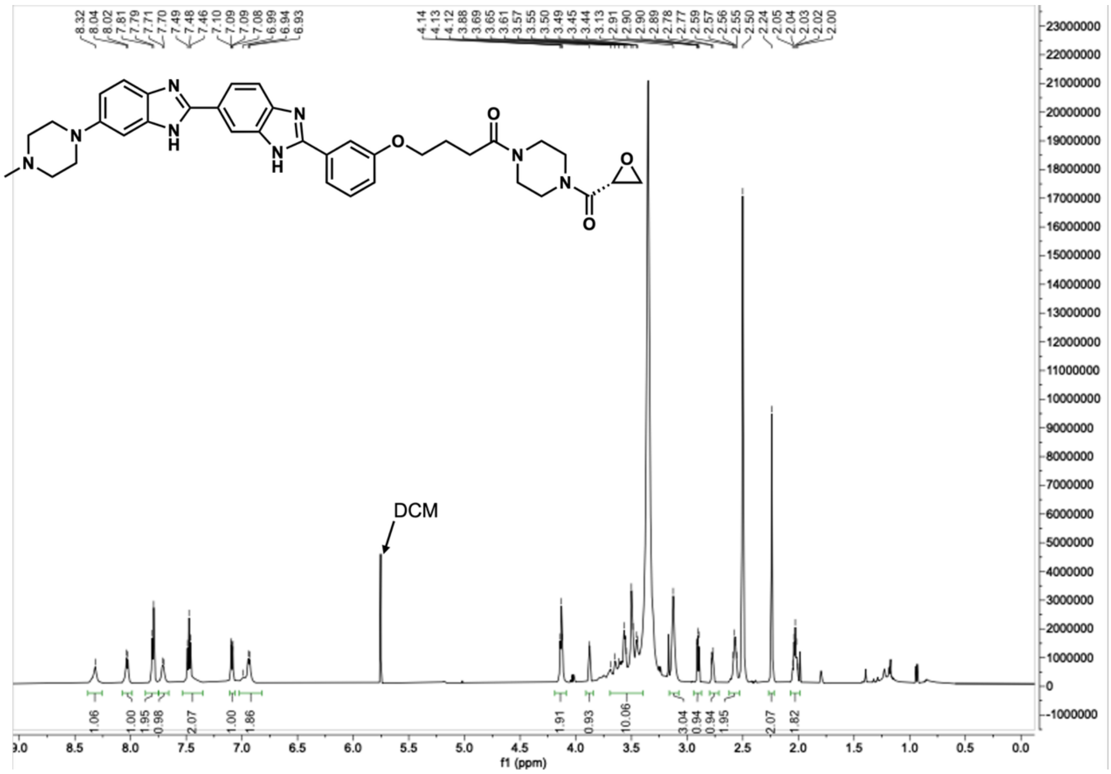

Compound **H4**, $^{13}$C NMR, DMSO-$d_6$, 150 MHz.

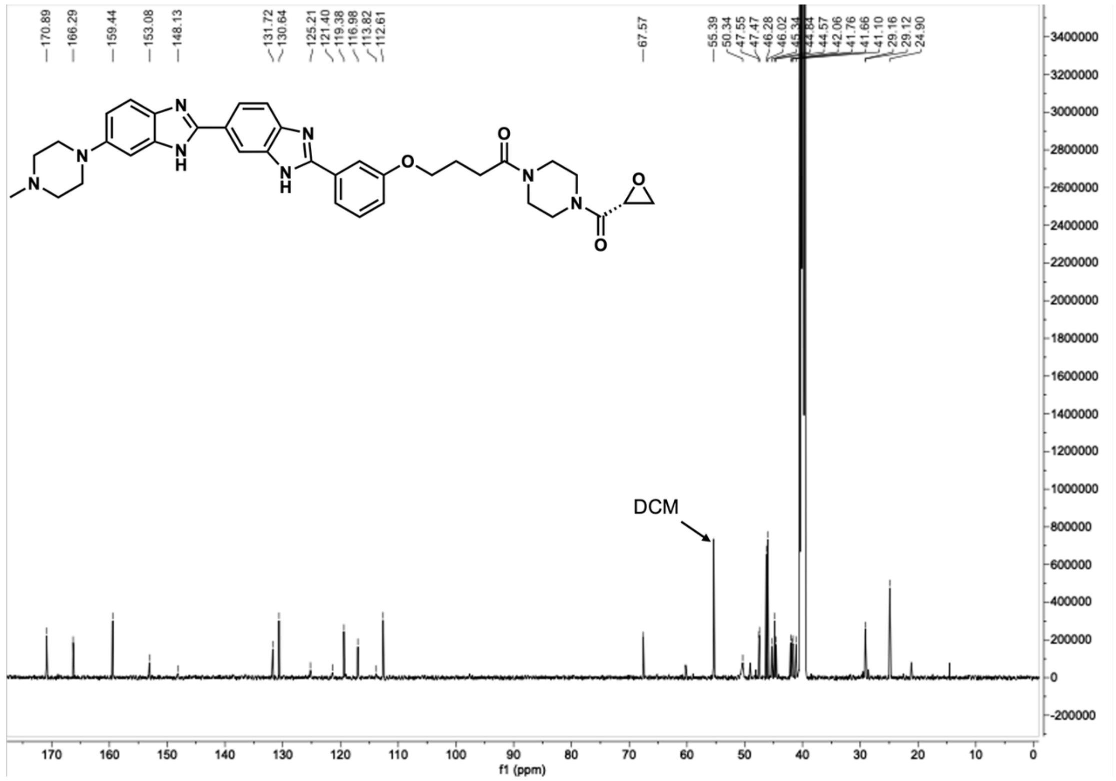

Compound **H5**, $^{1}H$ NMR, DMSO-$d_6$, 600 MHz.

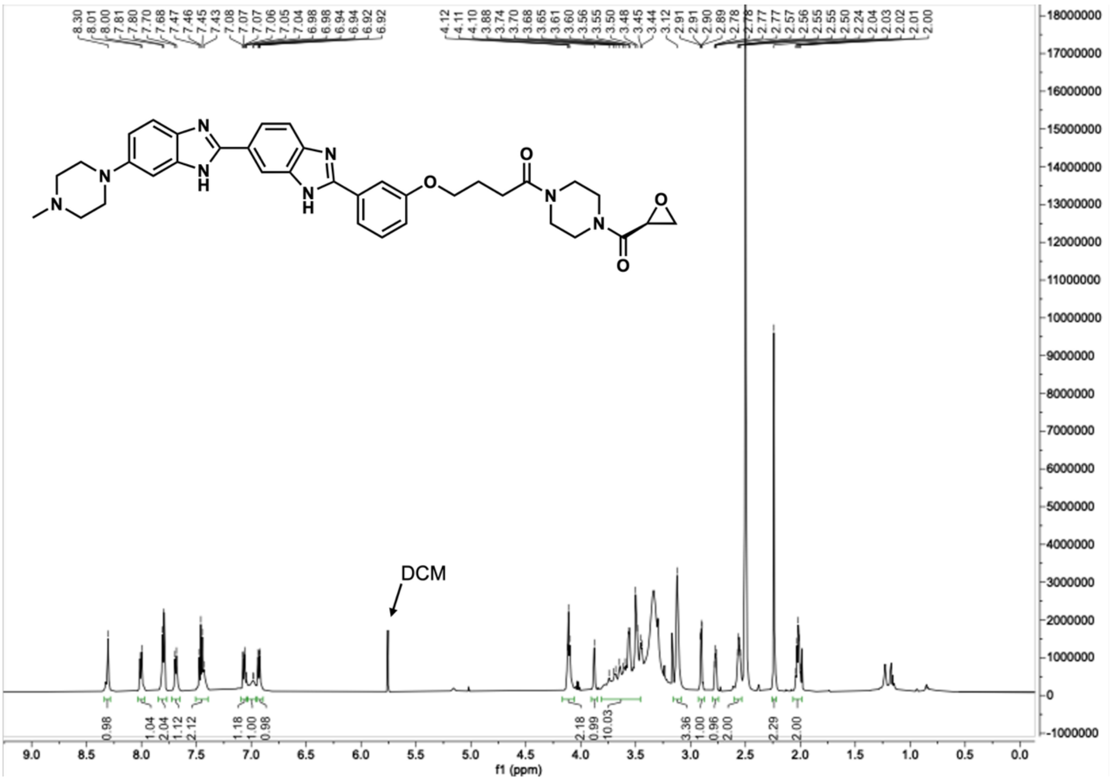


Compound **H5**, $^{13}C$ NMR, DMSO-$d_6$, 150 MHz.

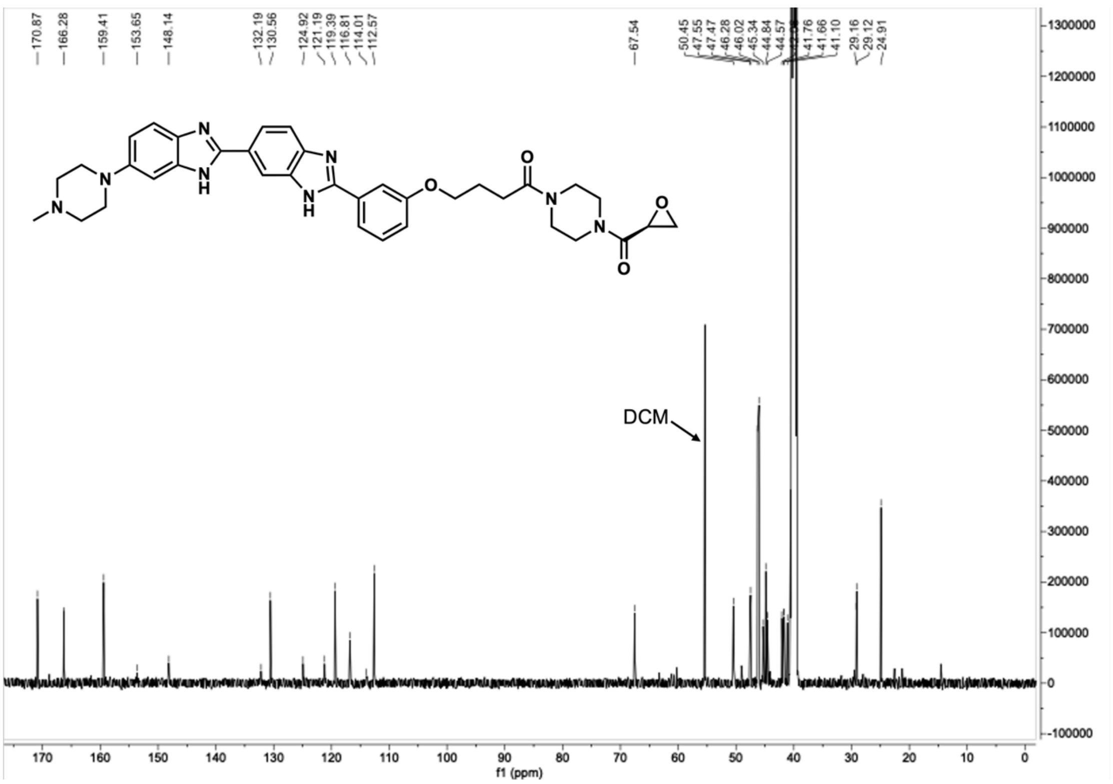

Compound **H6**, $^{1}H$ NMR, DMSO-$d_6$, 600 MHz.

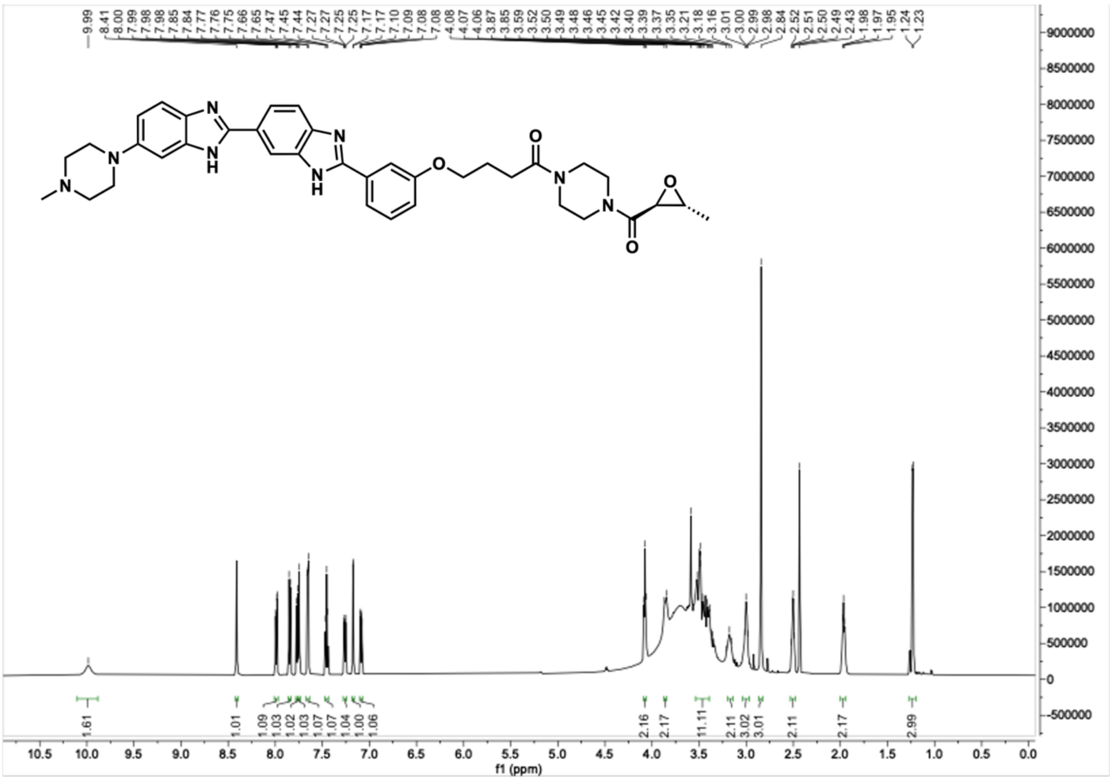

Compound **H6**, $^{13}C$ NMR, DMSO-$d_6$, 150 MHz.

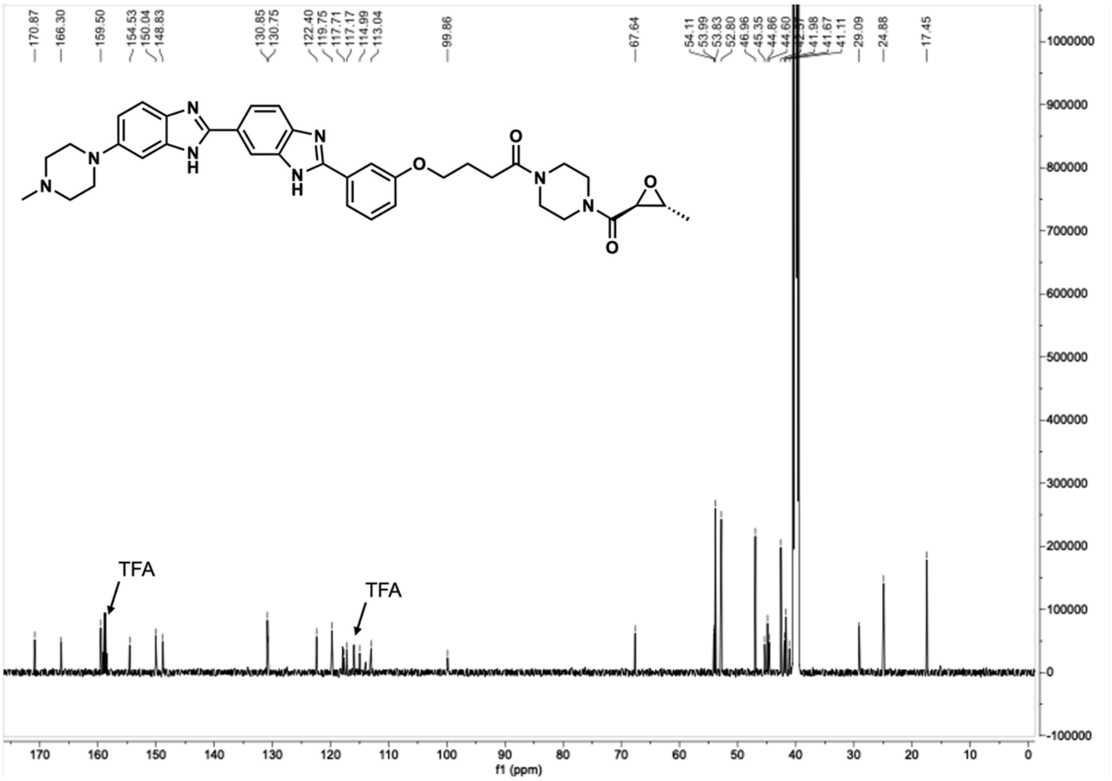

Compound **H7**, $^{1}$H NMR, DMSO-$d_6$, 600 MHz.

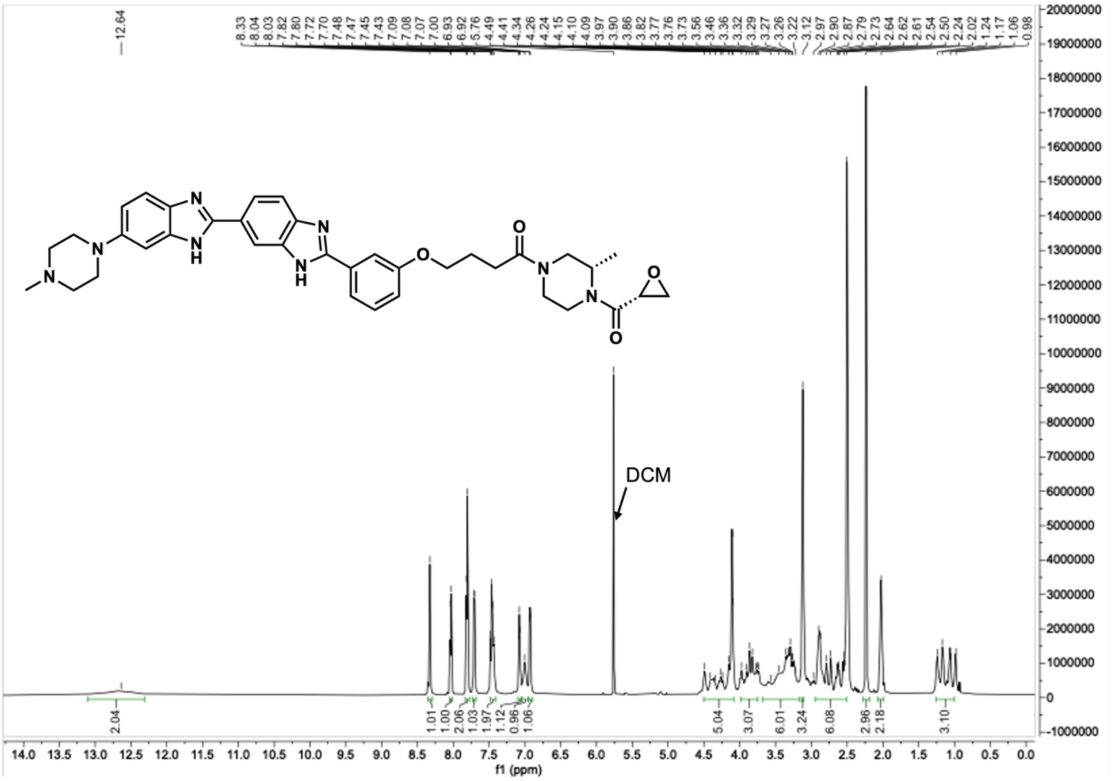


Compound **H7**, $^{13}$C NMR, DMSO-$d_6$, 150 MHz.

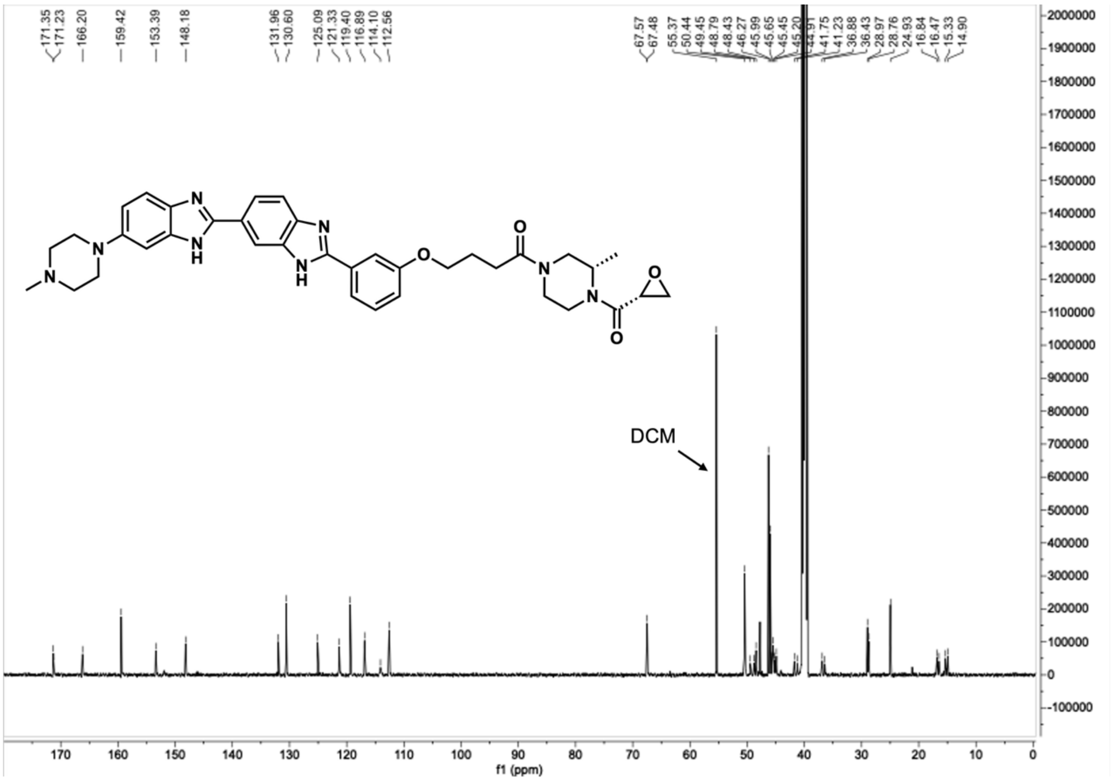

Compound **H8**, $^{1}$H NMR, DMSO-$d_6$, 600 MHz.

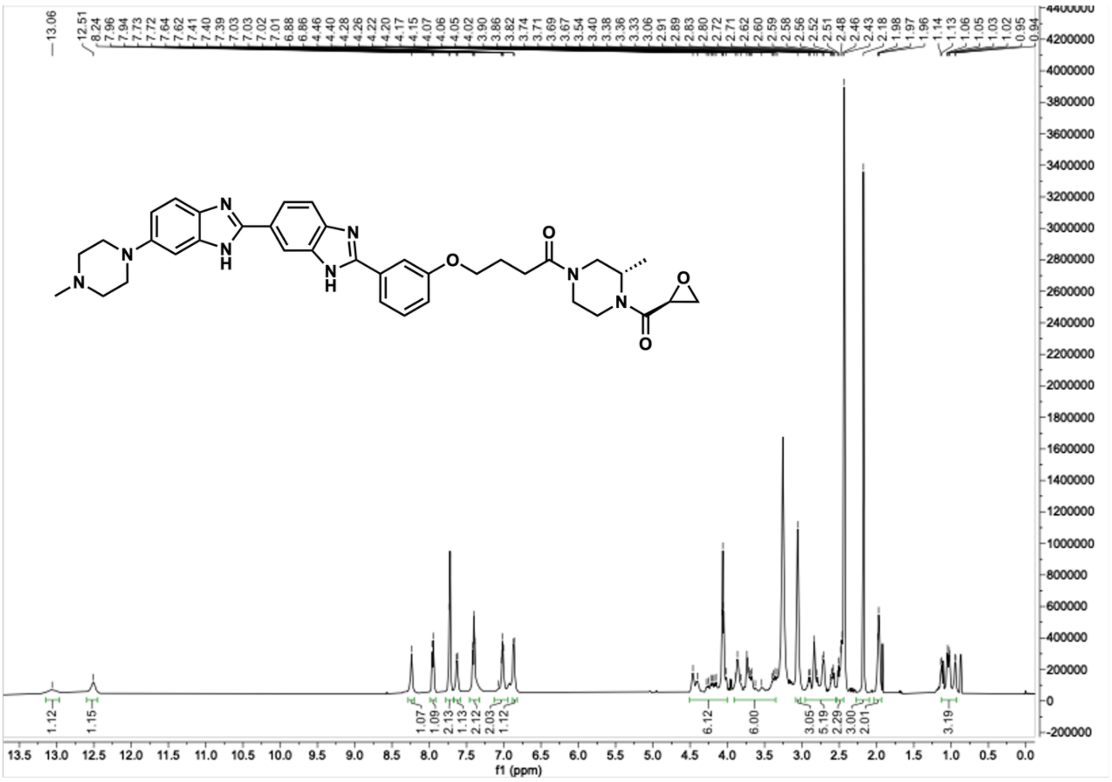

Compound **H8**, $^{13}$C NMR, DMSO-$d_6$, 150 MHz.

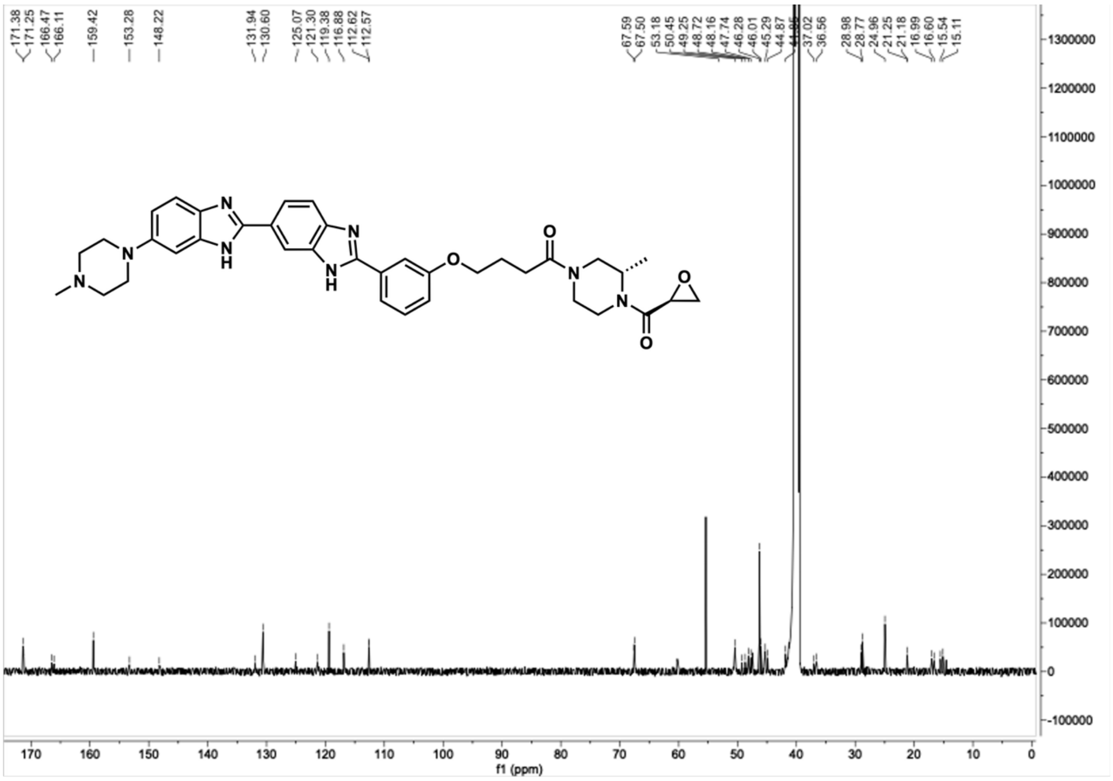

Compound **H9**, $^{1}H$ NMR, DMSO-$d_6$, 600 MHz.

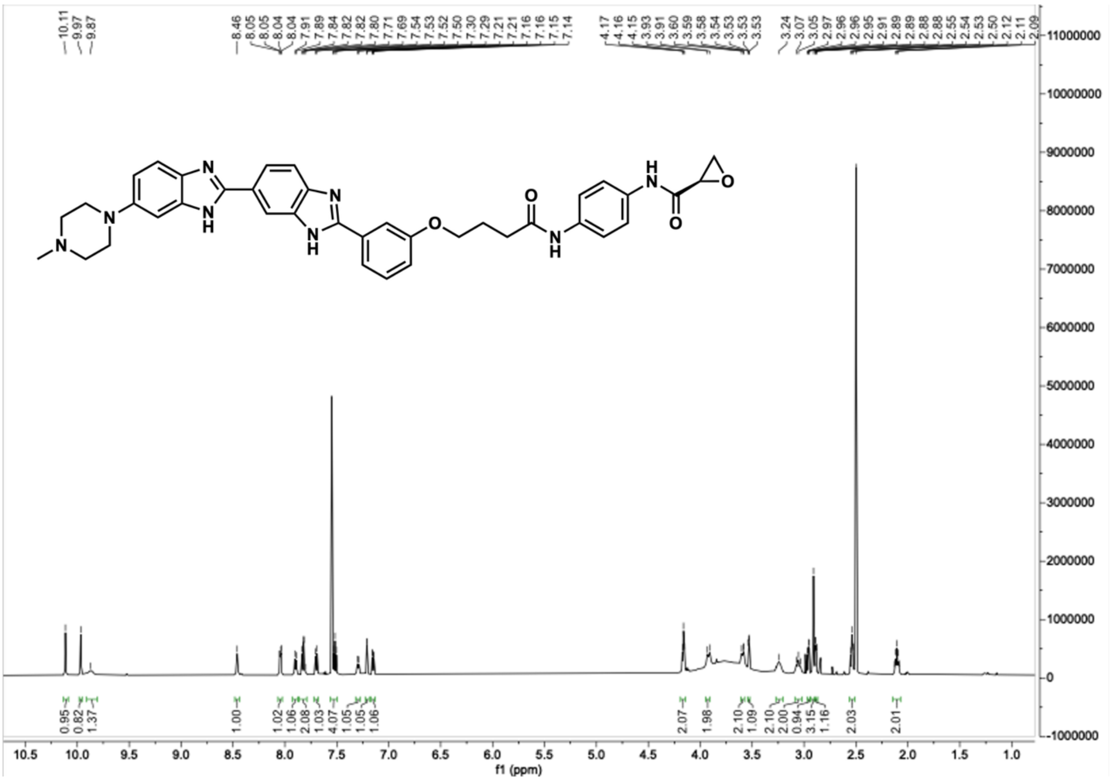

Compound **H9**, $^{13}C$ NMR, DMSO-$d_6$, 150 MHz.

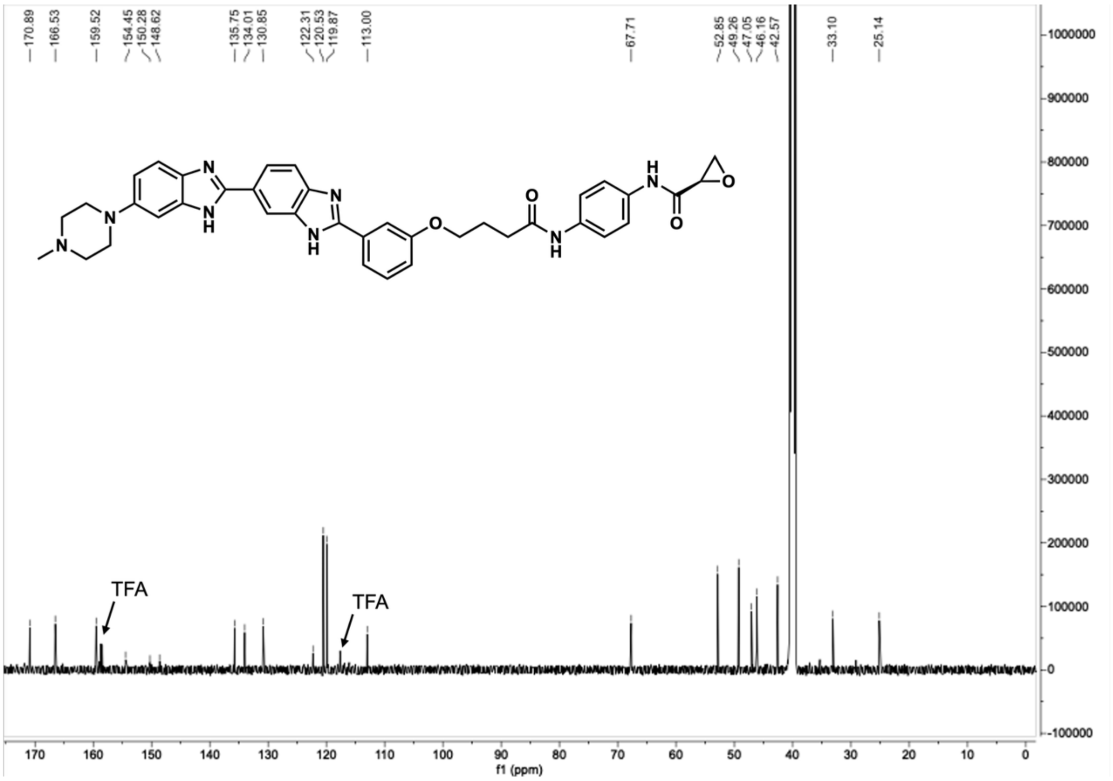

Compound **H10**, $^{1}H$ NMR, DMSO-$d_6$, 600 MHz.

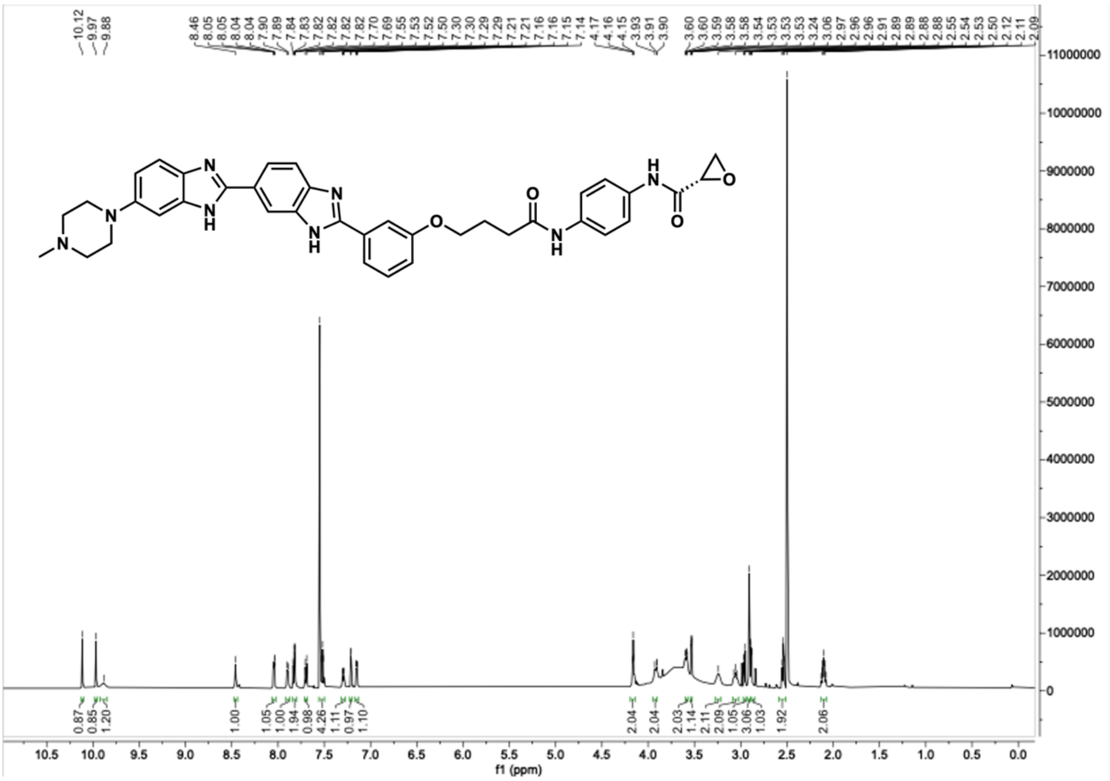


Compound **H10**, $^{13}C$ NMR, DMSO-$d_6$, 150 MHz.

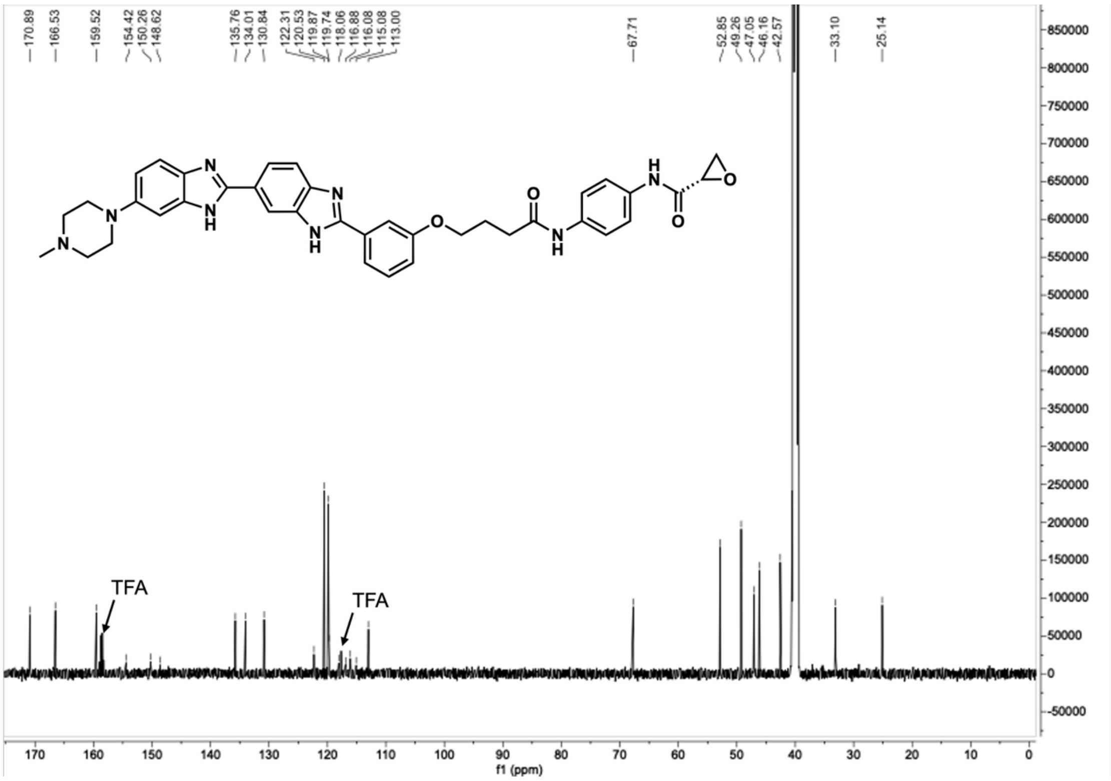

Compound **H11**, $^{1}H$ NMR, DMSO-$d_6$, 600 MHz.

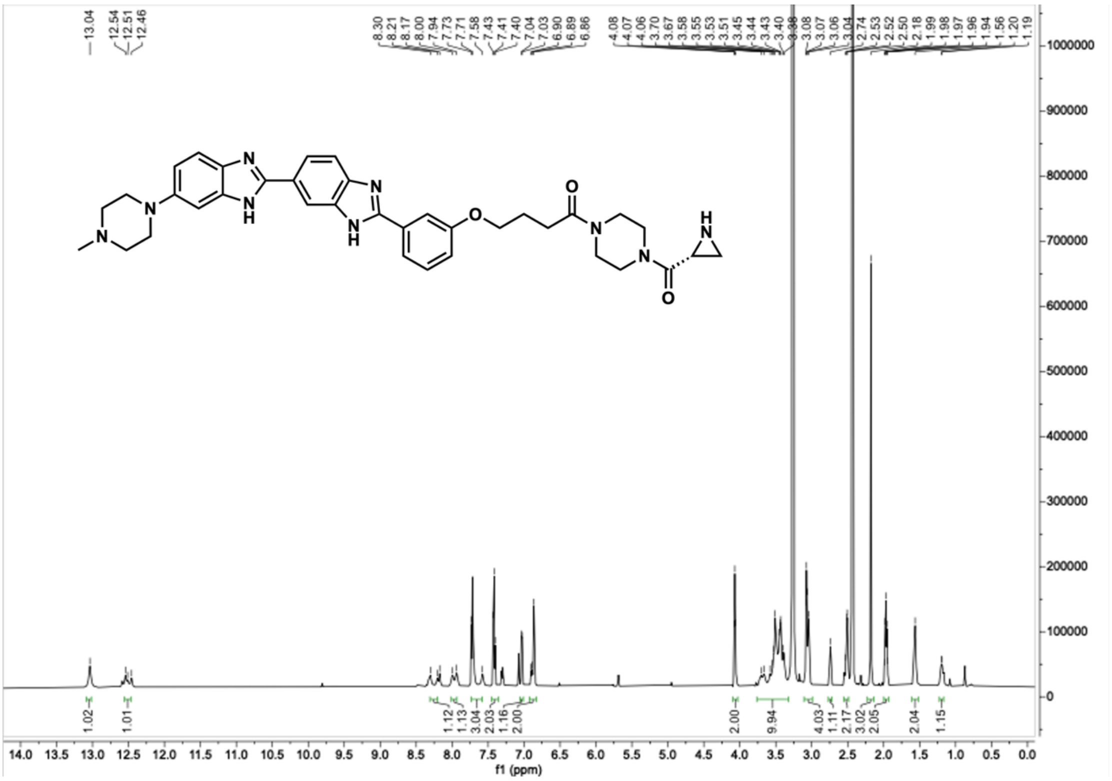

Compound **H11**, $^{13}C$ NMR, DMSO-$d_6$, 150 MHz.

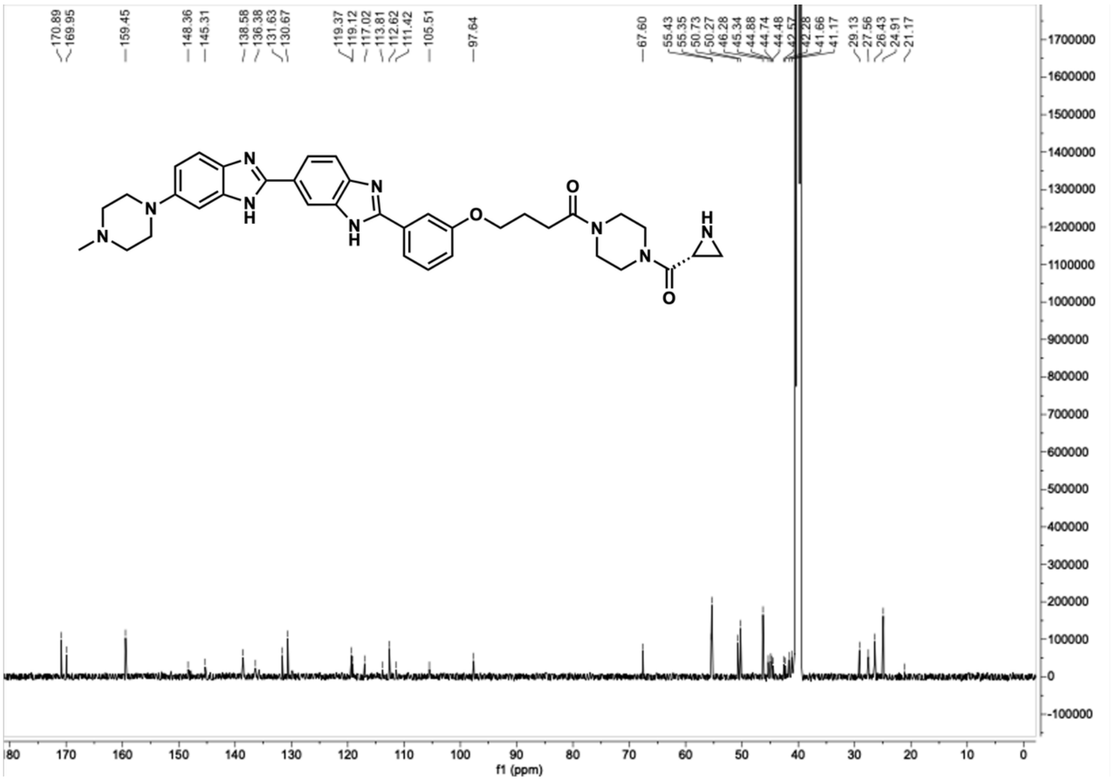

Compound **H12**, $^{1}$H NMR, DMSO-$d_6$, 600 MHz.

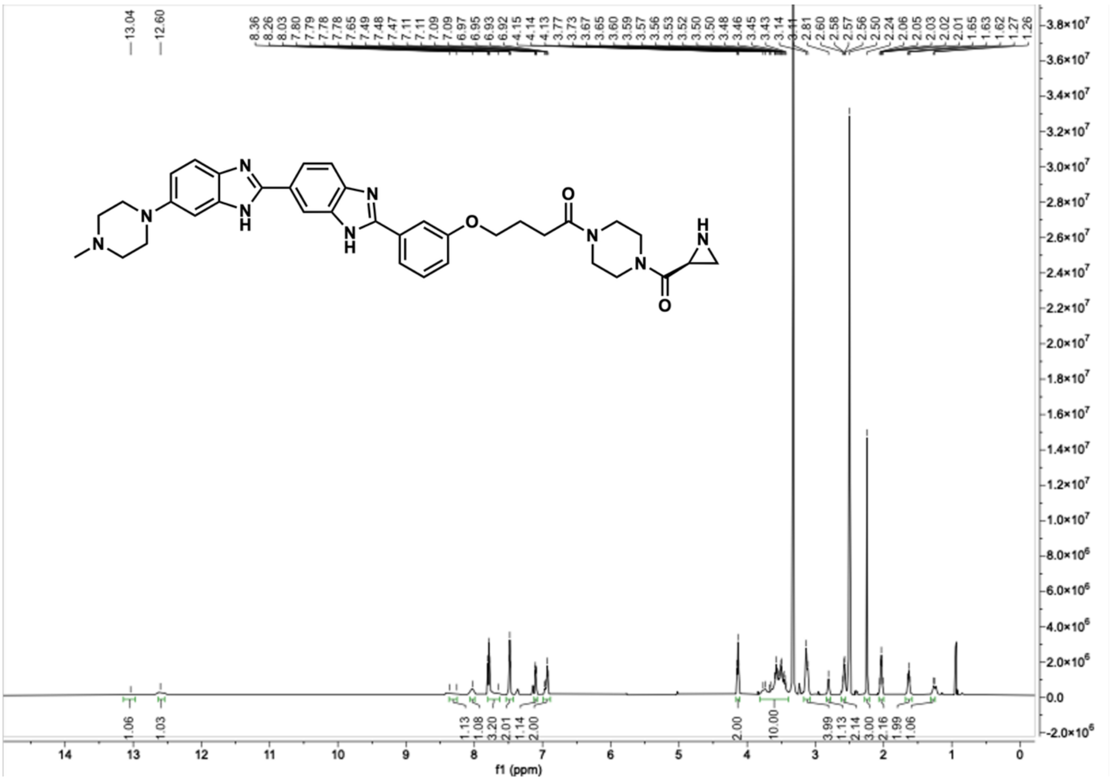


Compound **H12**, $^{13}$C NMR, DMSO-$d_6$, 150 MHz.

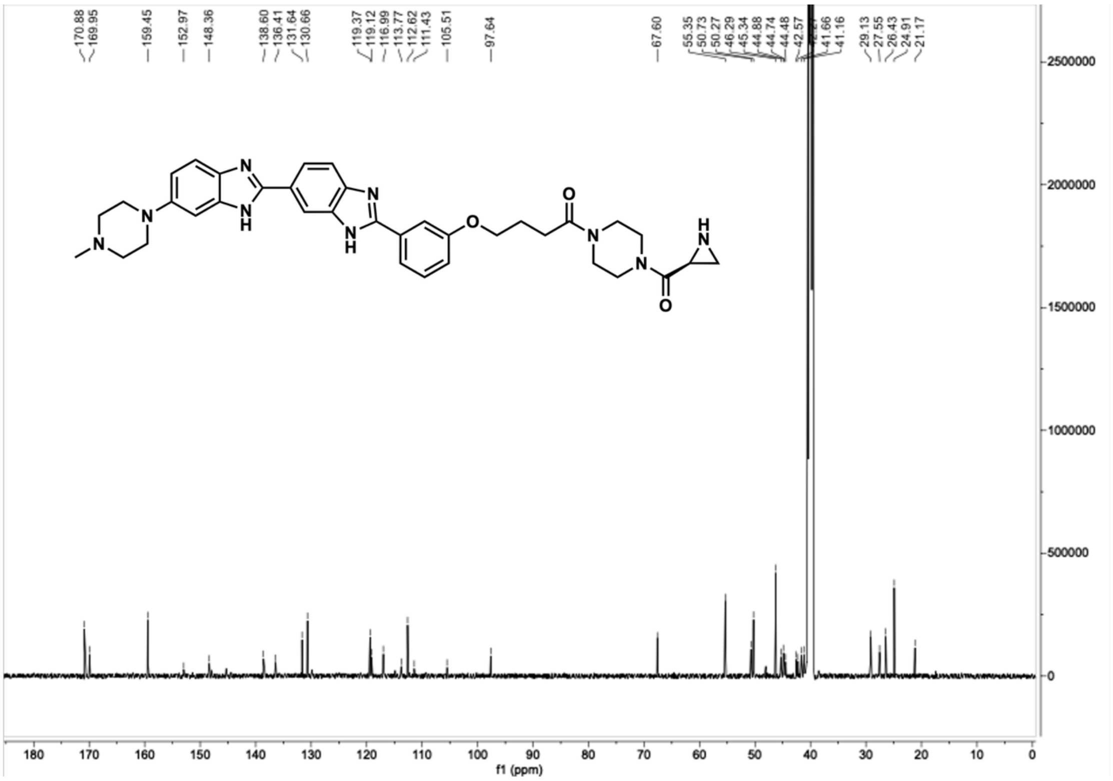

Compound **H13**, $^{1}H$ NMR, DMSO-$d_6$, 600 MHz.

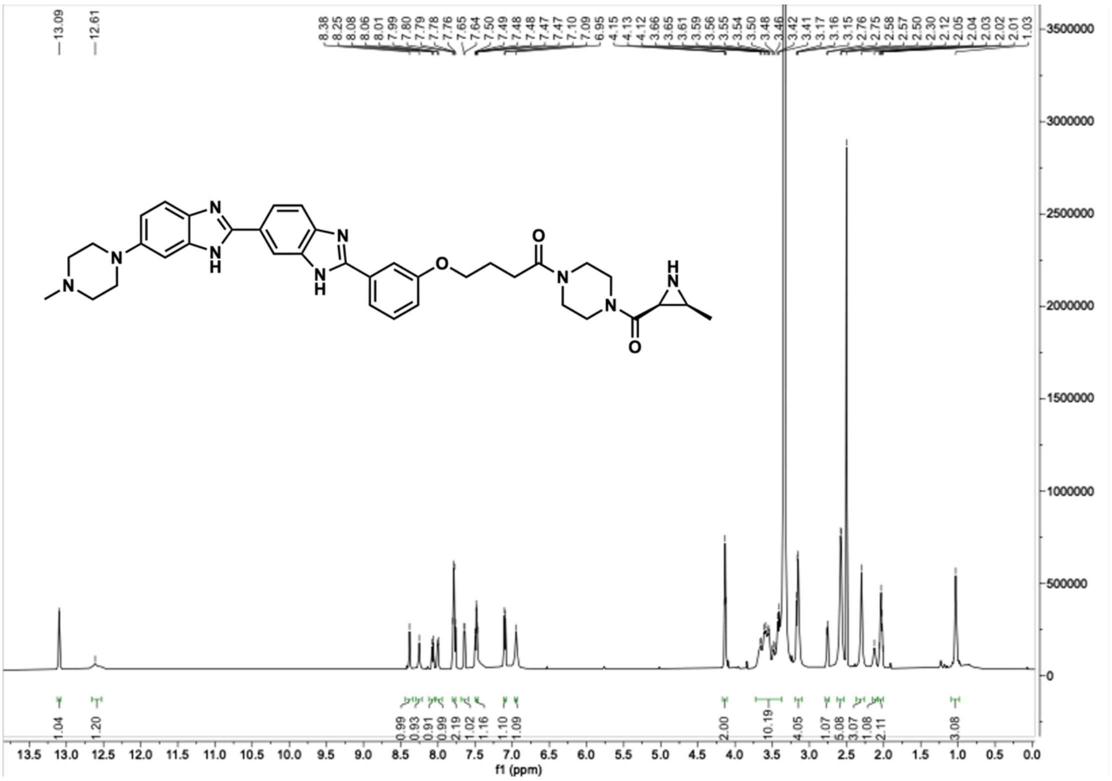


Compound **H13**, $^{13}C$ NMR, DMSO-$d_6$, 150 MHz.

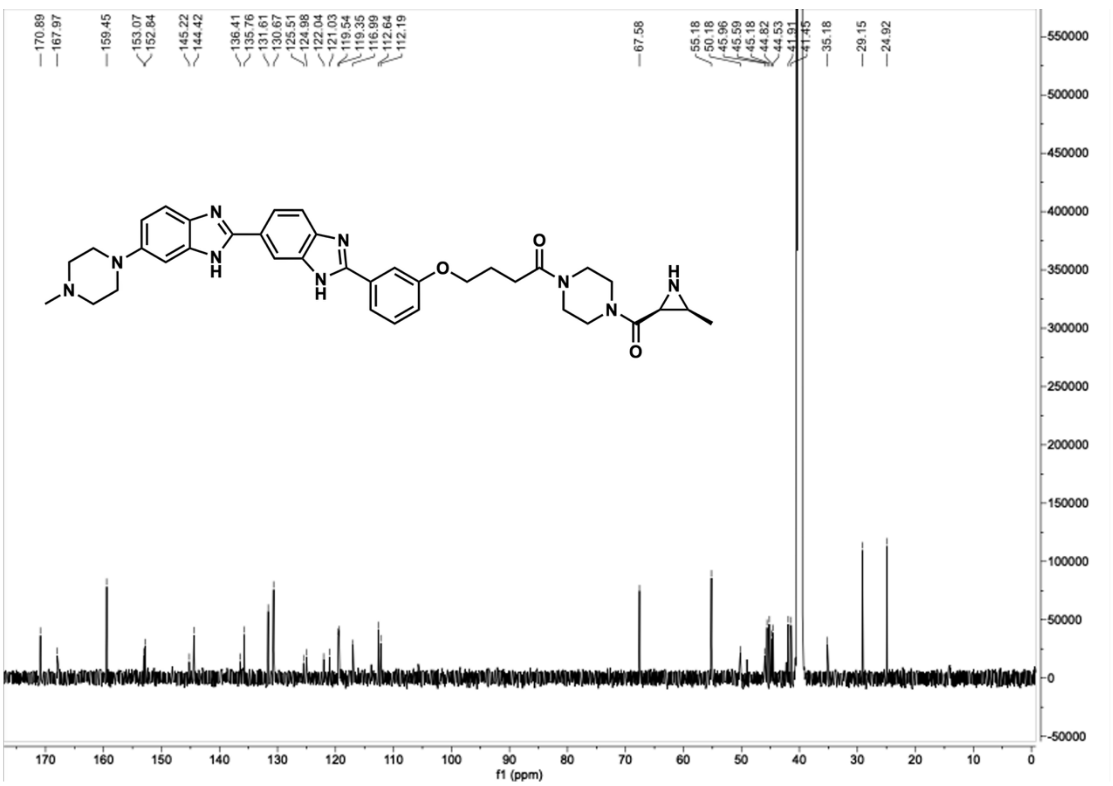

Compound **H14**, $^{1}H$ NMR, DMSO-$d_6$, 600 MHz.

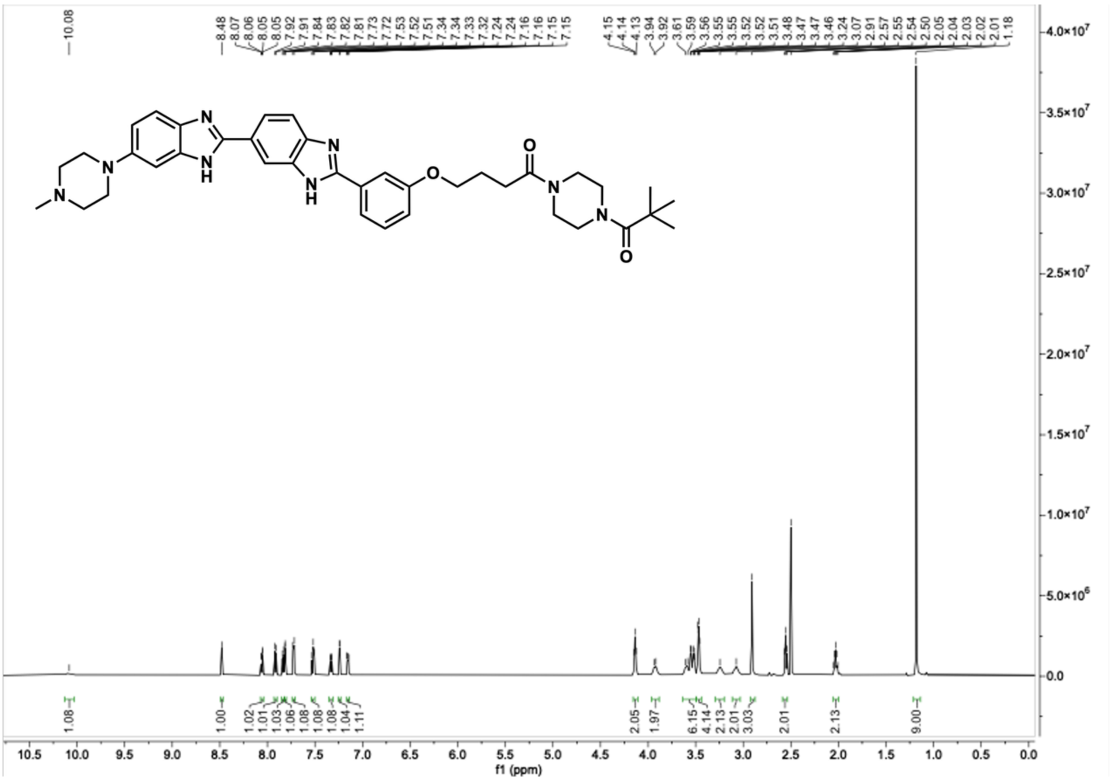

Compound **H14**, $^{13}C$ NMR, DMSO-$d_6$, 150 MHz.

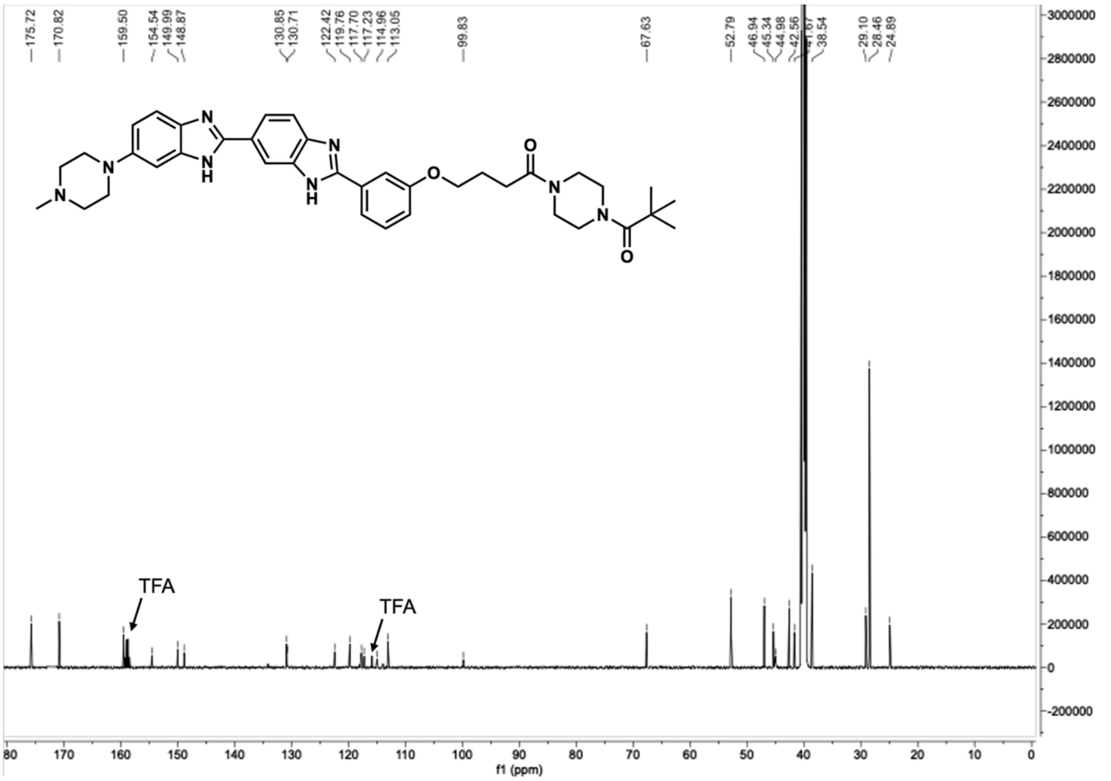

Compound **C1**, $^{1}H$ NMR, DMSO-$d_6$, 600 MHz.

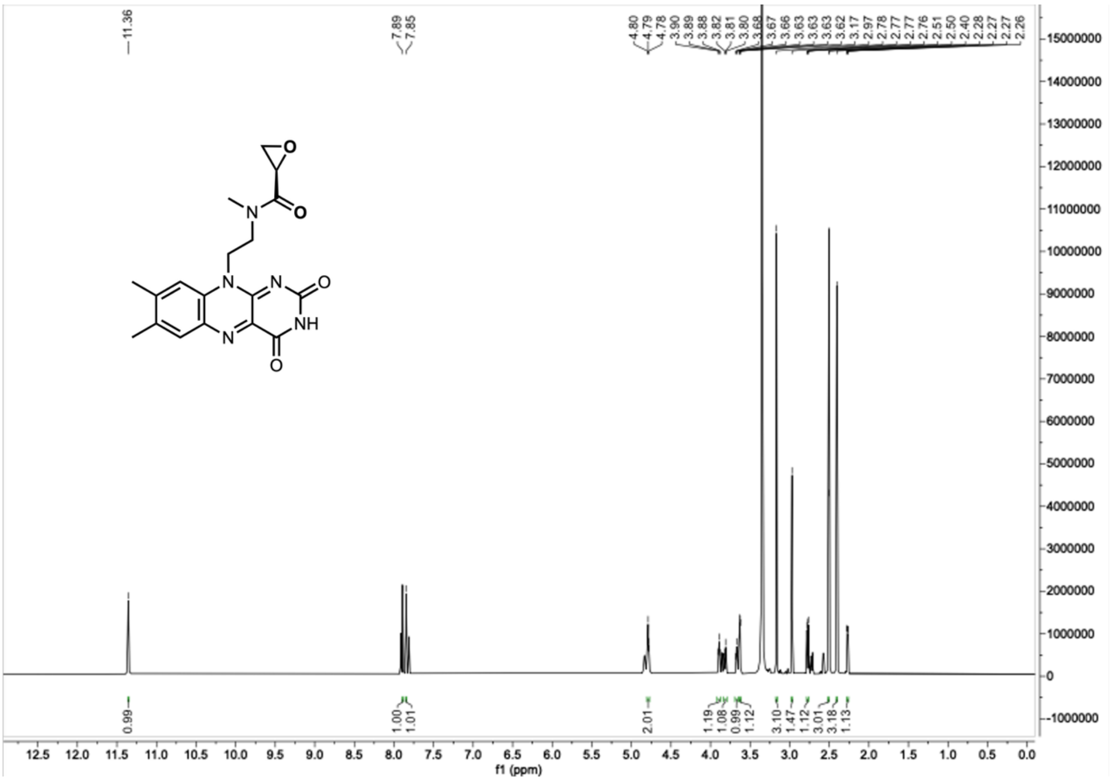

Compound **C1**, $^{13}C$ NMR, DMSO-$d_6$, 150 MHz.

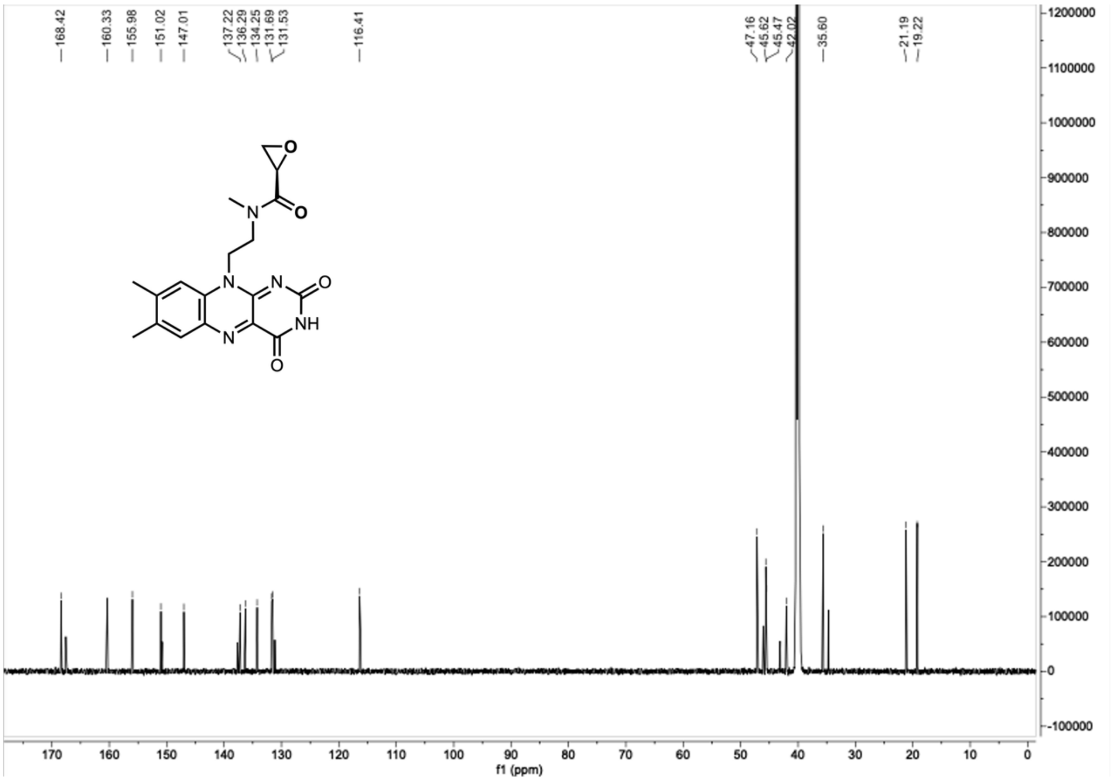

Compound **C2**, [1]H NMR, DMSO-$d_6$, 600 MHz.

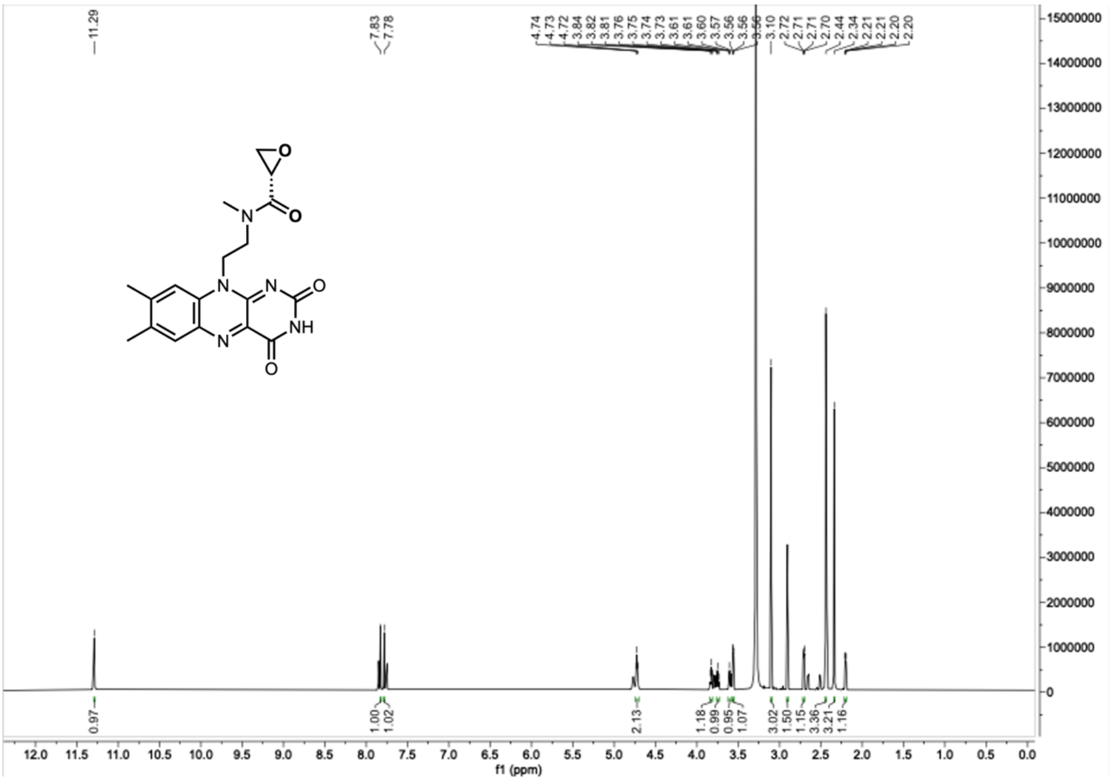


Compound **C2**, [13]C NMR, DMSO-$d_6$, 150 MHz.

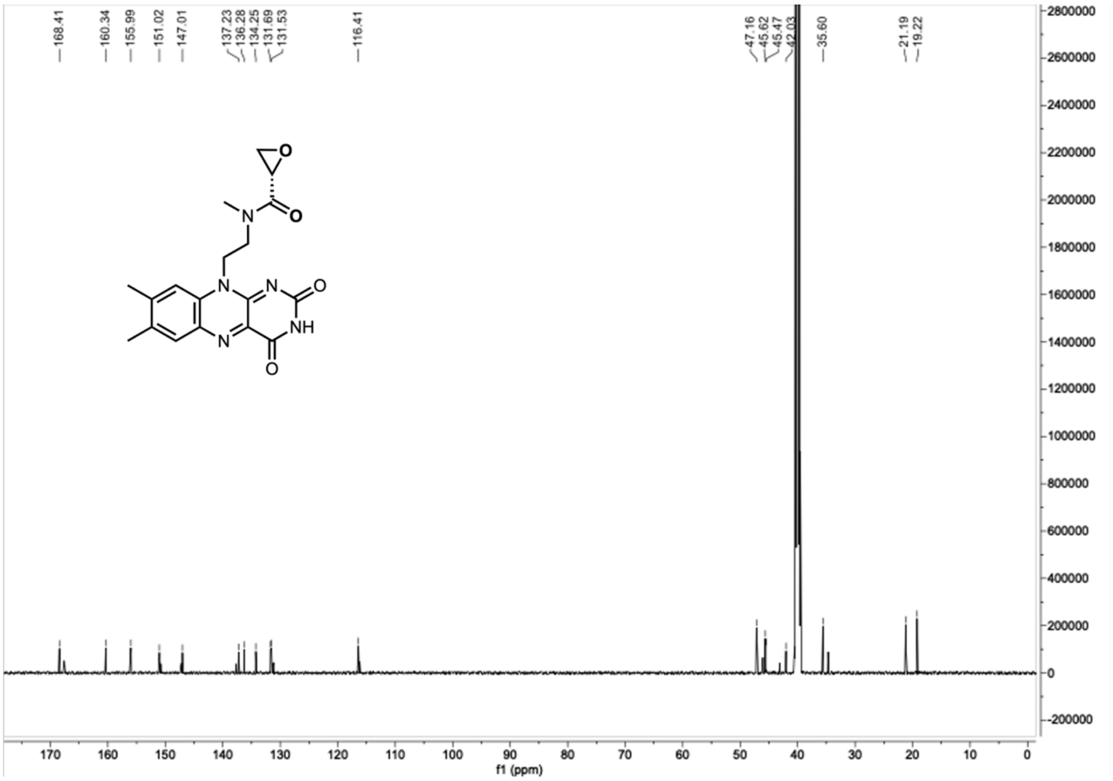

Compound **C3**, $^{1}H$ NMR, DMSO-$d_6$, 600 MHz.

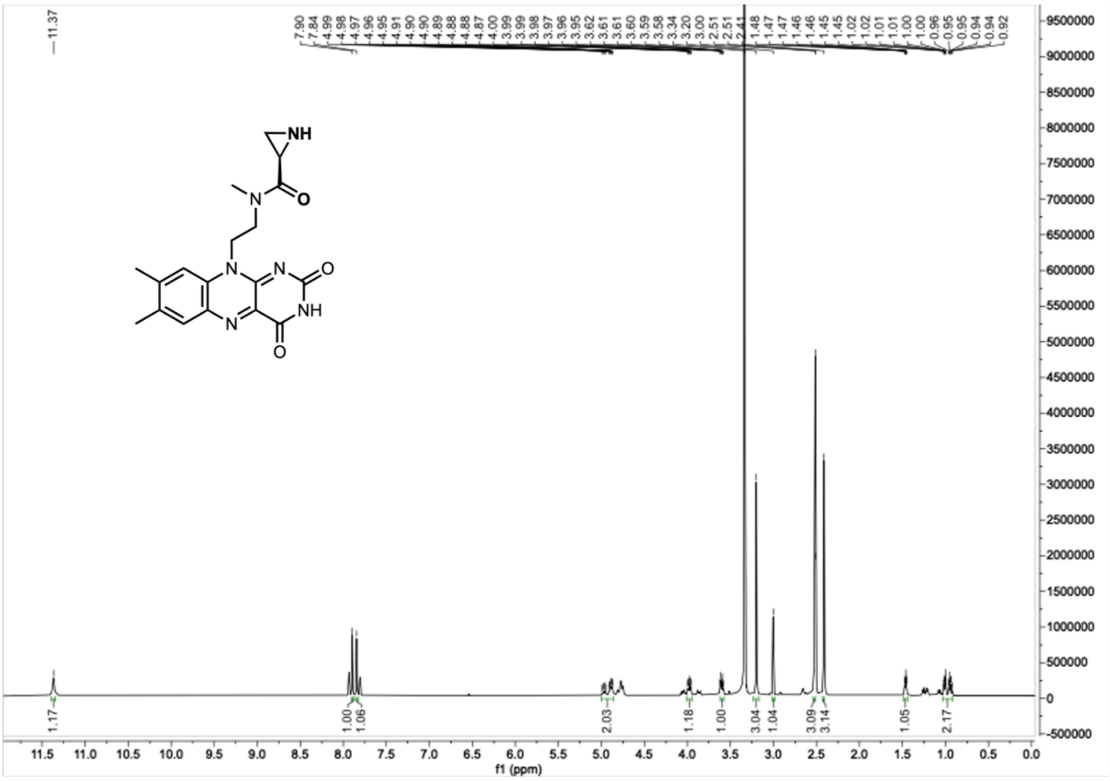


Compound **C3**, $^{13}C$ NMR, DMSO-$d_6$, 150 MHz.

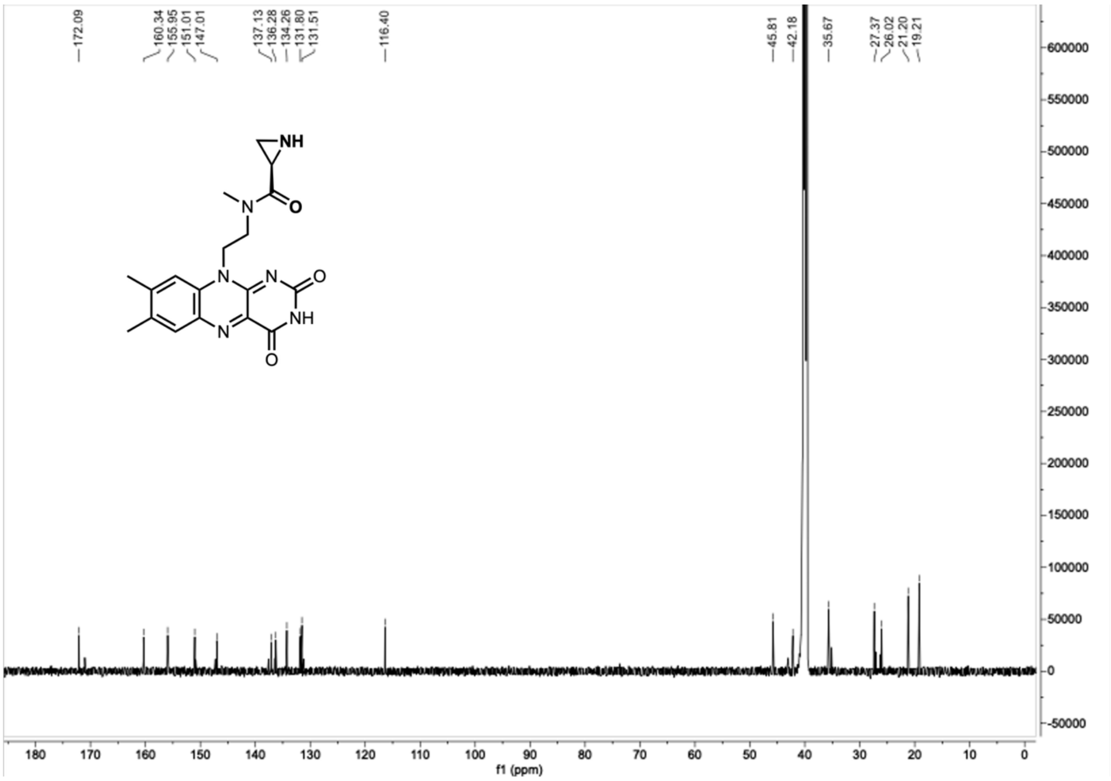

Compound **C4**, $^{1}$H NMR, DMSO-$d_6$, 600 MHz.

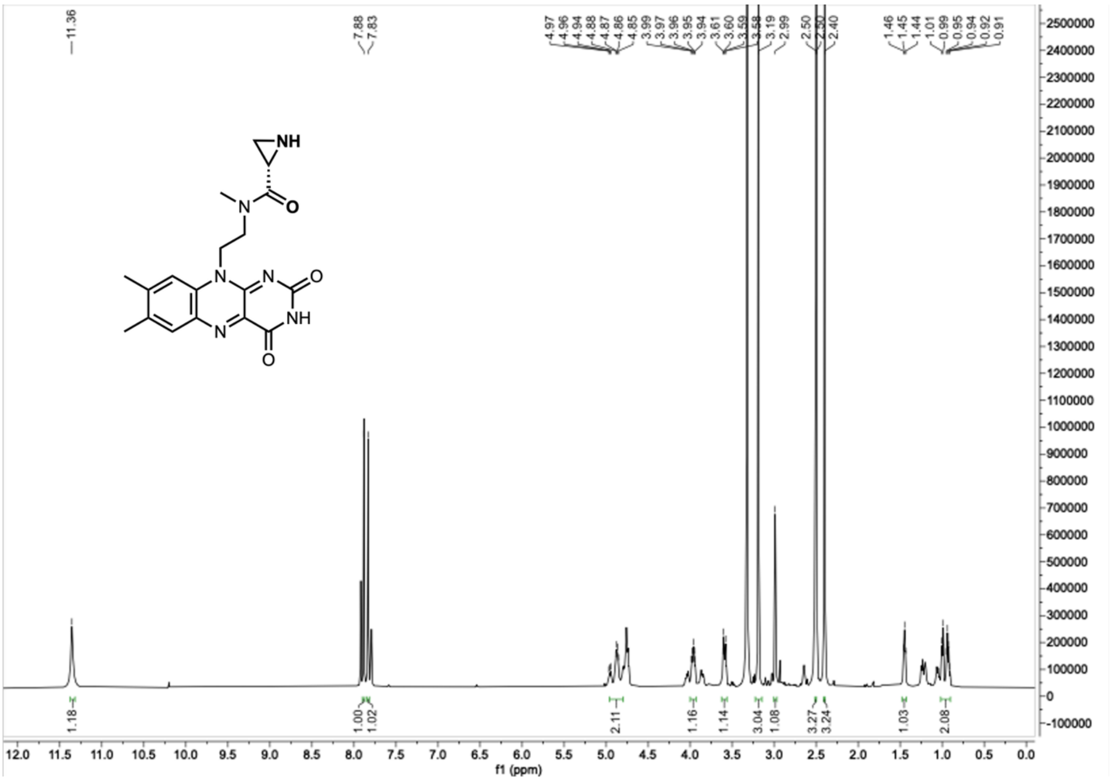


Compound **C4**, $^{13}$C NMR, DMSO-$d_6$, 150 MHz.

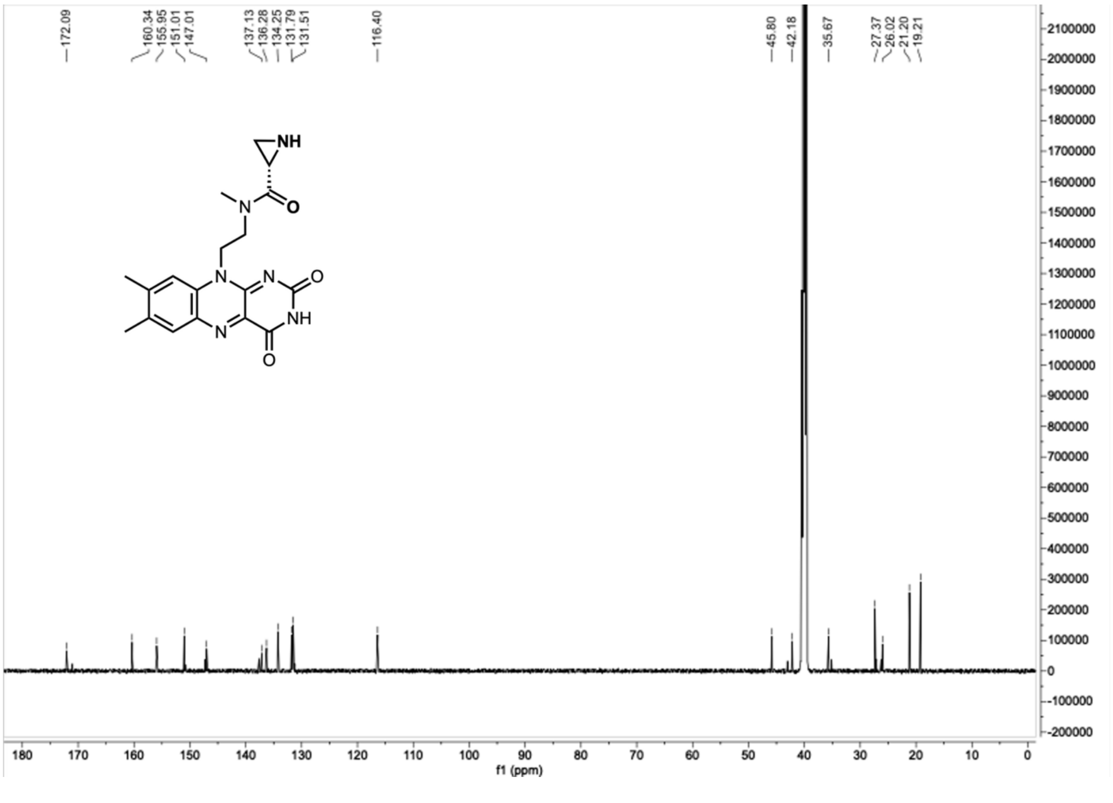
